# The spatial landscape of clonal somatic mutations in benign and malignant tissue

**DOI:** 10.1101/2021.07.12.452018

**Authors:** Andrew Erickson, Emelie Berglund, Mengxiao He, Maja Marklund, Reza Mirzazadeh, Niklas Schultz, Ludvig Bergenstråhle, Linda Kvastad, Alma Andersson, Joseph Bergenstråhle, Ludvig Larsson, Alia Shamikh, Elisa Basmaci, Teresita Diaz De Ståhl, Timothy Rajakumar, Kim Thrane, Andrew L Ji, Paul A Khavari, Firaz Tarish, Anna Tanoglidi, Jonas Maaskola, Richard Colling, Tuomas Mirtti, Freddie C Hamdy, Dan J Woodcock, Thomas Helleday, Ian G. Mills, Alastair D Lamb, Joakim Lundeberg

## Abstract

Defining the transition from benign to malignant tissue is fundamental to improve early diagnosis of cancer. Here, we provide an unsupervised approach to study spatial genome integrity in situ to gain molecular insight into clonal relationships. We employed spatially resolved transcriptomics to infer spatial copy number variations in >120 000 regions across multiple organs, in benign and malignant tissues. We demonstrate that genome-wide copy number variation reveals distinct clonal patterns within tumours and in nearby benign tissue. Our results suggest a model for how genomic instability arises in histologically benign tissue that may represent early events in cancer evolution. We highlight the power of an unsupervised approach to capture the molecular and spatial continuums in a tissue context and challenge the rationale for treatment paradigms, including focal therapy.

## Main Text

Mutations can either be inherited or acquired (somatic). Inherited genomic alterations are easy to identify as these are present in all cells while somatic mutations are usually present in a fraction of cells. Mutations can occur *de novo* in germ cells, estimated to 2-10 events per cell division^1^ or arise during development and adulthood as a result of DNA repair mechanisms as well as various environmental cues. For example, it has recently been demonstrated that somatic mutations in early development contribute to the genetic differences that exist between monozygotic twins^2^. The frequency and spatial distribution of these mutations has an important impact on phenotype. In order to obtain spatial information of clonal genetic events, studies have used laser capture microdissection of small regions or even single cells. These studies have an inherent bias as only a limited number of spatial regions per tissue section can be retrieved and examined. Furthermore, because investigators have selected such regions based on morphology, previous studies have limited their analyses to histologically defined tumour areas while excluding regions populated by benign cells. The possibility to perform unsupervised genome and tissue-wide analysis would therefore provide an important contribution to delineate clonal events.

Spatially resolved transcriptomics has emerged as a genome-wide methodology to explore tissue architecture in an unsupervised manner^3^. In this study we explore a method to infer genome-wide copy-number variations (CNV) from spatially resolved mRNA profiles in situ (**Fig. 1A**). Gene expression has previously been used to inferCNVs in single cells, successfully identifying regions of chromosomal (chr) gain and loss^4^. Here we expand into a spatial modality generating genome-wide CNV calls in each spatial region represented by barcoded spots (Supplementary Material). First we sought corroboration that transcript-derived inferCNV (iCNV) data could accurately mirror DNA-based phylogenies, using simultaneously extracted single cell RNA, and DNA^5^ (fig. S1-2). Next we successfully recapitulated published DNA-based phylogenies in prostate cancer using RNA from the same samples^6–8^ (fig. S3). To ensure that we robustly could capture sufficient and accurate CNV information from approximately 10-20 cells per spot and use this information to deduce clonal relationships between cells, we then designed an *in-silico* system to synthesize a tissue containing multiple clones determined by stochastic copy number (CN) mutations in a single artificial chromosome. Using a probabilistic method to generate gene expression from such mutations we then interrogated the expression data using iCNV, while blind to the underlying ‘ground-truth’ CN status, and successfully recapitulated both the CN status and the clonal groupings (Supplementary material; fig. S4).

**Fig. 1.**
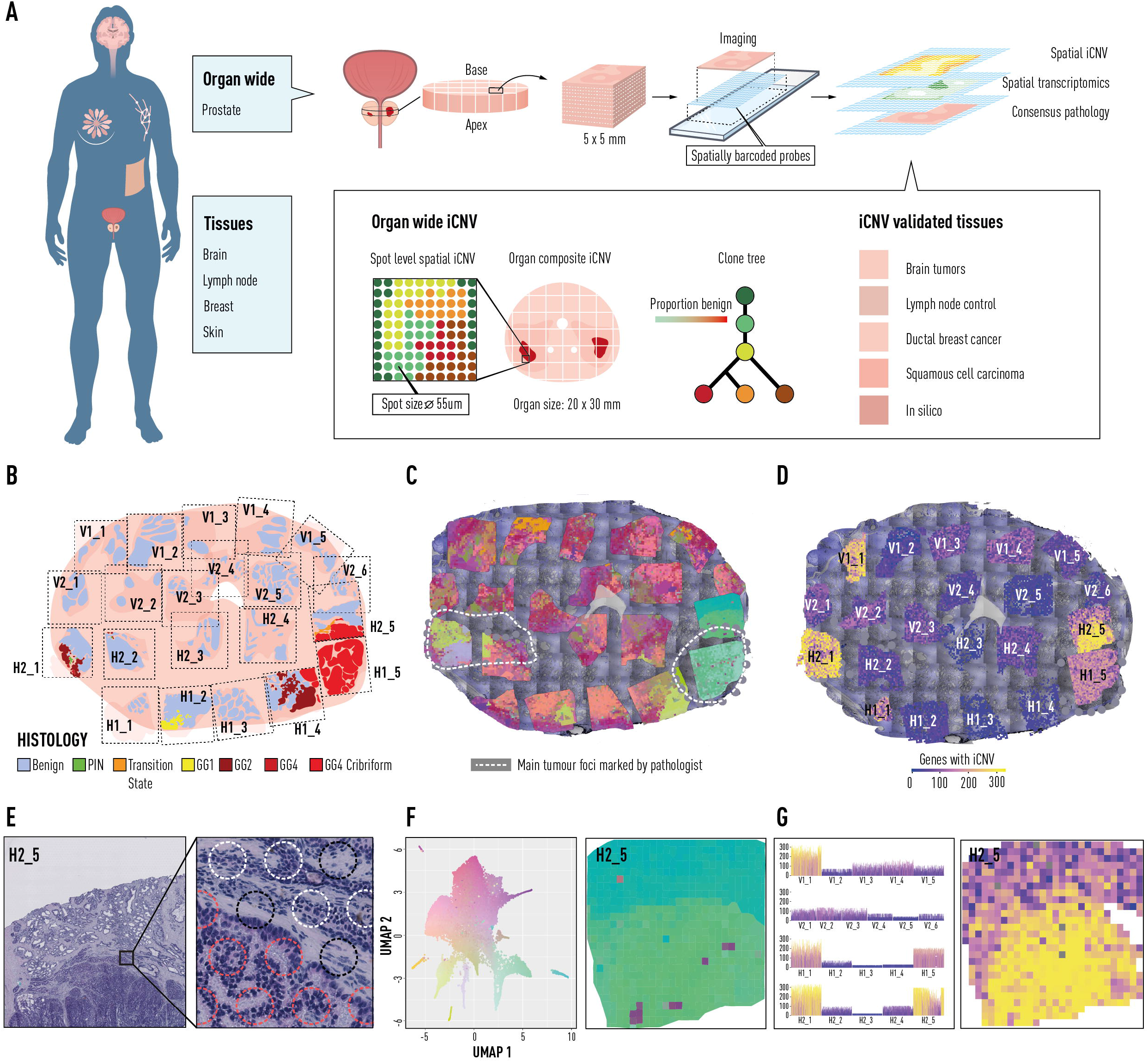
Organ-wide spatial determination of transcript and CNV status. **(A)** For organ-wide assessment axial segments of the prostate were divided into 5×5mm blocks for spatial transcriptomic analysis with spatially barcoded probes. The resulting spatial gene expression profile was accompanied by inferred copy number profile supported by spot-by-spot consensus pathology calls. Copy number features were used to detect clonal groups and instruct phylogenetic tree construction. Tissue specific analyses of multiple phenotypes were performed. (**B)** Histology status for each organ-wide section. Black dotted lines represent the area covered by spatial transcriptomics array surface. (**C)** Spatial distribution of gene expression (see panel F). (**D)** Spatial distribution of summed copy number events (see panel G). (**E)** Representative spot-level consensus pathology for Section H2_5. Red circles = >50% cancer. Blue circles = >50% benign epithelium. Black circles = <50% of a single cell-type. (**F)** UMAP principal component analysis of gene expression factors with representative close-up for Section H2_5. (**G)** Total copy number events for each section with representative close-up for Section H2_5.

Next we used prostate cancer to explore the spatial iCNV landscape of a commonly multifocal malignancy^9^. The specimen was obtained by open radical prostatectomy and an axial section was taken from the mid-gland (fig. S5). The axial section was subdivided into cubes (**Fig. 1A, B**) and corresponding tissue sections were histologically graded using the Gleason grading system^10^ identifying extensive intratumoural heterogeneity (ITH) in the context of benign tissue (**Fig. 1B, E**; fig. S6, Supplementary Material). We obtained transcriptional information from 21 cubes (tissue sections) and > 21 000 barcoded regions (100 micron spots) with a mean average of 3500 genes detected per barcoded spot, from the cross section of the entire prostate^11^ (fig. S7). Next, we analysed the barcoded gene expression data using factorized negative binomial regression (fig. S8). This provided an unsupervised view of gene expression factors (GEFs) over the cross section of the prostate (**Fig. 1C**). Twenty-five gene expression factors showed overlap between histology and GEFs representing tumour, hyperplasia and benign epithelia annotated by the factor marker genes, as previously reported^12^ (**Fig. 1F**, fig. S9, table S1). Several GEFs provided distinct ‘clonal’ appearances and were associated with tumour regions (**Fig. 1F, right panel**). We then undertook a spatial iCNV analysis to provide an overall landscape of genome integrity (**Fig. 1D**) identifying certain regions with increased iCNV activity (V1_1, H2_1, H1_1, H1_5, H2_5; **Fig. 1G**) while the majority of the tissue area appears to be CN neutral. This suggested that iCNVs could identify tissue regions of interest, distinct from morphology or expression analysis.

To increase the precision in our analysis of these iCNV regions we took advantage of smaller 55 micron diameter barcoded spots (Visium, 10x Genomics), reducing the number of cells to approximately 5-10 per spot, to perform a more detailed interrogation of seven key sections. We first validated the increased precision of this higher resolution platform using the synthetic tissue method (fig. S10). We next obtained data from approximately 30 000 spots using factorized negative binomial regression resulting in 24 spatially distinct GEFs (fig S7, S11, S12, table S2). Two pathologists independently annotated each spot to provide consensus pathology scoring (**Fig. 1E**, fig. S13). We then investigated clonal relationships across the prostate using iCNVs. Having established the association between gene expression factors and certain regions of interest (**Fig. 1C, F**) we wanted to determine the degree of clonal CN heterogeneity in these regions. After designating all histologically benign spots as a reference set (fig. S14A) it was immediately apparent that while certain GEFs displayed a fairly homogenous inferred genotype (e.g. GEF 7, 14 and 22, fig. S14B), others were strikingly heterogeneous (e.g. GEF 10, fig. S14C).

Prompted by the realization that certain regions annotated as histologically benign displayed CN heterogeneity (**Fig. 1D**, fig. S15), we refined the reference set to those spots which were both histologically benign (outside the regions of interest) and also lacking any iCNV (Supplementary Material, fig. S16). This constituted a ‘pure benign’ reference set for all subsequent iCNV analyses, unique to each patient. It was apparent from cancer-wide inferred genotype (**Fig. 2A-E**) that there were distinct spot groupings with evidence of CN heterogeneity within areas of spatially homogeneous Gleason pattern (**Fig. 2A, D, E**). We constructed a clone-tree to describe sequential clonal events versus independently arising cancer-clones (**Fig. 2B**). It was apparent that two cancer clones lacked key truncal events including a loss of a region of chr 16q and 8p which were otherwise ubiquitous across all cancer clones (clones A and B, **Fig. 2A, B**). These clones were spatially restricted to section H1_2 containing a region of low-grade Gleason Grade Group 1, discussed later. The majority of clonally related spots were spatially located around the largest focus of Gleason Grade Group 4 disease with a striking pattern of truncal and branching events (clones H, I, J and K). We therefore focused on this dominant region of cancer (spanning sections H1_4, H1_5 and H2_5), to first establish spatial and clonal dynamics in this one area (remaining sections in fig. S17).

**Fig. 2.**
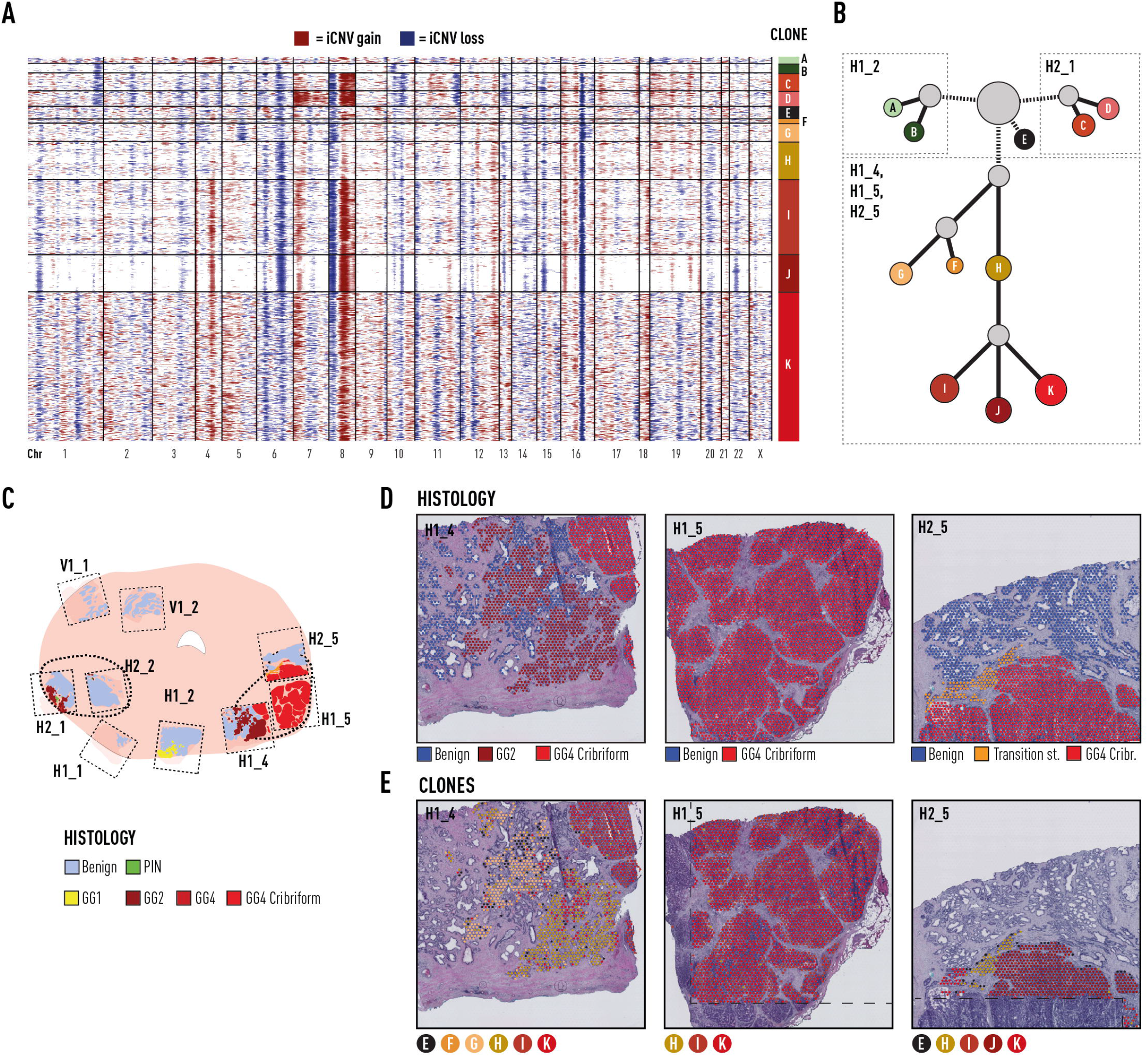
Specific somatic alterations in all cancer organ-wide analysis. (**A)** Genome-wide derived analysis (inferCNV) for all Visium spots harbouring tumour from prostate patient 1. Clonal groupings of spots (approx. 10-15 cells each) determined by hierarchical clustering (Supplementary Methods). (**B)** Phylogenetic clone tree (Supplementary Material) of tumour clones (from panel A), with grey clones representing unobserved, inferred common ancestors. Clone circle area is proportional to number of spots and branch length determined by weighted quantity of CNVs, both on a logarithmic scale (Supplementary Material). **C**. Representation of all tissue sections from prostate patient 1. Thicker black lines denote original boundaries annotated by initial clinical pathology. **D**. Consensus epithelial histological annotations for sections H1_4, H1_5 and H2_5, corresponding to the right tumour focus. **E**. Spatial visualization of tumour clones (Panel A). Dashed line marks areas where no spatial transcriptomics data was obtained due to technical limits. Chr = chromosome. iCNV = inferred copy number variant. PIN = prostatic intra-epithelial neoplasia. GG = ISUP Gleason ‘Grade Group’

To construct clone-trees, we assumed that: (i) groups of cells containing identical CN profiles were more likely to be related, than to have arisen by chance; and (ii) somatic CN events must be acquired sequentially over time, the more numerous the events, the more distinct the clone. Using this approach, we observed a common ancestral clone (clone H, **Fig. 2B**) containing truncal events including CN loss on chr 6q and 16q, and CN gain on 12q and 16q. These were spatially located in two regions: an area of Gleason Grade Group 2 on the medial side of the main tumour focus (section H1_4) and a region described as ‘transition state’ by consensus pathology at the upper mid edge (section H2_5). These conserved features in distinct spatial locations are noteworthy. A possible explanation is that clone H represents a linear sequence of branching morphology in the prostatic glandular system^13^, and that further somatic events took place giving rise to clones I, J and K forming a high-grade tumour focus (**Fig. 2B**), which pushed apart the branching histology due to an aggressive expansile phenotype. For the first time we have a spatial imprint of these events in prostate tissue. We also propose that some CNVs may be of particular pathological significance based on spatial molecular phylogeny. Our analysis therefore provides insight into processes of tumour clonal evolution, identifying discriminating events by spot-level CNV calling in a spatial context.

Given the discordance between cellular phenotype and inferred genotype, we then undertook a detailed interrogation of section H2_1 in the left peripheral zone of the prostate (**Fig. 1C, 2C**) containing roughly equal proportions of cancer and benign tissue. We profiled CN status of every spot in this section and ordered these spots by hierarchical clustering into ‘clones’ A to G based on defined levels of cluster separation (**Fig. 3A, B**; Supplementary Material). Spatially, we observed that these data-driven ‘clone’ clusters were located in groups, broadly correlating with histological subtype, but with some important distinctions (**Fig. 3C, D**). We also observed that many CNVs already occurred in clone C (**Fig. 3A-D**), most notably in chr 8, which has been well-described in aggressive prostate cancer^14–16^, but also several other CN gains and losses. Spatially, this clone constituted a region of exclusively benign acinar cells branching off a duct lined by largely copy neutral cells in clone A and B (**Fig. 3D**). The unobserved ancestor to clone C then gave rise to a further unobserved clone, and then clones E, F and G. While clone G was made up exclusively of Gleason Grade Group 2 cancer cells, clones E and F were mixed cancer and benign (**Fig. 3D**) highlighting, for the first time, that these clone groups traverse histological boundaries.

**Fig. 3.**
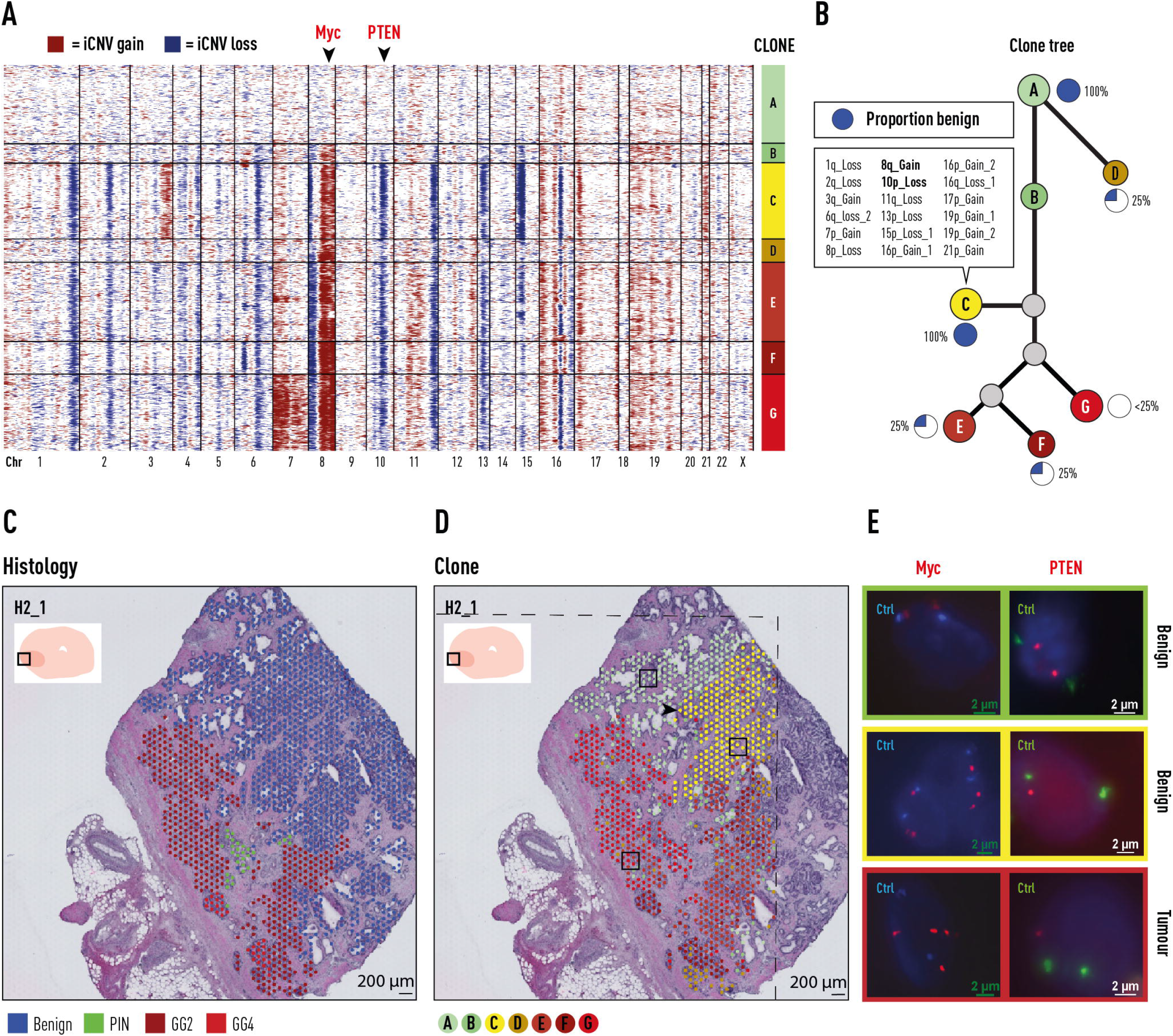
Somatic events in both cancer and benign prostate epithelium. **(A)** Genome-wide derived CNV analysis (iCNV) for each barcoded high-resolution spatial transcriptomic (ST) spot from Section H2_1, which contained a mixture of tumour and benign epithelia (red = gain; blue = loss). Clonal groupings of spots (approx. 10-15 cells each) determined by hierarchical clustering (Supplementary Material). **(B)** Phylogenetic clone tree (Supplementary Material) of all clones (from panel A). Proportion of benign epithelial cells in each clone as indicated. Specific CNV locations unique to Clone C are listed (summarized by chr number where event is located, p/q arm and gain/loss; remainder of iCNV changes in supplementary data S1). **(C)** Spatial visualization of histopathological status of each spot. Each spot assessed by two pathologists for consensus annotation with only spots >50% cellularity included (Supplementary Material). **(D)** Spatial visualization of clone status of each spot. Clonal groupings cross histological boundaries. Branching point of prostatic duct (black arrow), indicates possible site of somatic events arising in Clone C (also fig. S18). Dashed line marks areas where no spatial transcriptomics data was obtained due to technical limits. **(E)** FISH validation of two iCNVs: MYC, from chr8q; and PTEN, from chr10p (black arrow heads on panel A). Control probes (Ctrl) target centromeres for chr8 and chr10 respectively. Chr = chromosome. CNV = copy number variant. PIN = prostatic intra-epithelial neoplasia. GG = ISUP Gleason ‘Grade Group’

In order to validate that this inferred CN status was truly representative of underlying genotype we used fluorescence in situ hybridization (FISH) probes to target two specific genes of discriminatory interest, *MYC* and *PTEN*, encompassed in the notable chromosomal changes in benign tissue clone C as well as high grade tumour clones, but absent in low grade disease. This confirmed that while the status of both genes was diploid in normal benign tissue (Clone A), *MYC* amplification and *PTEN* loss were evident in altered benign (Clone C) as well as in tumour clones (Clone F; **Fig. 3E**, fig. S18-19). This evidence suggests that somatic events, creating a mosaic of branching clones during ductal morphogenesis, are present even in histologically benign disease. It therefore follows that an understanding of this somatic mosaicism could distinguish which regions of benign glandular tissue may give rise to lethal cancer, and which will not.

We considered the place of branching morphogenesis in the sequential acquisition of transformative events in a predominantly benign section of the prostate (section H2_1, also in section H2_2) (fig. S20). Here we noted that such events seem to occur during the development of prostatic ducts and acinar branches, with changes occurring at key branching points (marked by X; fig. S20), and the altered genotype passed on to daughter cells lining the ducts and glands of associated branches. Interestingly, not all cells in such branches displayed the same phenotype, raising important questions as to why epithelial glands with seemingly identical inferred genotypes might display divergent histological phenotypes.

In view of the above findings, we hypothesized that analysis of the inferred genotype of low grade cancer might reveal important differences to that of high grade cancer. Section H1_2 contains a region of Gleason Grade Group 1 prostate cancer (fig. S21A, B). As noted previously there were two clones (**Fig. 2A**, clones A and B) which lacked key changes in both chr 8 and 16, with little in common with other cancer-bearing clones (**Fig. 2C**). A spot-wise re-analysis of section H1_2 (including benign spots) revealed that these two clones, now labelled F and G (fig. S21) were spatially grouped as two approximately equal halves of this region of Gleason Grade Group 1 cancer (fig. S21C). This is evidence that low grade prostate cancer is indeed fundamentally distinct from high grade, and raises the hypothesis that such cancer cannot become higher grade because it lacks essential somatic events.

To generalize our findings, we first performed validation through an additional cross-section of a prostatectomy sample that confirmed the spatial continuum of benign clones in proximity to cancer with shared truncal events. We confirmed the high degree of ITH of iCNV clones within prostate tumour loci (fig. S22-26, table S3), and the presence of key somatic events in benign prostate glands. To further generalize we corroborated our findings in multiple organs (**Fig. 4**, fig. S27, table S4-8). While some of the samples did not have an annotated benign reference set, interestingly, unsupervised iCNV could still segment different histological clones. However, the lack of a reference set did reduce the ability to identify specific inferred CNVs (fig. S28). First, we analysed skin tissue with histology including areas of both benign squamous epithelia and squamous cell carcinoma (SCC). For this, we obtained a patient-matched, benign reference set of RNA sequenced single skin cells with confirmation from adjacent sections of benign histology^17^. Spatial iCNV identified four clones within the tissue, one of which corresponded to SCC containing several CN events. Importantly two key events (partial chr 1 and 12 gain) were shared with another nearby clone composed entirely of histologically benign tissue (**Fig. 4B**). To contrast the tumour enriched samples, we analysed a benign lymph node displaying distinct gene expression clusters for the different histological entities (such as germinal centres) and the iCNV analysis provided a copy neutral profile for the entire tissue section (**Fig. 4C, D**). This corroborated further the distinction of gene expression and iCNV. Extending our investigation of the heterogeneity of genomic clones within tumours, we performed additional analysis of other types of tumours: ductal breast cancer and an adult glioblastoma sample (**Fig. 4E-H**). Here we observe a multifaceted spatial iCNV tumour landscape with multiple co-existing clone types in histological similar-appearing tumour tissue. For example, in the case of ductal breast cancer (**Fig. 4E, F**) we observed two distinct clone types (C and F), separated by stroma, with little or no CNV overlap (**Fig. 4F**, fig. S29). In the glioblastoma tissue we similarly identify five clone types that have sharp spatial demarcations separating the iCNV clones, despite being histologically similar (**Fig. 4H**). Overall, the clonal appearances of ITH are clear as well as the overlap with tumour morphology. To contrast these observations, we performed analysis of a sonic hedgehog (SHH) paediatric medulloblastomas (**Fig. 4I-J**) with sex and age matched samples as reference benign samples cannot be attained for these brain tumours. The results show a uniformly homogeneous spatial iCNV clone type throughout the tumour with key expected genetic alterations such as 3q gain (encoding *PIK3CA*) and a 9q deletion (encoding the *PTCH1* gene) as well as small partial gain in 9p. These findings were validated by whole genome sequencing (WGS) of the tumour displaying distinct CNV calls for the three altered chromosomal regions identified by our iCNV analysis (fig. S30-32). The tissue clone diversity over the five investigated tissue types is strikingly variable from homogenous to highly variable genomes in both tumours and benign tissue (**Fig. 4K**). Combining the inferred CNV information with the spatial gene expression patterns, which provides some functional understanding, and cell type mapping (using scRNAseq) could enable targeted treatment options for individual clones, ‘benign’ or tumour, that would not be easily attainable by any other means.

**Fig. 4.**
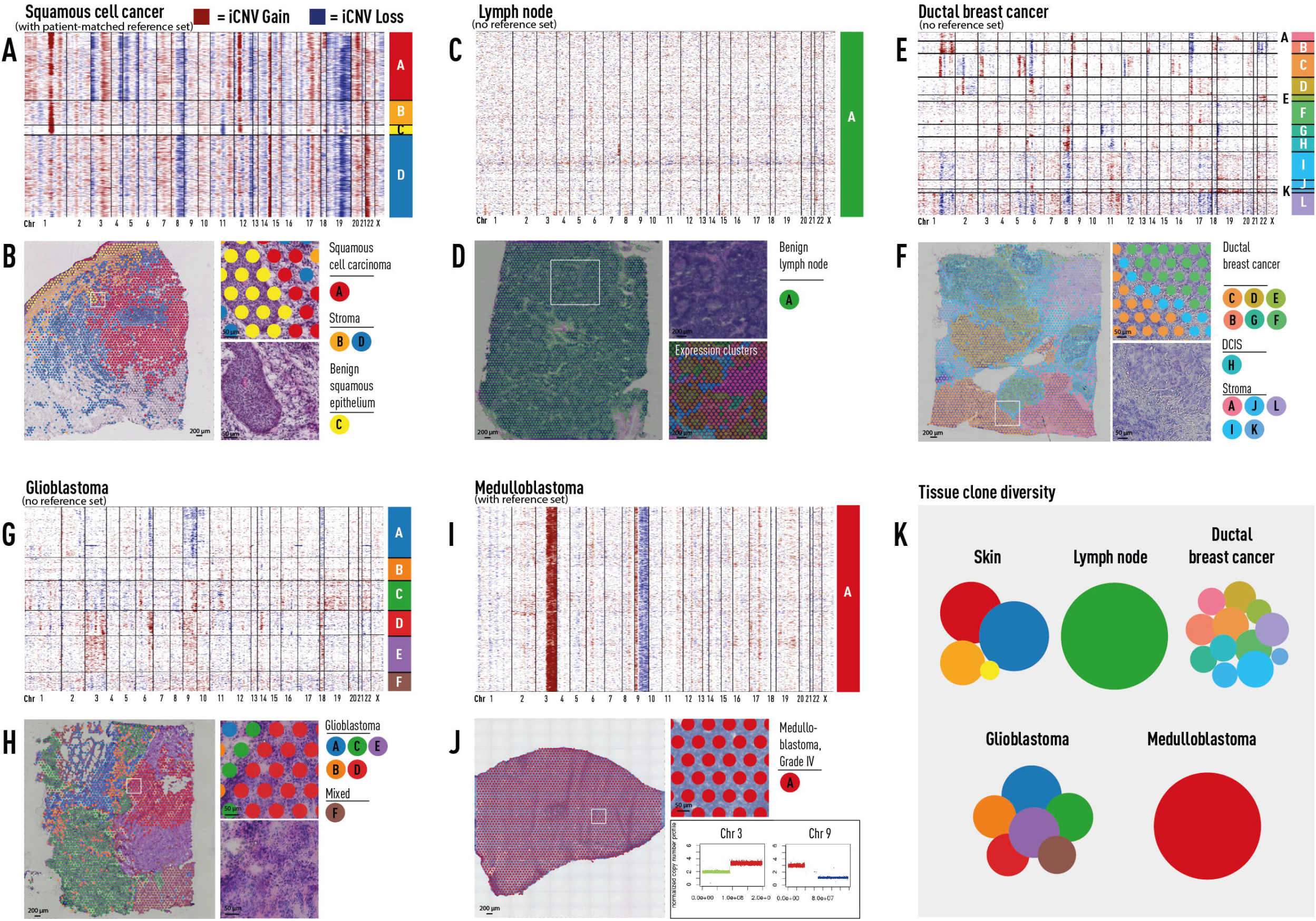
Somatic copy-number alterations in cancer and benign histologies. **(A-B)** Skin containing squamous cell carcinoma (Clone A, red) as well as benign squamous epithelium (Clone C, yellow). A subset of somatic events visualized in cancer Clone A are also detected in adjacent benign epithelial Clone C. Panel B demonstrates representative spot placement for adjacent benign and cancer regions. **(C-D)** Benign lymph node with distinct histological features and gene-expression heterogeneity (Panel D) harbouring no detected copy-number alterations (Panel C). Gene expression clusters determined by UMAP (Panel D). **(E-H)** Breast tissue containing ductal breast cancer and DCIS (E and F). Brain tissue containing glioblastoma (G and H). Genome-wide iCNV clones display spatial tumour clonal heterogeneity. **(I and J)** A monoclonal childhood medulloblastoma. iCNV in Chr 3 and 9 (Panel I), were corroborated by CN calls from WGS (Panel J, lower right). **(K)** Clone distribution per tissue type. Circle area corresponds to numbers of spots per clone. Chr=chromosome. InfCN = inferred Copy-Number Variant. UMAP = Uniform Manifold Approximation and Projection.

In summary, we show that spatial transcriptomic data across multiple cancer types can robustly be used to infer copy number variation, as validated by FISH and WGS. Specifically, we performed an in-depth spatial analysis of prostate cancer that included an unprecedented interrogation of up to 50 000 tissue domains in a single patient, and 120 000 tissue domains across 10 patients. For these domains we inferred genome-wide information in each spot, which facilitated data driven clustering in a tissue wide fashion at high resolution. Furthermore, the spatial information allowed us to identify small clonal units not evident from morphology and hence would be overlooked by histologically-guided laser microdissection or even random sampling of single cells. We continue to show that in some tumour types, particularly prostate, glioma and breast cancers, CNV analysis reveals distinct clonal patterns within tumours. Focusing on prostate cancer, those patterns, as defined by the conservation of CNVs across regions of the tumour, indicate hitherto unappreciated molecular relationships between histologically benign and cancerous regions. It is known that CNVs occur early in tumorigenesis^16^, we propose that CNVs can precede tumorigenesis and are a feature of glandular morphogenesis, with propagation of particular variants traversing disease pathology. This study shows that CNVs in regions of the genome that encode cancer drivers are truly early events, occurring in tissue regions currently unknown to and therefore ignored by pathologists. Currently the risk stratification delivered by pathologists dictates to a significant degree treatment decisions and subsequent clinical outcome. Our study adds an important new approach to the armamentarium of cancer molecular pathology, and should lead to improved early detection of clinically important cancers and improve patient outcomes for ubiquitous malignancies such as prostate cancer. It also raises important questions about cancer evolution and we expect our approach to be of interest to researchers investigating the biological basis of somatic mosaicism and tissue development.

## Acknowledgments

The authors acknowledge The Swedish Childhood Tumor Biobank, supported by The Swedish Childhood Cancer Fund, for access and handling of patient biobank material/sequencing data, the National Genomics Infrastructure (NGI), Sweden for providing infrastructure support. We also acknowledge the Oxford Biomedical Research Computing (BMRC) facility, a joint development between the Wellcome Centre for Human Genetics and the Big Data Institute supported by Health Data Research UK and the NIHR Oxford Biomedical Research Centre. We thank Annelie Mollbrink, Monica Nistér and Anders Ståhls, for helpful assistance and discussions.

## Funding

The Swedish Research Council (KT)

The Swedish Cancer Society (MH, MM, LB, ALJ, PAK, TH)

The Swedish Childhood Cancer Fund (LK, AS, EB, TDDS)

Swedish Foundation for Strategic Research (EB, NS, LL, FT, AT, TL)

Knut and Alice Wallenberg Foundation (AA, JB, LL, RM)

Science for Life Laboratory/ Postdoc program (JM)

European Research Council Advanced Grant (JL)

Cancer Research UK Clinician Scientist Fellowship C57899/A25812 (ADL, AE)

John Black Charitable Foundation (IGM)

Cancer Foundation Finland 180132; Hospital District of Helsinki and Uusimaa TYH2019235; Academy of Finland 304667 (TM)

NIHR Oxford Biomedical Research Centre - Innovation & Evaluation Theme (ADL, AE, IGM, FCH, DJW, RC)

## Author contributions

EB, MH, MM, RM, and NS performed the experiments, AE, LB, LK, JB, LL, TR, KT, ALJ, MM, TDDS analyzed the data. LB and JM developed the factorization approach, AA designed and implemented the method for synthetic data generation, AS, EB, GB, ALJ, PAK, FT, AT, RC, TM, FCH, DJW and TH provided tissue, pathological annotation and biological insight. ADL, IGM, J.L, wrote the manuscript. All authors read and approved the final manuscript.

Correspondence should be addressed to Alastair Lamb or Joakim Lundeberg.

## Ethics declaration

The study was performed according to the Declaration of Helsinki, Basel Declaration and Good Clinical Practice. The study was approved by the Regional Ethical Review Board (REPN) Uppsala, Sweden before study initiation (Dnr 2011/066/2, Landstinget Västmanland, Sari Stenius), Regional Ethical Review Board (EPN), Stockholm, Sweden (DNR 2018/3-31, Monica Nister). All human subjects were provided with full and adequate verbal and written information about the study before their participation. Written informed consent was obtained from all participating subjects before enrolment in the study.

## Data and materials availability

Sequence data have been deposited at the European Genome-Phenome Archive (EGA, www.ebi.ac.uk/ega/), which is hosted by the European Bioinformatics Institute (EBI), under accession numbers EGAXXX. Raw fastq files for the childhood brain tumor samples are available through a Materials transfer agreement with M.N. / The Swedish Childhood Tumor Biobank, in line with GDPR regulations” Count matrices and high-resolution histological, are available on Mendeley: https://data.mendeley.com/datasets/svw96g68dv/draft?a=3f263217-2bd3-4a3c-8125-8c517c3a9e29

Details of the ST analysis pipeline can be found at https://github.com/SpatialTranscriptomicsResearch/st_pipeline.

The factor analysis software (STD) is available under the GNU General Public License v3 at https://github.com/SpatialTranscriptomicsResearch/std-nb.

