## Supplementary Text and Figures for "The spatial landscape of clonal somatic mutations in benign and malignant tissue"

**Other Supplementary Materials for this manuscript include the following:**

Supplementary Data D1 to D2
D1 – Clone Tree Annotations Tables
D2 – Synthetic data files

### Materials and Methods

Tissue specimens

Whole prostates were obtained by open radical prostatectomy at Västerås Hospital. Each prostate was cut in two halves by a horizontal cut, the upper part (closest to the patient's head) was used and cut on a stepped 5 mm mould to obtain a 5 mm high cylinder. Next, stripes were cut out from the cylinder, and each stripe was cut into smaller cubes (total 21 for patient 1 and 28 for patient 2). All tissue cubes were fresh-frozen in liquid nitrogen and stored at −80 °C until embedding for cryosectioning. The childhood brain tumours were collected and provided by The Swedish Childhood Tumour Biobank and stored at −80 °C until embedding for cryosectioning.

Datasets

Human squamous cell carcinoma and case-matched dissociated normal skin cells (reference set) were provided from a published dataset[1]. The human lymph node, human adult glioblastoma multiforme (tumour grade IV) and human breast cancer (ductal carcinoma in situ, lobular carcinoma in situ, invasive carcinoma) datasets were provided from 10x Genomics (https://support.10xgenomics.com/spatial-gene-expression/datasets).

Spatial Transcriptomics (1k arrays)

For prostate (patient 1), all 21 tissue cubes were cryosectioned into 10 µm sections from the bottom (two sections per cube) for ST analysis. The sections were mounted onto spatially barcoded microarray slides. The protocol described in Ståhl et al. and Salmén et al was used to prepare all mounted sections with few modifications[2,3]. Fixation on the slide surface was performed using 4% formaldehyde for 10 min at room temperature. The sections were stained using standard H&E staining and imaged through bright-field imaging to identify and record the tissue histology. Tissue sections were permeabilized using exonuclease I buffer for 30 min at 37 °C and 0.1X pepsin (pH 1) for 10 min at 37 °C. Polyadenylated transcripts are captured by the surface probes, which function as primers in the overnight reverse transcription reaction (42 °C). The release of the cDNA–mRNA hybrids and subsequent steps were performed exactly as described in the accompanying protocol by Ståhl et al and Salmén et al[2,3]. The material was processed into libraries as described in Jemt et al. and sequenced on an Illumina Novaseq using paired-end 300-bp reads[4].

Spatial Transcriptomics (10X Genomics Visium)

Visium Spatial Tissue Optimization Slide & Reagent kit (10x Genomics, Pleasanton, CA, USA) was used to optimize permeabilization conditions for the tissue sections. One 10µm section from each patient was processed according to the manufacturer's instructions. Spatially barcoded cDNA of every tissue section was generated using Visium Spatial Gene Expression Slide & Reagent kit (10x Genomics). 10µm thick tissue sections from prostate patient 1 were fixed according to manufacturer's instructions and permeabilization was performed for 8 min. Sections of the same thickness from prostate patient 2 were fixed for 10 min using acetone at -20 °C and permeabilized for 15 min. Childhood brain tumour sections of 12µm were processed according to manufacturer’s instructions and permeabilized for 30 min. Libraries for all tissue sections were generated following 10x Genomics Visium library preparation protocol and sequenced on Illumina sequencing instruments.

Data processing

1k arrays: FASTQ files were processed using the ST Pipeline v.1.5.1 software[5]. Read 1 contained only a spatial barcode followed by a UMI. Read 2 transcripts were mapped with STAR[6] to the GRCh38.79 human reference genomes. Mapped reads were counted using the HTseq count tool[7]. Spatial barcodes were demultiplexed using an implementation of TagGD[8] UMI filtering is carried out to remove duplicated reads. The output is a matrix with gene counts for each spatial barcode.

10X Visium arrays: Specifics regarding data processing prior to data analysis after demultiplexing of fastq files has been described elsewhere for the human squamous cell carcinoma[1] and datasets provided from 10X Genomics (https://support.10xgenomics.com/spatial-gene-expression/datasets). For the childhood brain tumour read 2 was trimmed to remove both the TSO adapter sequence and polyA homopolymers using Cutadapt[9]. In brief, both the full-length and partial TSO adapter sequences were removed by setting a non-internal 5' adapter (error tolerance = 0.1 and minimum overlap = 5) to the TSO sequence (AAGCAGTGGTATCAACGCAGAGTACATGGG). By setting a sequence of 10 A's for a 3' adaptor (minimum overlap = 5) the polyA homopolymer sequences were removed. Raw read 1 and the trimmed read 2 fastq files were then run through Space Ranger (version 1.0.0, 10X Genomics) where reads were mapped to the human reference genome (GRCh38, release 93). The raw sequencing reads of the prostate samples were directly processed using Space Ranger (version 1.0.0 prostate 1, version 1.2.1 prostate 2, 10X Genomics) and mapped using the same human reference genome as above.

Factorized negative binomial regression of prostate samples

To deconvolve the expression data into GEFs, we developed a method for factorized negative binomial regression. Briefly, we assume that the observed number of transcripts for a given spot *s* and gene *g* is a sum of hidden counts *y* from *T* GEFs,

$$x_{sg} = \sum_{t=1}^{T} y_{stg}$$

We assume that the hidden count $y_{gt}$ follows a negative binomial distribution with a type-specific rate. Model parameters are found by maximum a posteriori estimation and used to infer $y_{gt}$. To visualize the GEFs spatially, we first reduce the relative hidden counts in each spot to three dimensions using UMAP. The resulting data is scaled into the unit cube using a channel-wise affine transformation and used as RGB color coordinates. In all analyses, we factorized the data into $T$ = 25, 24 & 20 GEFs (1k, Visium patient 1 and Visium patient 2) and ran the optimization for 5000 iterations. Spots were annotated based on their section to control for sample-wise batch effects.

Processing and visualization of non-prostate samples

Data processing and visualization was carried out using Seurat (version 3.2.2)[10] and STUtility (version 0.1.0)[11] R packages. Filtering of UMI counts was performed using the *InputFromTable* function in STUtility. Genes were removed if they were present in less than 5 spots or had a total UMI count below 100. All spots containing less than 500 UMI counts were also removed. Samples were normalized using variance stabilizing transformation implemented in the

*SCTransform* function from Seurat. Dimensionality reduction was performed using principal component analysis (*RunPCA*) and the top principal components were selected as follows: top 20 components for human squamous cell carcinoma, human lymph node, human glioblastoma multiforme, human invasive ductal breast carcinoma and top 10 components for the childhood brain tumour were used. The expression-based clustering was done by, (1) constructing a shared nearest neighbour (SNN) graph (*FindNeighbors*) from the previously established components and (2) running *FindCluster*s with the following resolution parameter settings: res = 0.8 for human squamous cell carcinoma, human lymph node, human glioblastoma multiforme and human invasive ductal carcinoma and res = 0.2 for the childhood brain tumour. Finally, a two-dimensional UMAP embedding was constructed from the previously established top principal components for each tissue type. For the human lymph node, differentially expressed genes for each Seurat cluster was determined using the *FindAllMarkers* function, only testing genes detected in at least 25% of the spots in either of the two populations, i.e. cluster or background.

DNA fluorescence in situ hybridization (FISH)

Optimal cutting temperature (OCT)-embedded block of fresh frozen prostate sample was sectioned at 5µm thickness and several consecutive sections were mounted on positively charged microscope slides (VWR, catalogue number MENZJ5800AMNZ) and placed at -80 °C until processing. Sections were fixed with methanol and acetic acid (3:1 ratio) for 15 minutes at room temperature, washed in 1x PBS for a few times and briefly air-dried. For H&E staining, one of the sections was set aside and stained for 4 minutes in hematoxylin (Sigma, catalogue number MHS32), followed by 1 minute in eosin (Sigma, catalogue number HT110216). These were used by histo-pathologists to confirm regions of interest as epithelial.

For FISH experiment, DNA FISH probes containing *MYC*/8cen (Cytocell, catalogue number MPD28000) or *PTEN*/10cent (Cytocell, catalogue number MPD15000) were added (10-15µl) on top of tissue sections, sandwiched with 18x18 coverslips and sealed with rubber glue (BioNordika AB, catalogue number PCN009). Slides were placed on a hot plate for exactly 6 minutes at 76 °C for DNA molecules to denature and immediately placed inside an incubator with 100% humidity for overnight incubation at 37 °C. Next day, coverslips were gently removed and slides were washed in a ceramic jar containing prewarmed 0.4x SSC for 3 minutes at 72 °C, transferred to 2x SSC/0.05 TWEEN® 20 for 2 minutes at room temperature, then quickly washed in 2x SCC and nuclease free water. In order to reduce the autofluorescence backgrounds, we applied quenching probes (Thermo Fisher Scientific, catalogue number R37630) on top of sections, incubated for 5 minutes at room temperature, washed in 1x PBS, nuclei were counterstained with DAPI and slides were mounted using mounting medium (Thermo Fisher Scientific, catalogue number S36936). Microscopy images were acquired using a ×100 1.45 NA objective mounted on an Eclipse inverted microscope system (Nikon) controlled by the NIS Elements. We collected multiple image stacks per sample, each consisting of 30-40 focal planes spaced 0.3µm apart.

Pathologist Workflow – Spot-level annotation for prostate patient 1

All Visium spots were annotated on a spot-by-spot basis using the Loupe Browser Version 5.0 (10x Genomics) for the Visium sections by two board-certified uro-pathologists (R.C. and T.M.). Using a >50% cellular coverage threshold, the pathologists annotated spots by histological class, or "Exclude" (eg. mixed coverage, array regions not covering tissue such as lumens, or if scanning/sectioning artefact rendered it impossible to determine a histological class). The resultant annotations were exported as CSV files, cleaned and unified as annotation classes, which returned an output file of merged annotations where annotation classes were discrepant. These were then exported as a CSV file for visualization in Loupe Browser for review.

Next, a consensus workflow for revising the discrepant annotations was performed, where-in R.C. and T.M. were asked to determine a final annotation class if there were discrepancies between benign or cancerous luminal epithelial cells. If there were discrepancies between luminal classes and stroma, A.E. performed review and re-classification, such that if over 50% of cells of one class could be identified, it was marked as the corresponding class. If there was uncertainty, the spot was marked as "mixed" and excluded from downstream analysis.

The final consensus annotation dataset consisted of a total n = 23 282, with the following distribution: Benign (Benign, Benign*, n = 5 942), Cancer (GG1, GG2, GG4, GG4 Cribriform, n = 6 356), Stroma (n = 8 623), PIN (n = 36), Inflammation (Inflammation, Chronic Inflammation, n = 38), Exclude (Exclude, Blank, n = 1 983), Fat (n = 137), Nerve (n = 33), Transition State (n = 117) and Vessel (n = 17).

We defined low-grade prostate cancers as Gleason Grade Group 1 and high-grade cancer as containing Gleason pattern 4.

Pathologist Workflow – Spot-level annotation for prostate patient 2

Visium data from 15 prostate sections were generated for patient 2. We used Loupe Browser files to annotate prostatic luminal epithelial cells, which were visually confirmed by pathologist T.M. The 15 prostate sections were annotated (A.E. and T.M.) for the presence of tumour histology, which was confirmed in sections H3_1, H2_1, H2_2, and H3_6. From the remainder of the 11 prostate tissue sections from patient 2 which did not contain tumour, luminal epithelial cells were analysed for selection of a benign reference set (fig. S26). Analysis of the histologically benign, luminal epithelial cells identified spatially distinct clones in section H3_2 (physically adjacent to the three tumour-bearing sections H3_1, H2_1, H2_2), and this section therefore was included in a joint SpatialInferCNV analysis of including the tumour sections using standard inferCNV parameters (Supplemental Methods - section **InferCNV - Parameters**).

InferCNV – Data Pre-processing

In order to systematically interrogate the data, we developed an R package called SpatialInferCNV: *https://github.com/aerickso/SpatialInferCNV*. Additional analyses were performed using a series of R packages (tidyverse, Seurat, infercnv, hdf5r), python, and BASH scripts as follows (fig. S1).

Histological annotations were imported from the final annotation consensus files for all sections, and the barcodes were appended with information corresponding to their section. Next, the annotations were filtered for a given feature of interest. Files output from the cell ranger pipeline (filtered_feature_bc_matrix.h5) were imported, and barcodes were appended with information of their corresponding section name. The count files were then filtered only for those within the analysis of interest. The count files further underwent a quality control (QC) filter[3] wherein spots containing 500 counts or less were removed. The annotations file and counts file were joined for each section, and these were then all combined into a final matrix that was then output (.tsv file) for downstream analysis with inferCNV. The barcodes for only those that passed the annotation and QC filters were merged again with the annotations, and these were separately exported (.tsv) files for further InferCNV analysis. Lastly, a genomic positions file was created following the instructions here: [https://github.com/broadinstitute/inferCNV/wiki/instructions-create-genome-position-file](https://github.com/broadinstitute/inferCNV/wiki/instructions-create-genome-position-file). These analyses were then run on a high-performance cluster.

Selection of Benign References

Inputs to InferCNV can include a reference set of UMI-barcoded objects, in order to improve precise inference of genomic CN events in the observed population. Common analyses can include a comparison of benign references to observed cancers. With this in mind, we wished to quality control our benign references in order to identify histologically benign cells with little-to-no structured CN events, to enable us to identify CN events within the observed (tumour) population. We first performed an unsupervised analysis of only the benign reference cells: (Parameters for inferCNV object: ref_group_names=NULL, Parameters for run: cutoff=0.1, cluster_by_groups=FALSE, denoise=TRUE). Using the denoised outputs, we identified by visual inspection a subgroup of all benign that harboured little-to-no inferredCNVs (fig. S16). The associated dendrogram file (harbouring the cluster structure and each barcode therein) was then further analysed in a R script for node selection.

InferCNV Parameters

For unsupervised SpatialInferCNV analysis, in addition to standard parameters and in order to include data from Chromosomes X and Y, we included the following parameter for the function CreateInfercnvObject(): chr_exclude = c("chrM"). For the run() function, we used the following parameter values: the following inferCNV run() parameters were used: “cutoff=0.1, num_threads = 10, cluster_by_groups=FALSE, denoise=TRUE, HMM=FALSE”.

In supervised SpatialInferCNV analysis (to call inferCNV’s Hidden-Markov Model functions), InferCNV was run as follows. The node identity file was used in place of the annotation file. The following “InferCNV run” parameters were used: “cutoff=0.1, num_threads = 10, cluster_by_groups=TRUE, denoise=TRUE, HMM=TRUE). Files of interest relevant for this manuscript are the default outputs: (infercnv.21_denoised.png, infercnv.17_HMM_predHMMi6.hmm_mode-samples.png, 17_HMM_predHMMi6.hmm_mode-samples.pred_cnv_regions.dat, 17_HMM_predHMMi6.hmm_mode-samples.pred_cnv_genes.dat, and 17_HMM_predHMMi6.hmm_mode-samples.genes_used.dat).

For the global visualization of iCNV events in Figure 1, we analysed spatial transcriptomics (1k arrays) data with inferCNV for all 21 sections in a global analysis without a reference set. We performed the analysis such that each individual spatial transcriptomics spot was ran with the following inferCNV run() parameters were used: cutoff=0.1, num_threads = 10, cluster_by_groups=FALSE, denoise=TRUE, HMM=TRUE, analysis_mode = "cells", HMM_report_by = "cell"”.

To spatially visualize global iCNV profiles across an entire prostate, we then determined the number of individual genes detected to harbour an inferred CN gain or loss. To reduce background noise in the visualization, the resultant HMM calls were thresholded for the number of gene-level iCNV events present in at least 35% of all spots across the entire dataset, and in at least 45% of the spots of a given section. These thresholds were selected after detailed interrogation of thresholds ranging from 10-90% in 5% increments with positive, neutral, and negative control sections for visual consistency.

Clone Selection

The dendrogram file was imported into R, and the dendrogram tree with numerical node identities was visualized. Dendrogram nodes were extracted, and the specific node members (Visium Spots) were digitally selected and assigned a clone identity. All members of a given analysis were merged, and a CSV file containing the clone identity and the barcode was output for each Visium section.

Clone Visualization

Loupe Browser Version 5.0 (10x Genomics) was used to spatially visualize resultant clones from clone selection. For the manuscript, if a clone in a given section had ≤ 10 1k or Visium spots, it was not visualized.

Clone Tree consensus iCNV event calling

Both HMM iCNVs, and manual interpretation of denoised outputs were used to identify putative subclonal CNVs. These were then merged in a final consensus set for building clone trees (Supplementary Data S1). Briefly, trees were constructed by identifying where CNVs were shared across clusters identified above as, under the assumption that a CNV cannot be reversed once it occurs, this indicates the cells in those clusters share a common ancestry. We therefore used this logic to identify ancestral relationships between clusters and build the clone tree.

More formally, each cluster, $C_i, i \in 1,2,\ldots,n$ was represented by the CNVs that were overved in the cluster $C_i = \{c_q\}$, where $q$ denotes the index of a subset of CNVs from the full consensus set. We established a set of putative clones, $\{Z_j\}$, that were defined as the pairwise intersection of all clusters ($Z_j = C_a \cap C_b; a, b \in 1,2,\ldots,n$). Note that the original clusters are included in this set by definition when $a = b$. Starting from a root parental clone corresponding to cells with no CNVs, (i.e., $Z_1 = \emptyset$), trees were constructed by identifying clones that were *immediate descendants* of the parental clone. Immediate descendants were defined as all clones that directly emerged from the common ancestor $Z_{par}$, and were determined by applying three steps

1. Descendancy: All $Zj$ such that the CNVs present in the parental clone, $Z_{par}$, were a subset of those in the descendant (i.e., fulfils the condition $Z_{par} \subseteq Z_j$).
2. Branching: Of the clones ($Z_j; j = 1,2,\ldots,n_{desc}$) identified in 1), identify branching into multiple descendant lineages by finding those where the pairwise intersection returns only CNVs that are present in the parent ($Z_c \cap Z_d = Z_{par}; c, d = 1,2,\ldots,n_{desc}$).
3. Immediate descendant: The immediate descendant in each lineage is the clone for which there can be no other clones that lie between it and the parent on the tree. If branching is not found ($Z_c \cap Z_d \neq Z_{par} \; \forall \; c, d$), then there is only one descendant lineage, and the immediate descendant is therefore the clone with the fewest

additional CNVs (where $|Z_j|$ is minimized). Otherwise, we can establish clones in the same lineage through the pairwise intersections, such that $Z_c \cap Z_d = \{Z_{par}, Z_{lin}, \ldots\}; c, d = 1, 2, \ldots, n_{desc}$, where $Z_{lin}$ is a set of CNAs common to all clones in that lineage. The immediate descendant is that which has no additional CNAs other than those found in the parent and those that are common to the lineage.

Clone Trees – Branch Lengths

To semi-quantitatively depict the 'evolutionary distance' between subclones, we determined the branch lengths by taking the logarithm (base 2) of the number of additional CNVs in the descendant clone, and adding an arbitrary value to ensure that branches were always visible even with few CNV differences. The formula is given as $b_k = 100\ log_2(|Z_{desc}| - |Z_{par}|) + 300$, where $b_k$ is the length of branch $k$ in pixels.

Clone Trees – Clone Diameters

We scaled the size of each circle denoting a clone by the proportion of spots in the sample that was assigned to a clone using the formula $d_l = 10\ log_2(s_l)$, where $d_l$ denotes the circle diameter in pixels, $s_l$ is the number of spots that correspond to a clone.

SpatialInferCNV Parameters (Figure 4)

Patient-matched, scRNAseq data from dissociated normal skin cells were analysed for selection (previously described) of a benign reference set. This reference set was then used as a reference control for all ST spots in section T28. Node selection was performed (previously described). One pathologist (R.C.) annotated the resultant clones with the percentage of spots for each clone that harboured either "Stroma", "Tumour Epithelia", or "Non-Invasive Epithelia" (table S9).

SpatialInferCNV analysis was performed on the Lymph Node specimen without a reference set using default parameters previously defined (Supplemental Methods - section **InferCNV Parameters**).

Publicly available data for the Breast Tumour specimen was accessed from the 10x Genomics website. InferCNV was run without a reference set using default parameters previously defined (Supplemental Methods - section **InferCNV Parameters**). Node selection was performed (Supplemental Methods - section **Clone Selection**). Two pathologists (T.M. and A.S.) annotated the resultant clones with the percentage of spots for each clone that harboured either "Stroma", "Tumour Epithelia", "DCIS", "Lymphatic cells" (table S10).

For inferCNV analysis of the childhood brain tumour, data were analysed using inferCNV (version 1.7.1) R package[12]. Patient 2 and 3 were selected as reference samples to patient 1. The selected reference samples appeared to demonstrate little to no inferred CNV gains and losses as shown by supplementary fig. S31-32. Whole genome sequencing of patient 3 confirmed the copy neutral genome while patient 2 did show rearrangements withing chr 2. Because Visium and WGS data were generated from different locations of each tumour, we speculate that the observed WGS CNV patterns in patient 2 could be due to the inherent spatial heterogeneity of DNA copy number alterations observed by others when sampling multiple sites of medulloblastoma tumours[13].

RNA vs DNA phylogenies from single-cell (SIDR)

DNA and RNA data, co-extracted from single tumour cells, were obtained from publicly available data repositories[14]. Genomic and transcriptomic libraries were aligned to GRCh38.79. DNA-based CNV profiles were analysed and clustered by GINKGO: https://github.com/robertaboukhalil/ginkgo[15]. RNA-profiles were analysed by inferCNV, without a reference set, using default parameters. Tanglegrams of hierarchical clustering of both DNA-based Copy-Number Profiles and RNA-based inferred Copy-Number Profiles were then analysed by the R package *Dendextend*[16].

RNA vs DNA phylogenies from published prostate data

RNA data were obtained for patient A21 [17,18], patient 499 [19], and Cases 6, 7 and 8 [20]. For patients A21 and 499, only a subset of all specimens had transcript data available. For cases 6, 7, and 8, only RNA microarray data were available, precluding their analyses by inferCNV. The transcriptomes were aligned to GRCh38.79, and RNA counts were obtained. These were then processed into inferCNV objects, and ran with standard inferCNV settings, without a reference set (Supplemental Methods - section **InferCNV - Parameters).** Dendrograms from the inferCNV outputs were then visualized using the R. For Cases 6, 7, and 8, we used R scripts to implement a framework to generate phylogenetic trees directly from transcriptomic data [21].

Synthetic Data – Generative Process

To evaluate our application of the computational method InferCNV to spatial transcriptomics data, we designed a generative process that results in an in silico spatial transcriptomics experiment of a tissue with a known - and spatially structured - clonotype population. In short, we construct a spatial domain (representing a tissue region) in which we place a set of virtual cells with a common genome structure, and then let these cells populate the tissue region by simulating growth. In the process, at every time point, cells can move, generate offspring, die or stay stagnant.

When generating offspring, either of the two mutational events (amplification or deletion) may occur with equal probability, where a set of one or more genes are affected (allowing for co-amplification/deletion to occur). The set of genes that will be affected in the mutational event is determined by first sampling a centre-of-event ($r$) location uniformly from the genome, the spread ($w$) of the event is then determined by sampling from a truncated (at zero) normal distribution with mean zero and standard deviation $\sigma$. All genes that are completely contained within the closed interval $[r-w, r+w]$ on the genome are then included in the affected set.

The tissue is allowed to grow until either the maximal number of allowed iterations is reached or if the number of cells in the tissue exceeds an upper limit. After growth, a user-specified fraction of cells will be randomly removed (a zero value equates to skipping this step), creating additional "empty" regions in the tumour foci, to emulate intermixing of tumour and normal cells. All unpopulated regions of the tissue domain, inside as well as outside of the foci, are then filled with "normal" cells (no copy number events).

During growth, we also introduce a two state Markov model to allow for different cycles of more intensive mutation versus expansion of clones in our tissue. In the end, we will have a mixture of normal and aberrant cells, with different genome profiles. To promote synchrony within the population, we force daughter cells to stay in the parent's state for the same number of iterations as the parent, after which the two's states are decoupled.

Every cell has the same baseline expression propensity for each gene, but these propensities may change as a consequence of mutational events. The baseline expression propensities were sampled from a Dirichlet distribution with concentration parameter 1, generating a vector (of the same dimensionality as the number of genes) on the probability simplex, these values were shared among all cells (aberrant as well as benign), and we referred to them as $\theta$-values. However, to account for the influence of copy number events, we used a modified value of $\theta$, denoting the adjusted $\theta$-values as $\theta'$. Finally, to sample gene expression vectors, we used a multinomial distribution with $\theta'$ as the probability vector. Hence, we may describe the process used to generate the cell-level gene expression profiles as in Eq. 1:

$$\theta \sim Dir(1, \dots, 1)$$
$$y_c \sim Mult(N, \theta'), \theta' = \frac{\theta * n_{cg}}{\sum_{k \in G} \theta * n_{kg}} \quad \text{(Eq. 1)}$$

Where $y_c$ is the gene expression vector of cell $c$, $n_{cg}$ is the number of copies of gene $g$ in cell $c$, and $N$ is the number of transcripts we seek to sample. To clarify, sampling of $\theta$ only occurs once, more precisely, when we initialize the system.

Having established a procedure to simulate tissue growth and sample transcripts from each cell in the tissue's cell population, we proceed to outline how we used these features in order to generate data similar to the spatial transcriptomics data in our study. First, a regular grid was constructed over the tissue domain, with each node representing a spot in the real spatial transcriptomics array. All spots had the same radius, and cells within this distance from a given spot's centre were assigned to said spot. Once cells had been assigned to spots, we sampled a gene expression vector from each cell in the tissue, and let all the transcripts from cells assigned to spot s constitute the pool of available transcript at said spot ($z_s$). Finally, we randomly drew $M$ transcripts from each spot's pool of available transcripts to constitute the spatial gene expression vector. More formally, we may express this as Eq. 2:

$$x_s \sim Mult(M, z_s), \; z_{sg} = \frac{\hat{z}_{sg}}{\sum_k \hat{z}_{sk}}, \; \hat{z}_{sg} = \frac{1}{|C_s|} \sum_{c \in C_s} y_{cg} \quad \text{(Eq. 2)}$$

Where $x_s$ is the expression vector for spot $s$, $C_s$ is the set of cells residing at spots $s$, and $| \, . \, |$ denotes the cardinality operator.

The cell-to-spot assignments are stored, which allow us to reconstruct the exact genomic profile for the population of cells at each spot; we also summarize this as an "average" genomic profile matrix for easy visualization. Spots that contain at least one aberrant cell (one or more copy number events), were annotated as non-benign. This somewhat stringent criterion allowed us to assess performance in border regions between benign and non-benign tissue, that often is hard to annotate properly.

The generative process described above is implemented in Python code and available as a CLI application that can be accessed at GitHub https://github.com/almaan/growmeatissue. The

GitHub repository also contains more extensive documentation and examples of how to use the code, the exact parameters - defined in a TOML design file - used to produce the presented data are included in supplementary Data S2.

Synthetic Data - Evaluation

The process described above was used to generate a set of synthetic data incorporating a single chromosome, from which the obtained spatial gene expression data together with associated annotations were entered as input to spatial InferCNV, this data can be found in supplementary Data S2. The synthetic data was analysed according to the same procedure as previously outlined for the real data, providing, as output, information regarding the clonal population as determined from the inferred genomic state.

To compare the results with the ground truth we focused exclusively on the set of cells not being used as a reference (non-benign). Spatial InferCNV assigns a state (either three or six depending on which hidden markov model approach is used) to every gene in each clone; we converted these states into a categorization that was more suitable for comparison according to the following scheme, given as "spatial inferCNV state" - "new category": 1 - deletion, 2 - deletion, 3 - neutral, 4 - amplification, 5 - amplification, 6 - amplification. For the ground truth data, we computed the average copy number of all cells assigned to each spot and rounded this value to the nearest integer. We considered a gene (within a clone) as deleted if the rounded average copy number within the given clone was less than one, amplified if it was higher than one, and neutral if it was equal to one. Having cast the two data sets (real and synthetic) into comparable formats, we then computed the accuracy (within each clone) as the number of equal gene annotations (deletion, neutral, amplifications) between the ground truth and the (from spatial InferCNV) inferred results.

Paediatric Tumour DNA sequencing and data analysis

Libraries for WGS were prepared using Illumina TruSeq PCR-free. WGS samples were sequenced 2x150bp paired-end, on HiSeqX v2.5 (sample 810) or NovaSeq6000 (samples 539 and 659) instruments (Illumina, SanDiego, California). DNA sequence data was processed with Sarek, following the GATK best-practice recommendations[22], on UPPMAX Clusters at Uppsala University (https://www.uppmax.uu.se/resources/systems/the-bianca-cluster/). Briefly, the steps run were: quality control of FASTQ files using FASTQC (https://www.bioinformatics.babraham.ac.uk/projects/fastqc/), alignment of short reads to the human reference genome sequence (GRCh38/hg38) using bwa-mem, with the ALT-aware option turned on[23], sorting of reads and marking of PCR duplicates with GATK MarkDuplicates and base quality scores recalibration and joint realignment of reads around insertions and deletions (indels), using GATK tools (https://github.com/broadinstitute). Tumour CNV profiles were generated using Control-FREEC[24]. The matched normal sample was used to call somatic CNVs.

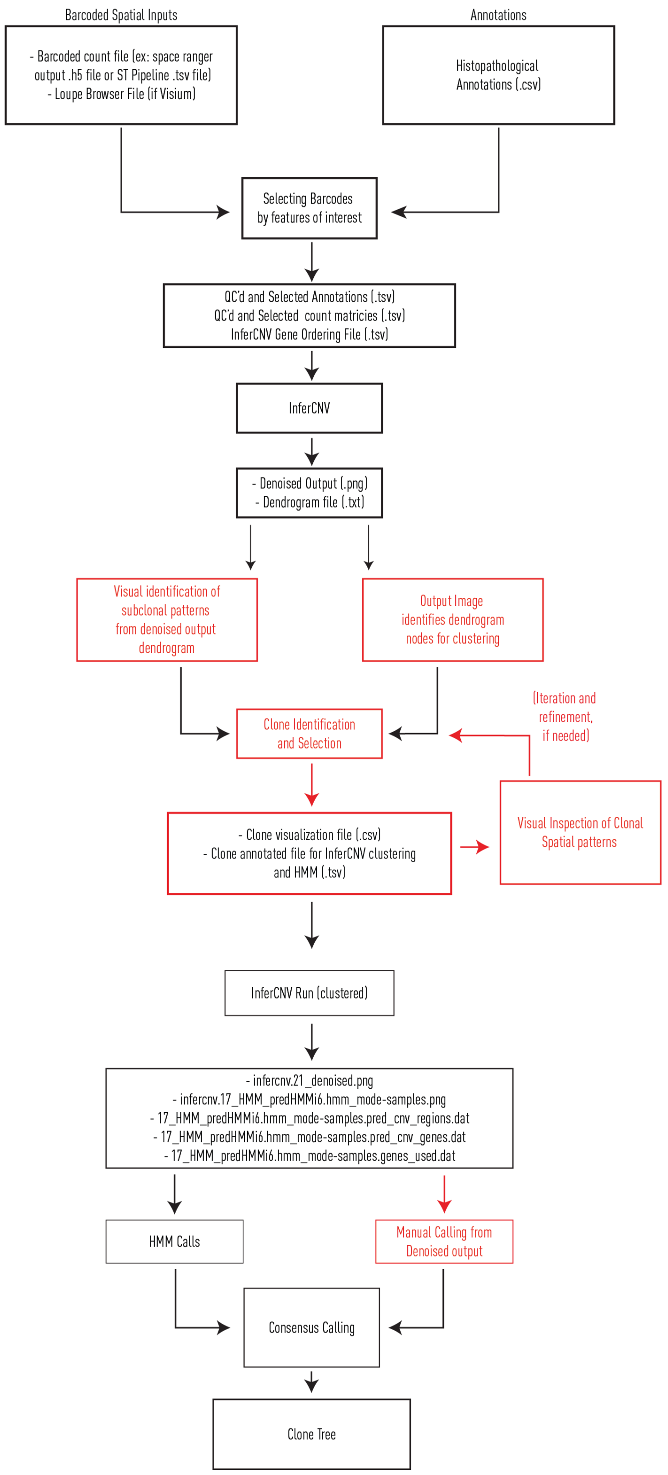


**Fig. S1. Overview workflow diagram for SpatialInferCNV.**
Diagram details input and outputs steps. Red labelling indicates steps requiring manual decision making. Further details can be found at: https://github.com/aerickso/SpatialInferCNV.

HMM = Hidden-Markov Model.

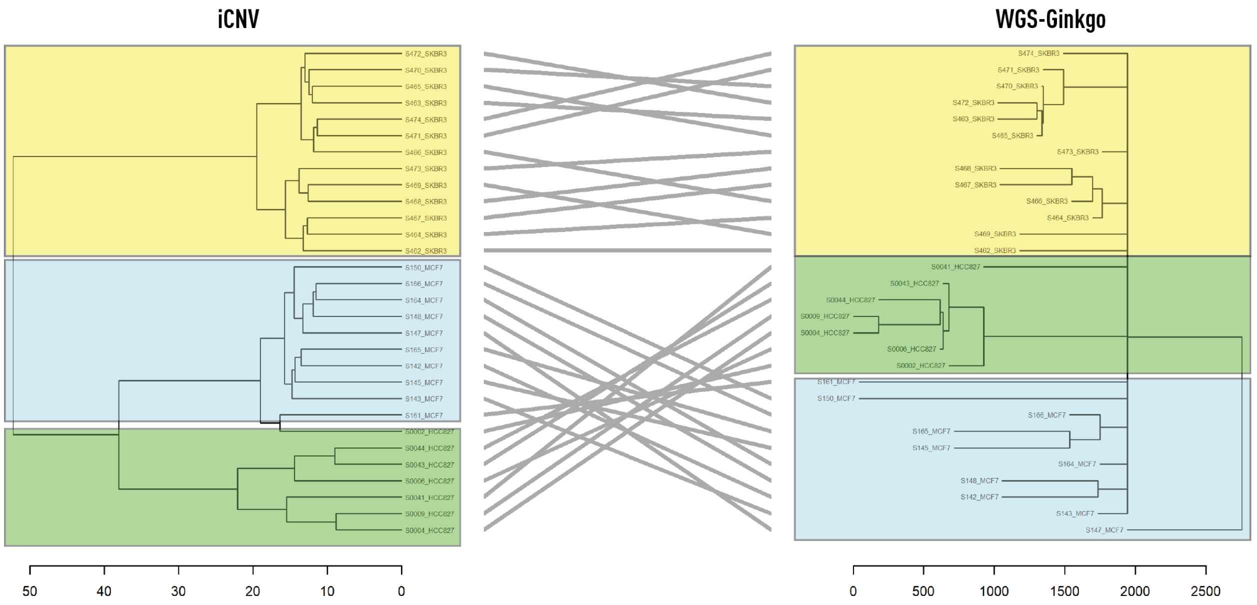


**Fig. S2. Comparison of phylograms created from whole genome sequencing CNVs and iCNV's from single tumour cells with co-isolated DNA and RNA[14].**
Colours correspond to individual cell lines (yellow: SKBR3, green: HCC827, and light blue: MCF7). Entanglement of the phylograms was 0.11 (an entanglement value of 1 corresponds with full entanglement of two phylograms, whereas an entanglement value of 0 corresponds with no entanglement).

iCNV = inferred Copy Number Variant

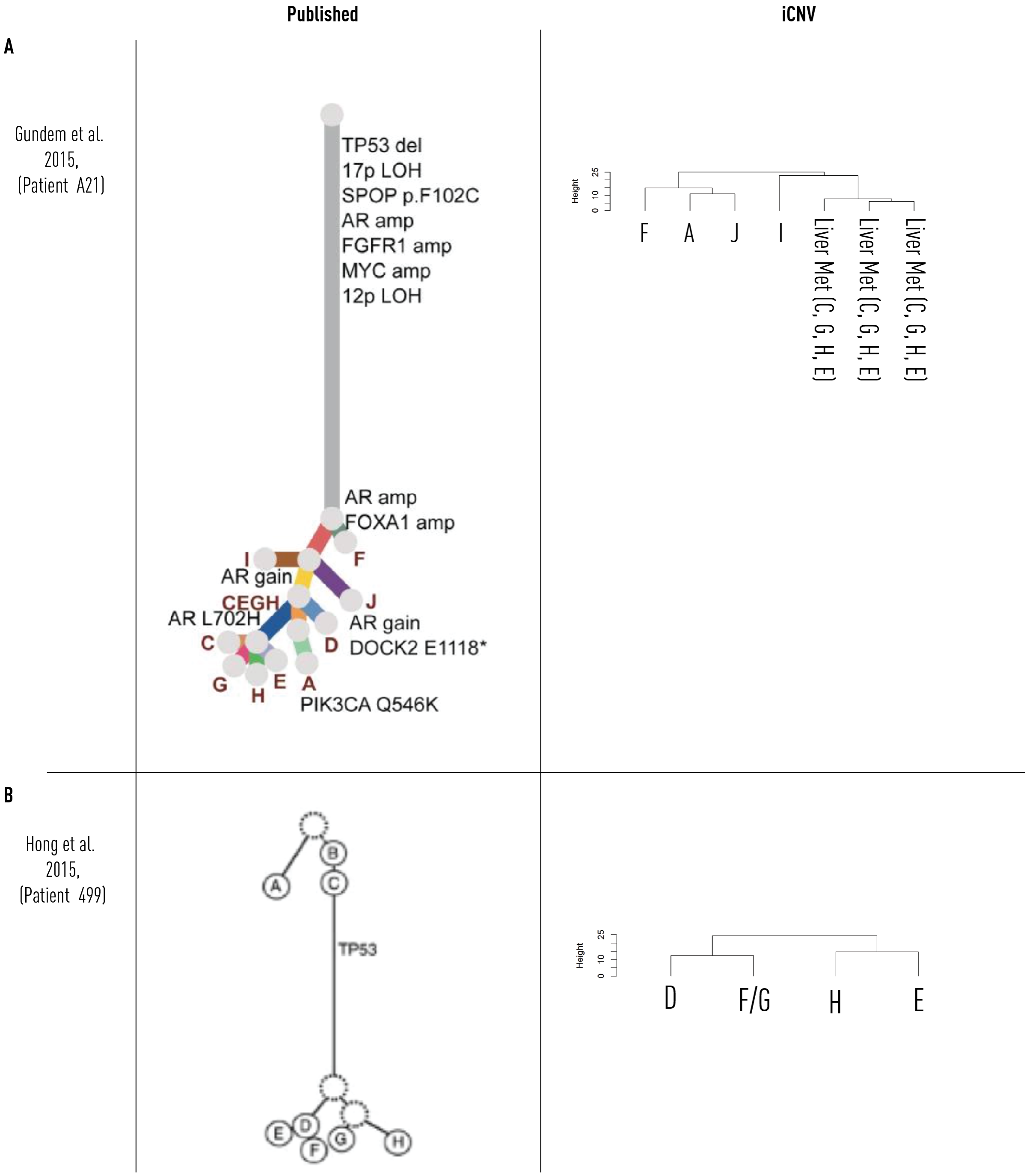


**Fig. S3. Comparison of published WGS-based phylogenies to iCNV and raw-transcript based clustering.**

**(A)** Phylogeny from patient A21, as published and reproduced from Gundem et al.[17]. Transcript data were available only for a subset of specimens. **(B)** Phylogeny from patient 499, as published and reproduced from Hong et al.[19]. Transcript data were available only for a subset of specimens (used to reproduce phylogenetic tree by iCNV).

iCNV = inferred Copy Number Variant

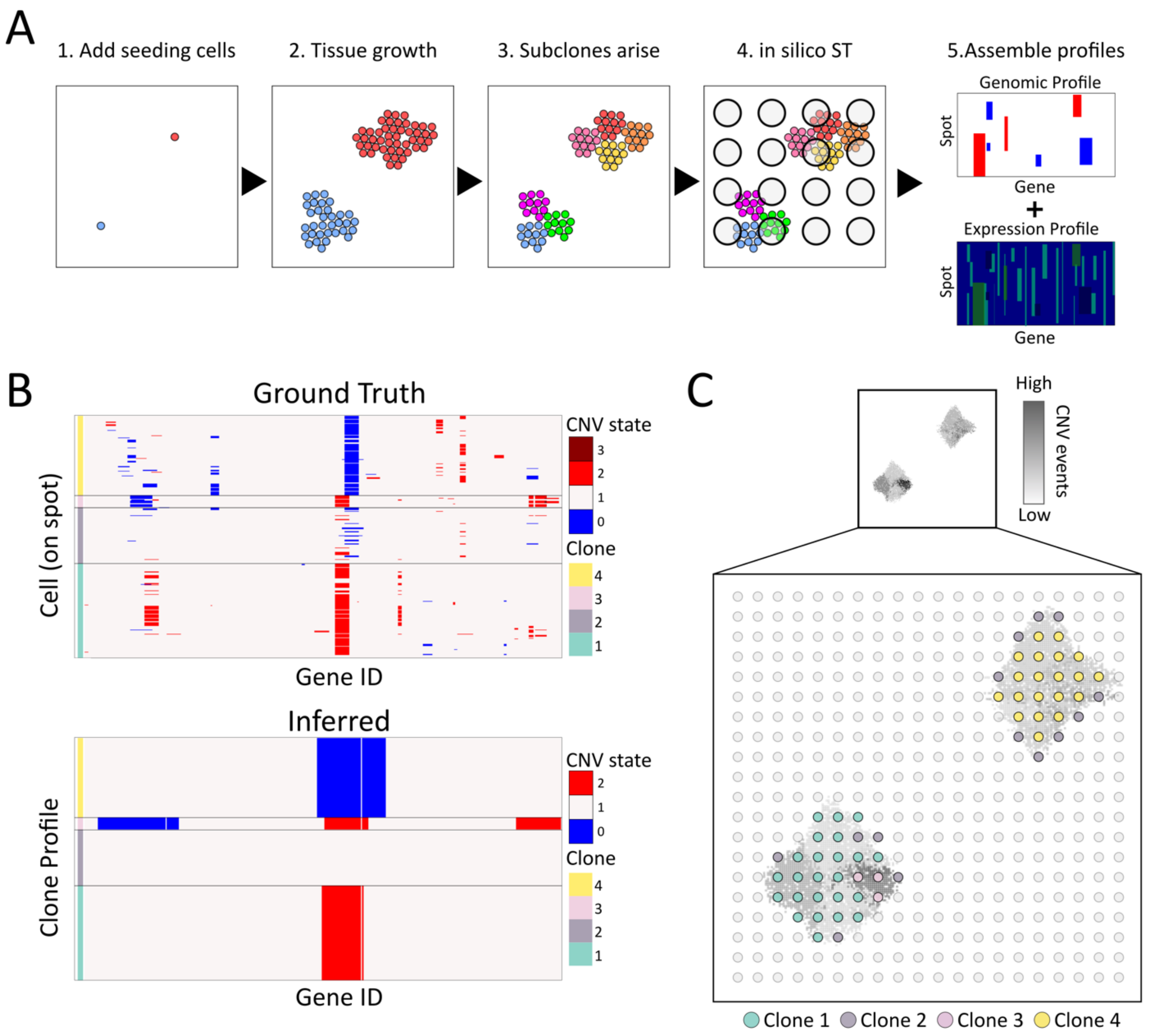


**Fig. S4. iCNV on synthetic data**
**(A)** Schematic overview of the generative process used to produce artificial spatial data. 1) First a set of seeding cells (red and blue circles) are placed in a defined tissue domain (square), every seeding cell hosts one unique copy number event. 2) The cells are allowed to "grow" within the tissue domain until the number of cells in the domain exceeds a predetermined number. 3) Mutations in the genome occur stochastically during growth and as a result, subpopulations (indicated by colour) of cells with similar genomic profiles arise. 4) Unoccupied space in the tissue domain is filled with benign cells (no copy number variations), spatial capture locations are placed in a grid over the grown tissue and transcripts are "captured" from the cells overlying each spot. 5) Synthetic spatial expression data is produced together with associated ground truth genomic data (both on spot and cell level). **(B)** Results from applying spatial inferCNV (bottom) to a set of synthetic data together with ground truth information (top), only cells residing at spots being annotated as non-benign are shown. Blue indicates a deletion event while red indicates an amplification event. The ground truth shows the genomic profiles for all cells contributing to the spots assigned to a given clone. Comparing the inferred state with the ground truth on a clone

level, the average accuracy across genes was 0.90 (standard deviation 0.10) **(C)** Spatial organization of the synthetic data analysed in (B), with thumbnail of the complete cell population in the artificial tissue, each pixel corresponding to a cell. The cells' intensity levels are proportional to their total number of associated copy number events. Circles represent the spots used to "capture" transcripts. Spots are coloured by their inferred clone identity. Note how Clone 2, predicted to have zero copy number events, is found along the borders of both foci, where there's a mixture of benign and non-benign cells.

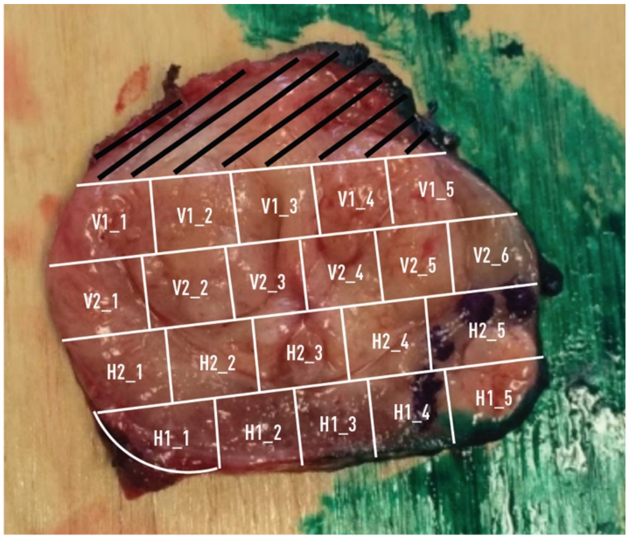


**Fig. S5. Demarcation of cubes from whole prostate cross-section.**
Image showing cut location from patient 1

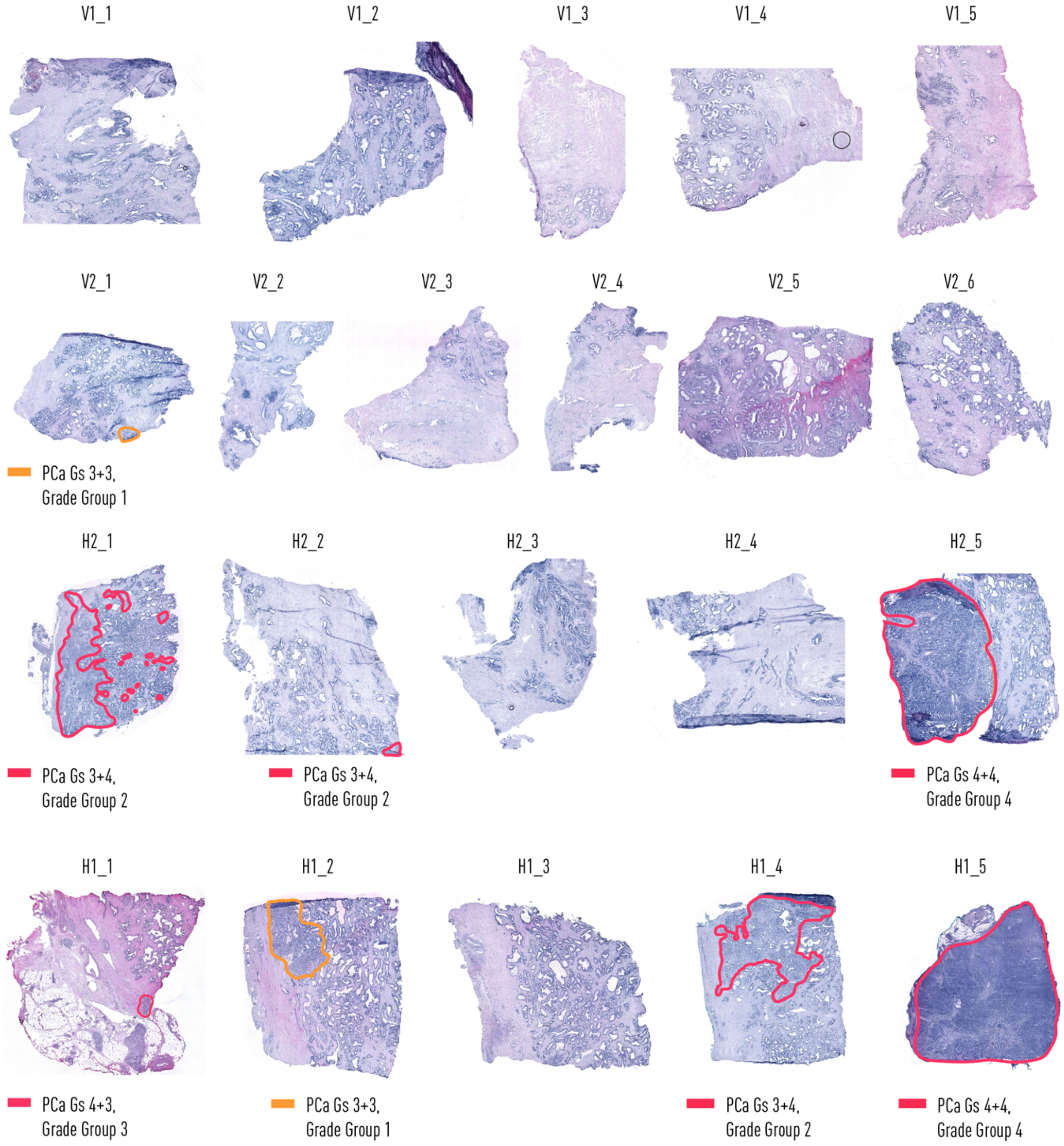


**Fig. S6. Prostate patient 1 histology**
Histology annotated by pathologist on sections for Spatial Transcriptomics (1k-arrays).

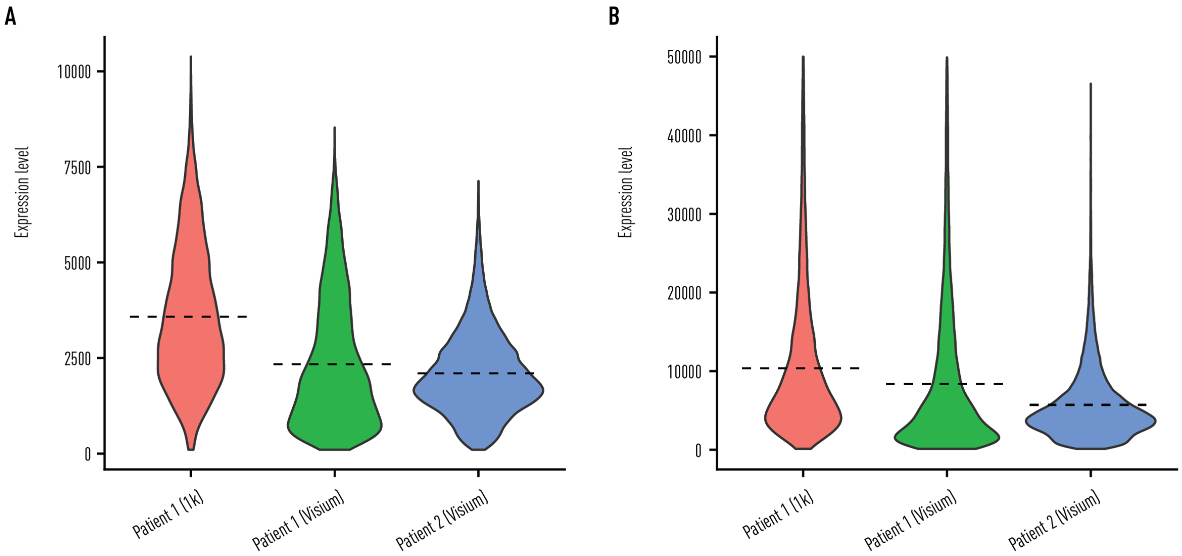


**Fig. S7. Quality control features for prostate spatial transcriptomics.**
(**A)** Quality control measures as violin plots with unique genes per spot and **(B)** unique transcripts per spot for all sections combined for each method and patient. Dotted lines represent the mean number of unique genes and transcripts per spot for each group: 3582, 2334 and 2104 for (A) and 10734, 10221 and 5711 for (B).

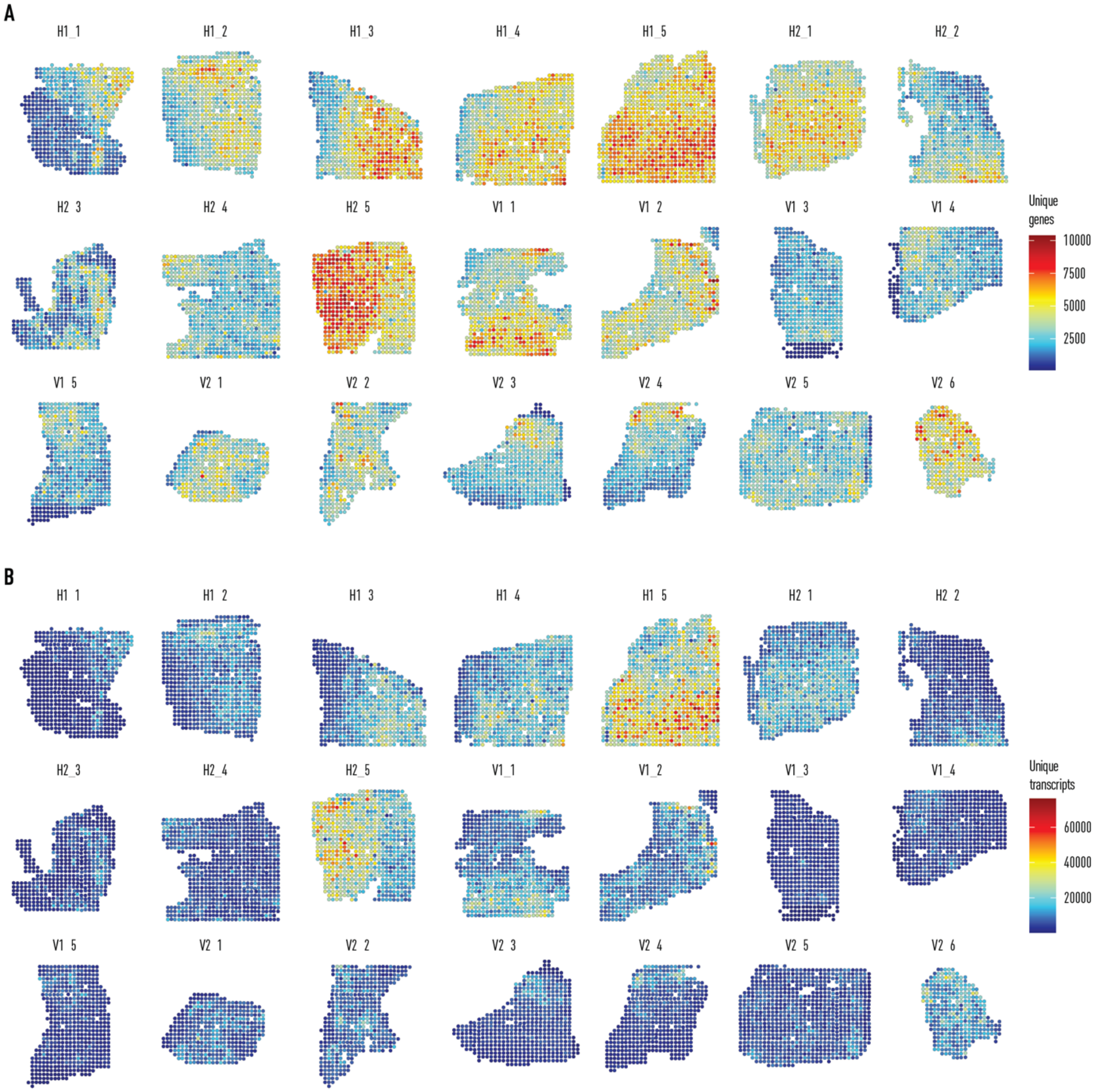


**Fig. S8. Quality control features per spot visualized spatially for whole organ spatial transcriptomics of patient 1 on 1k spatial transcriptomic arrays.**
(A) Unique genes per spot. (B) unique transcripts per spot.

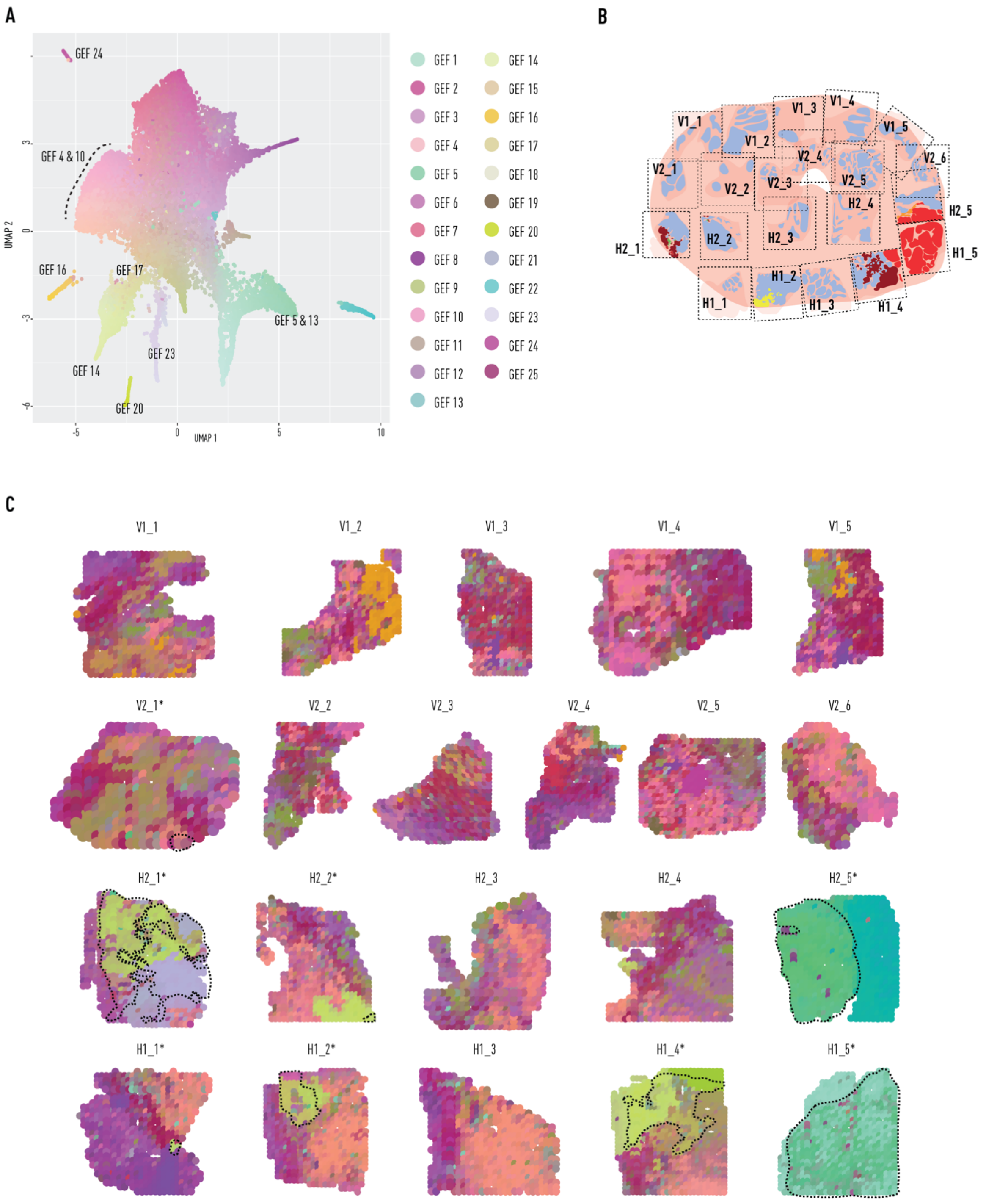


**Fig. S9. Organ wide analysis of prostate from patient 1 using 1k spatial transcriptomics arrays**.

**(A)** UMAP summary of GEFs. Marker genes for each factor available in table S1. **(B)** Cartoon diagram representing all tissue sections assessed through spatial transcriptomics from prostate patient 1 and their location in the organ. **(C)** UMAP projections overlaid onto tissue sections. Similar colours represent similar gene expression patterns, * mark sections annotated with cancer; dotted lines show cancer areas.

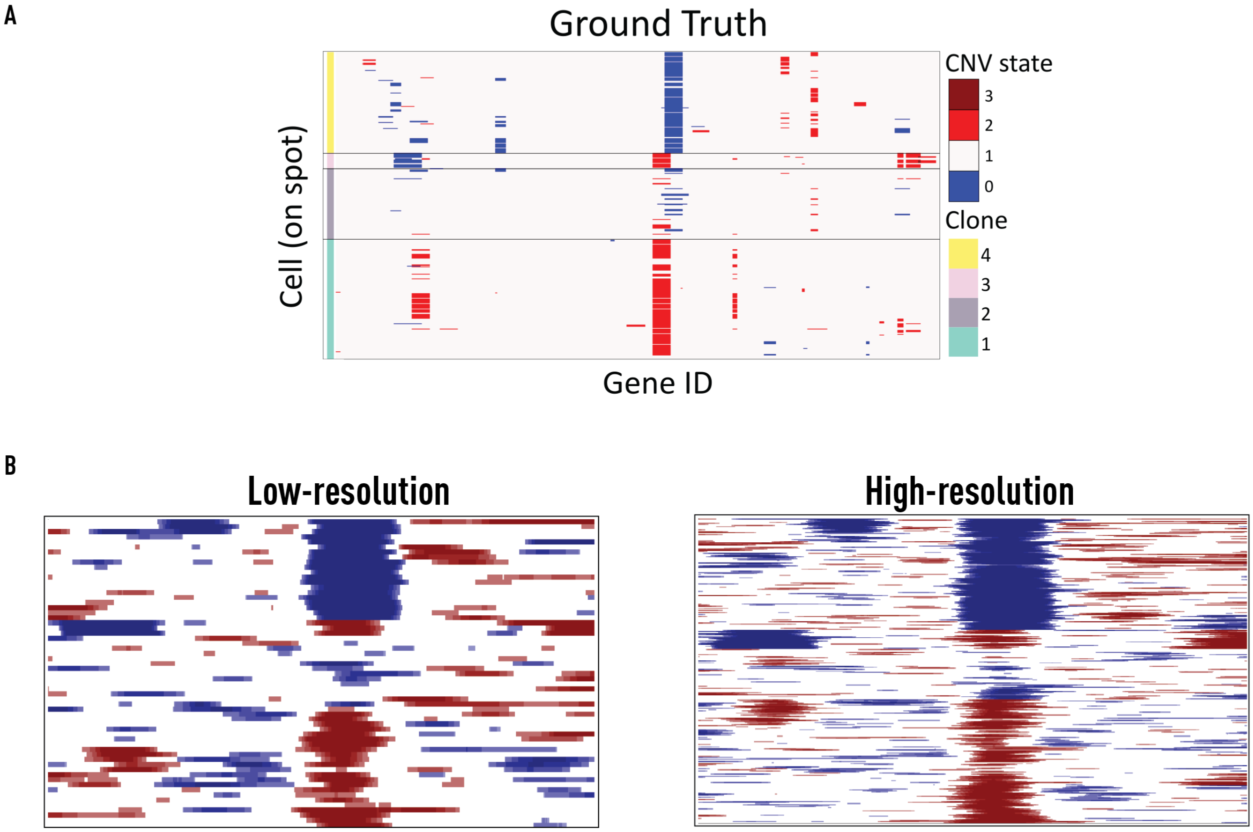


**Fig. S10. Low vs high resolution iCNV output from synthetic data.**
**(A)** Ground truth (from fig. S4). **(B)** iCNV outputs from simulated synthetic data of spots simulating ST 1k array (low-resolution) and Visium (high-resolution). High resolution spots were 0.55x size of low resolution and had 5x more spots per area. The synthetic ground truth data were identical for both.

iCNV = inferred Copy-Number Variant

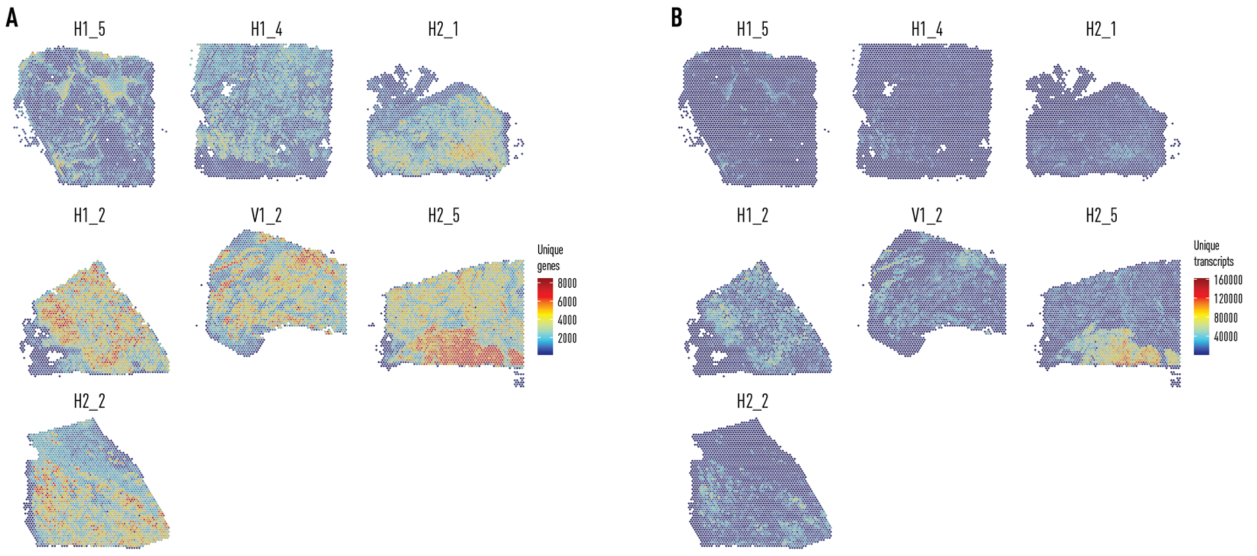


**Fig. S11. Quality control features per spot visualized spatially for high resolution spatial transcriptomics (Visium) of patient 1.**
**(A)** Unique genes per spot. **(B)** unique transcripts per spot.

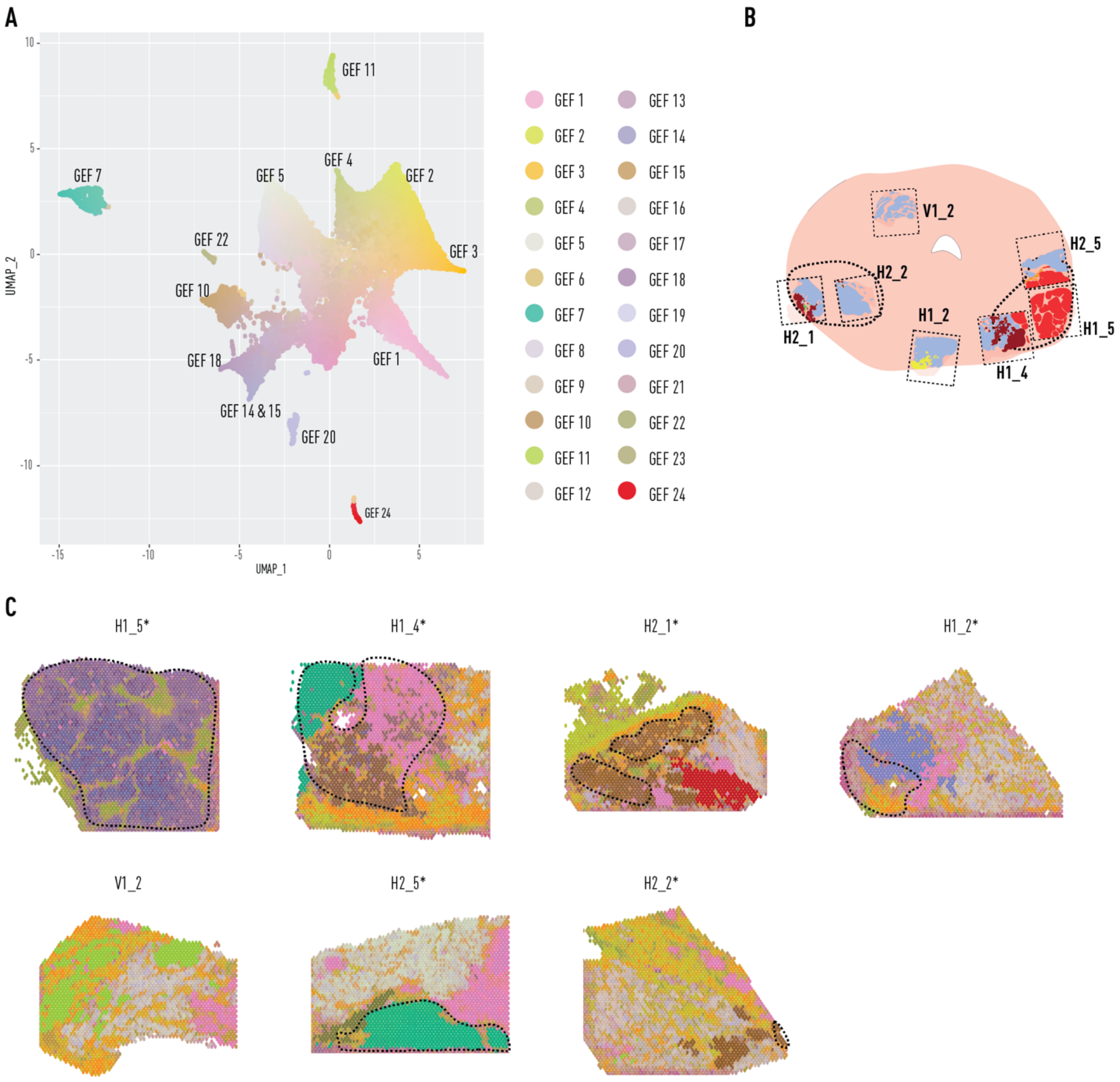


**Fig. S12. High resolution spatial transcriptomics (Visium) analysis of prostate tumour regions from patient 1.**

**(A)** UMAP summary of GEFs. Marker genes for each factor available in table S2. **(B)** Cartoon diagram representing tissue sections assessed through high resolution spatial transcriptomics from prostate patient 1 and their location in the organ. **(C)** UMAP projections overlaid onto tissue sections. Similar colours represent similar gene expression patterns, * mark sections annotated with cancer; dotted lines show cancer areas.

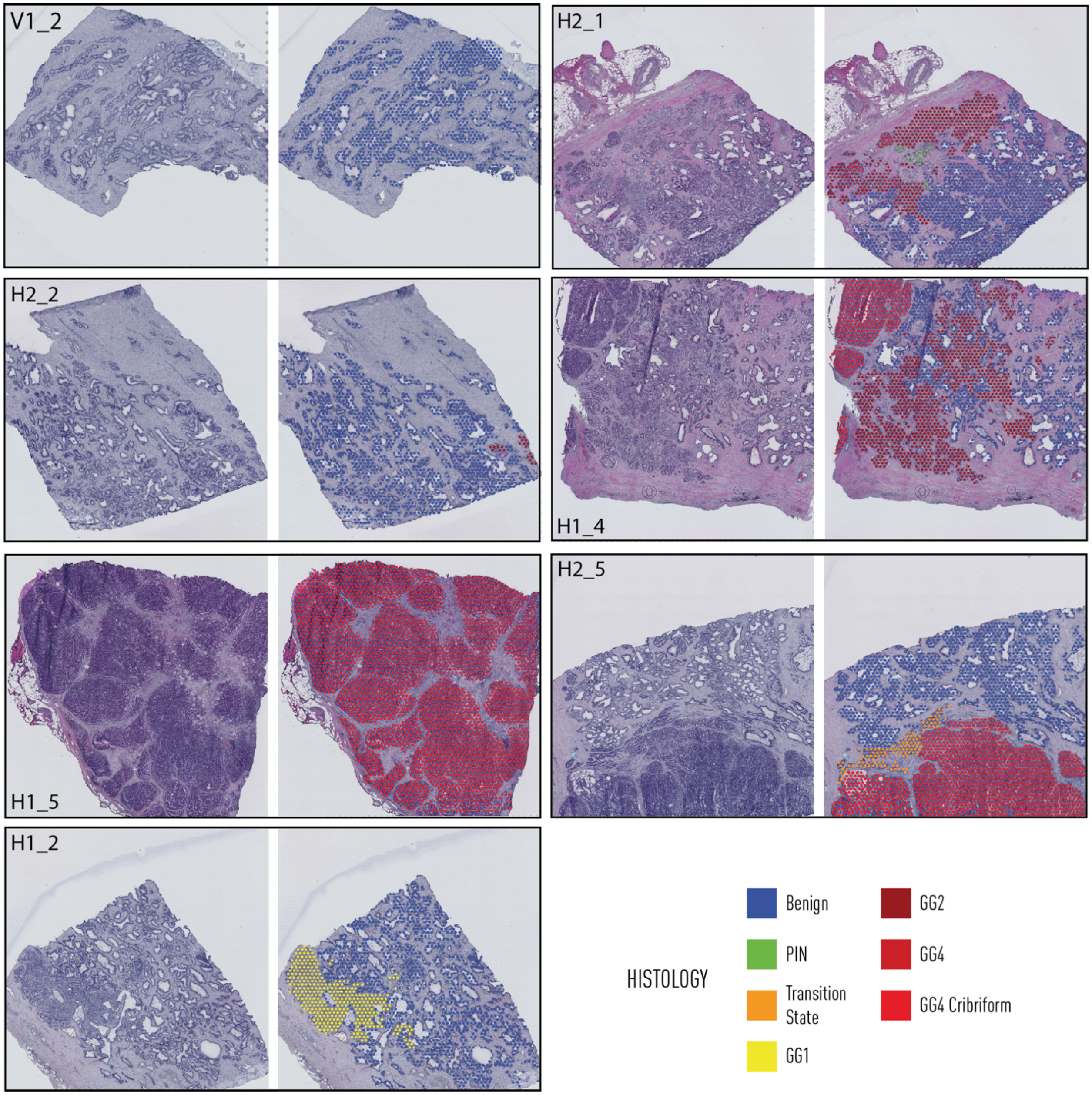


**Fig. S13. Prostate patient 1 histology.**
Spot-level consensus annotation for all Visium run sections from prostate of patient 1.

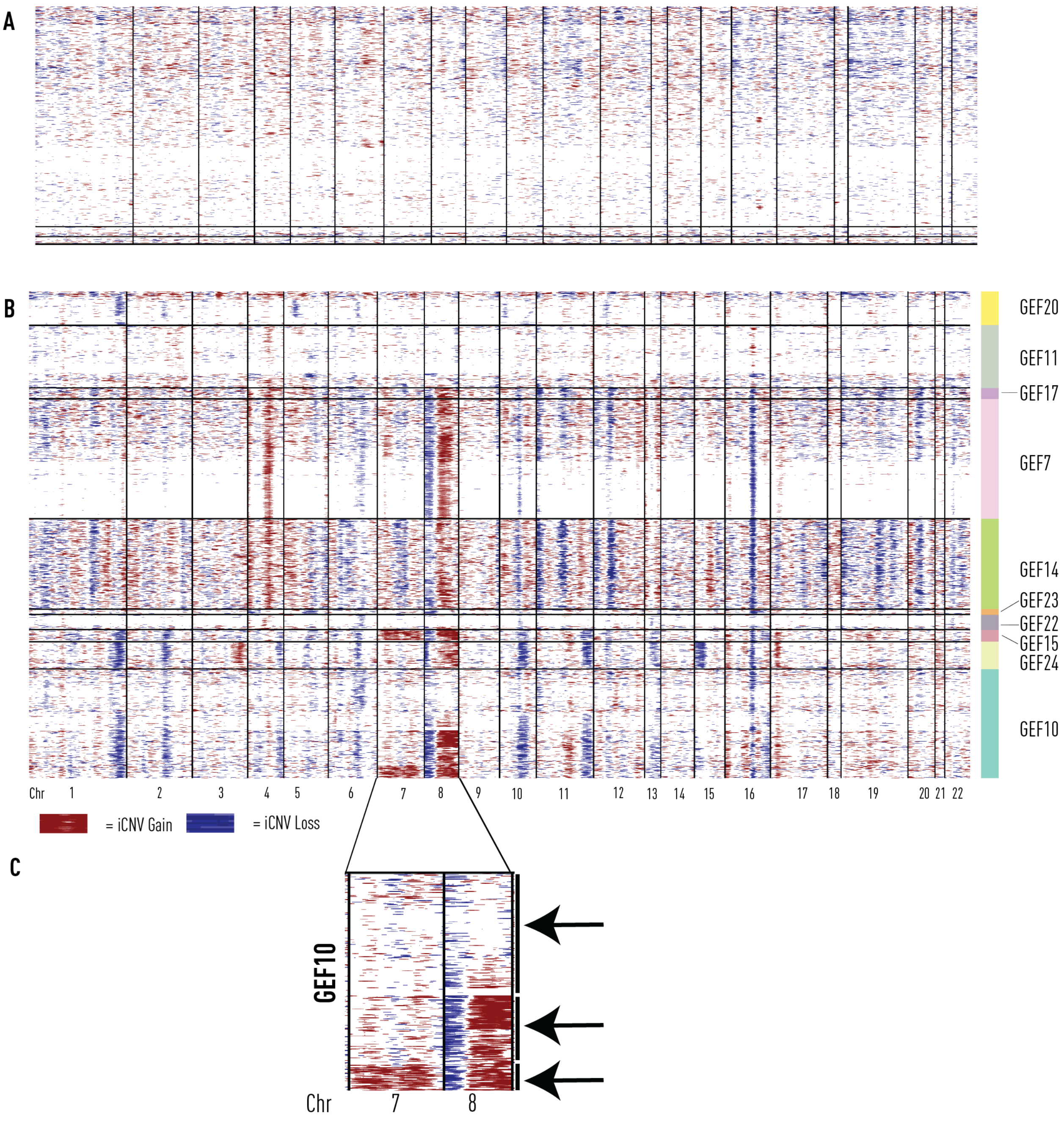


**Fig. S14. GEF directed iCNV analysis from prostate patient 1 (high resolution Visium analysis).**

**(A)** Benign GEFs (GEF5, GEF9, GEF12, GEF13, GEF16) were used as a reference set for analysis of **(B)** Tumour GEFS. **(C)** Snapshot of iCNV profiles for chr 7 and 8 from GEF10. GEF iCNV heterogeneity is highlighted by 3 subclones: the first harbouring no changes to chr 7 and 8, the second having a deletion and amplification in chr 8, and the last having the alterations in chr 8 as well as an amplification of chr 7.

GEF = Gene Expression Factor, chr = Chromosome, iCNV = inferred Copy-Number Variant.

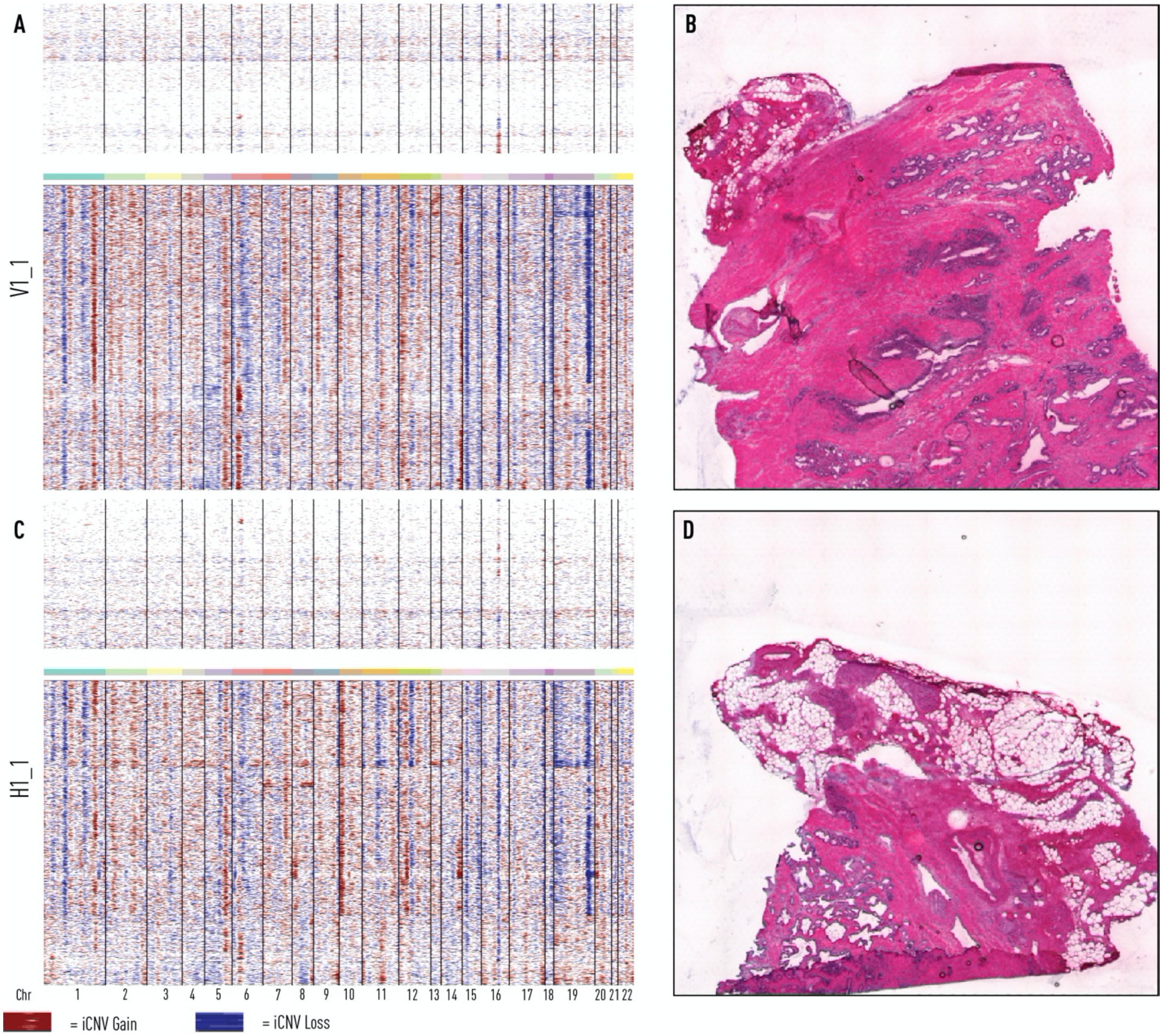


**Fig. S15. Somatic events in prostate tissue sections from prostate patient 1.**

**(A)** iCNV profile for section V1_1. **(B)** Histological image for prostate patient 1, section V1_1. **(C)** iCNV profile for section H1_1. **(D)** Histological image for prostate patient 1, section H1_1.

iCNV = inferred Copy-Number Variant.

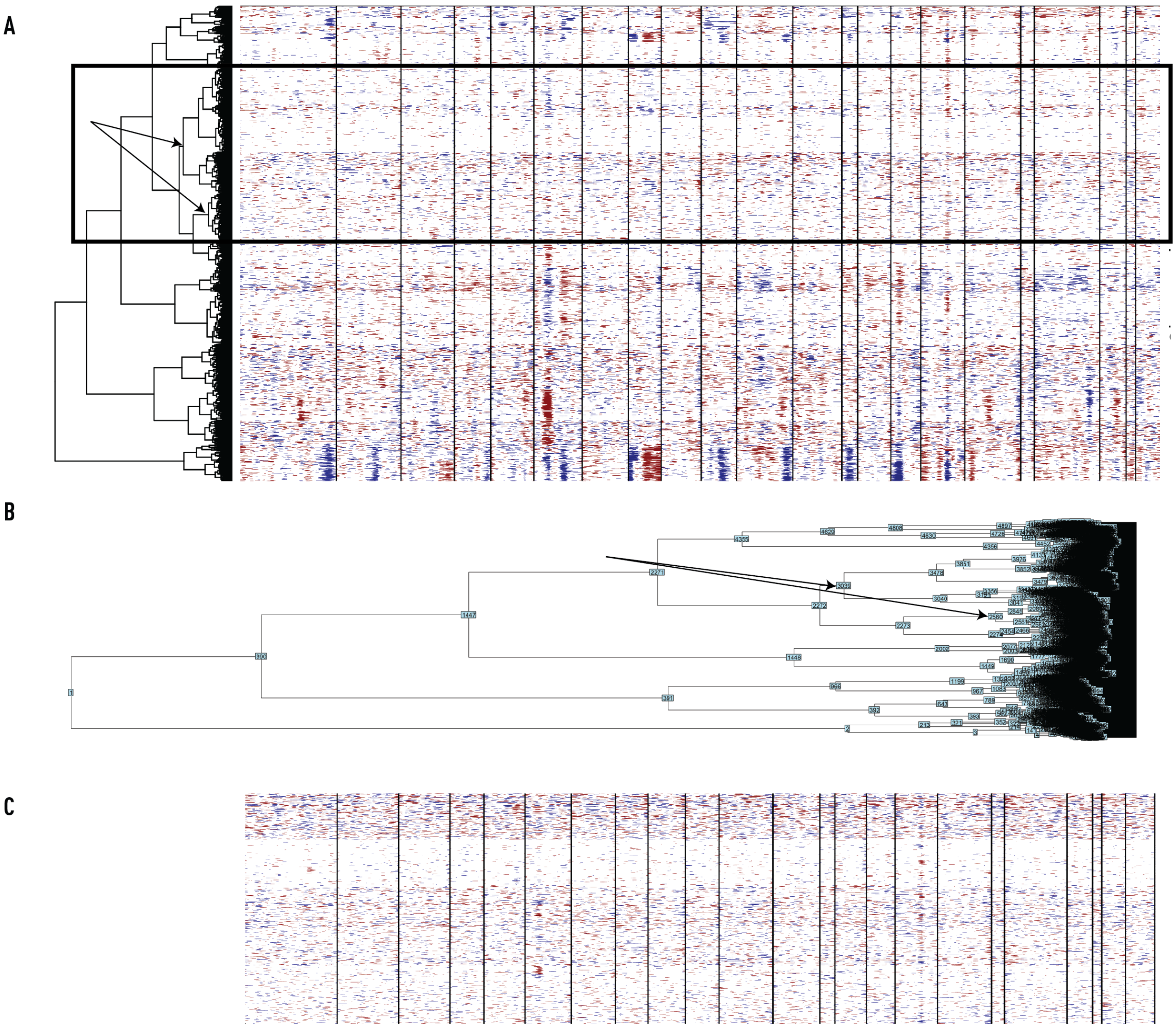


**Fig. S16. Identification of a histologically benign reference set.**

**(A)** Visual selection of benign epithelial spots harbouring the least amount of inferred copy number variations (iCNV), as outlined by the black box bounding box. Arrows identify dendrogram nodes corresponding to barcoded spots within the box. **(B)** SpatialInferCNV output of the dendrogram nodes with numerical identifiers for selection corresponding to Panel A. **(C)** Finalized benign reference set from analysis of epithelial cells in prostate patient 1, section H2_1 (Figure 3).

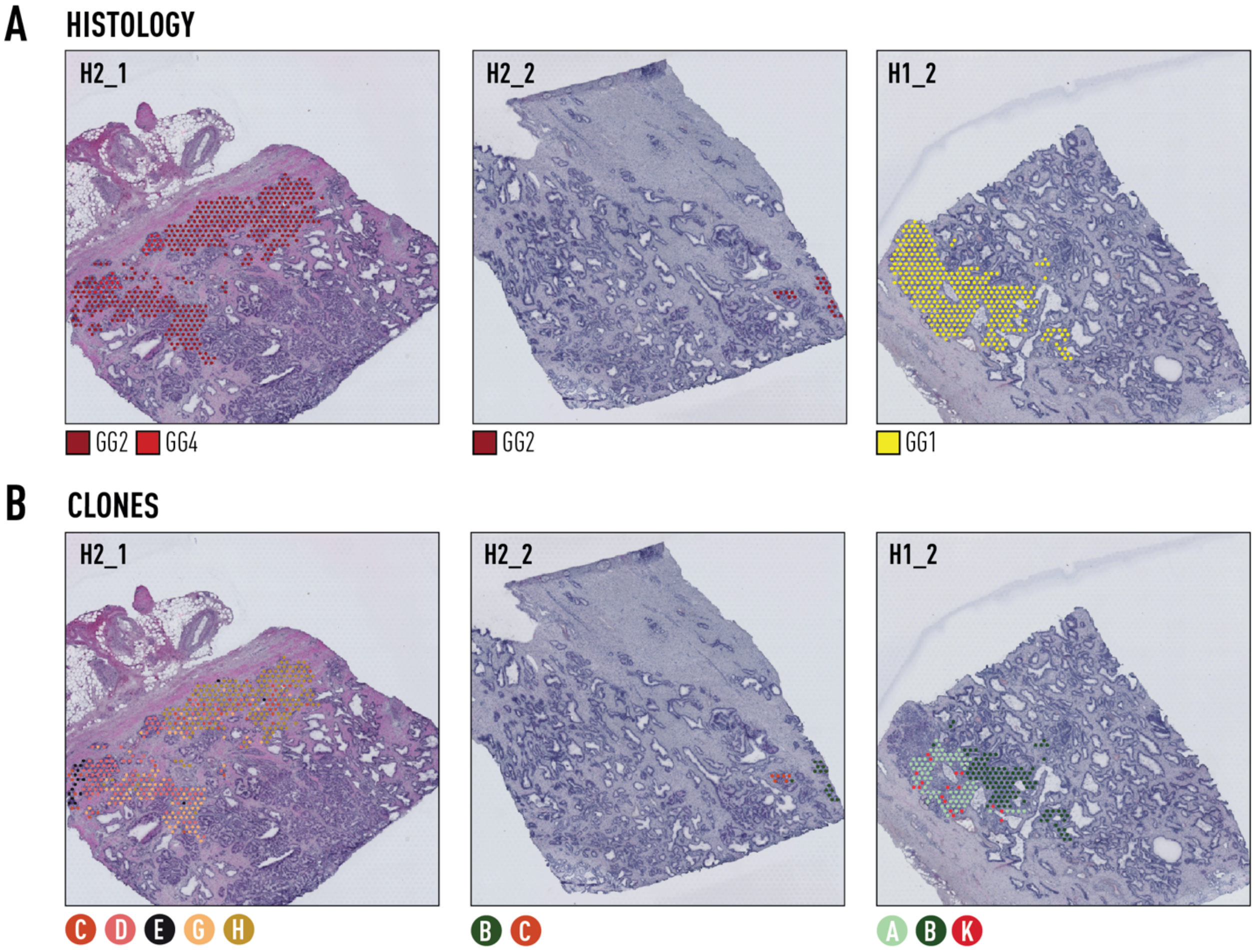


**Fig. S17. Histology and clones (Figure 2A) from prostate patient 1.**
**(A)** Consensus pathology annotations for tumour spots from sections H2_1, H2_2, and H1_2. **(B)** Clonal groupings of spots (approx. 10-15 cells each) determined by hierarchical clustering (Supplementary Material).

GG=ISUP Gleason 'Grade Group'

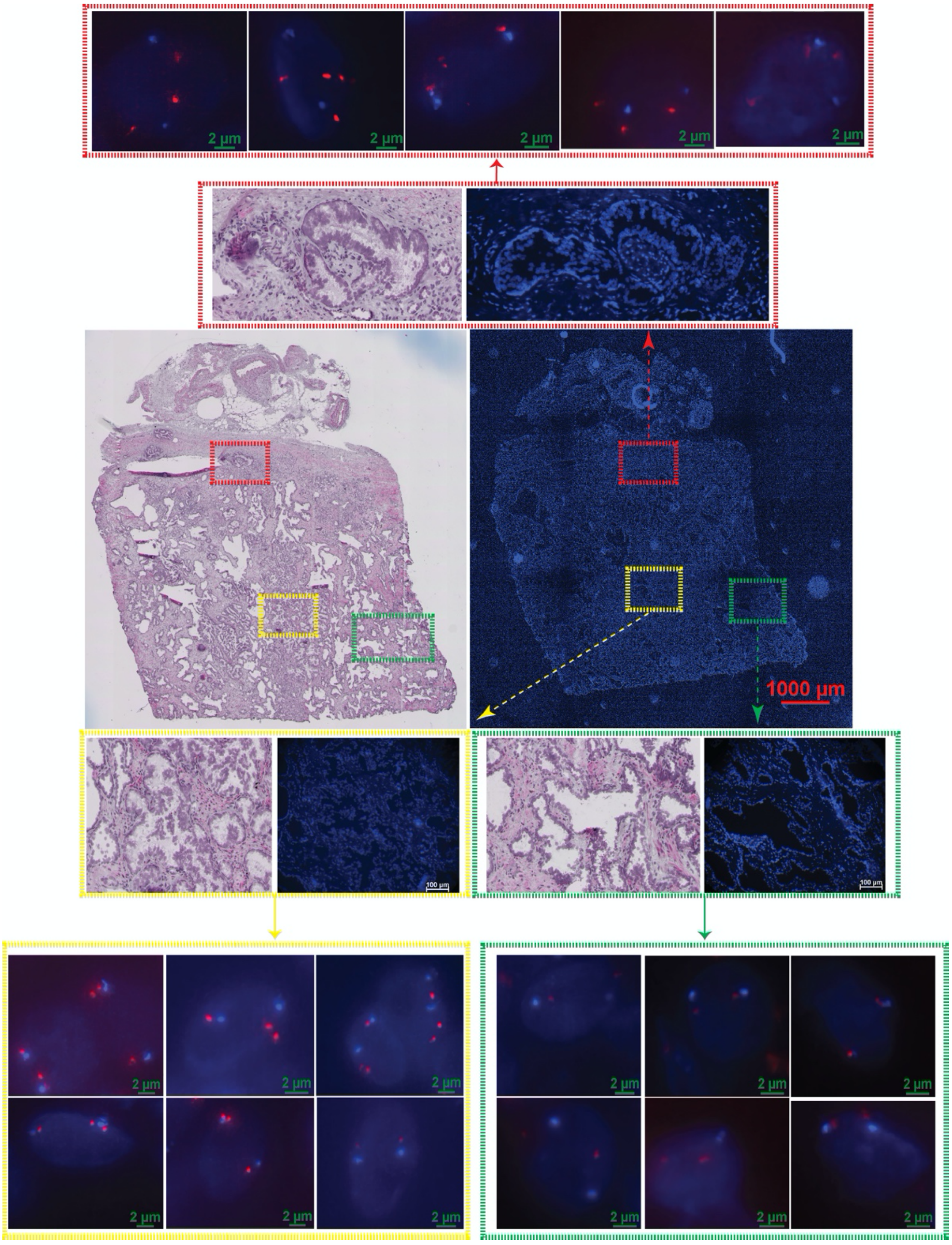


**Fig. S18. DNA FISH targeting MYC gene.**
Representative DNA FISH images of fresh frozen prostate tissue sections obtained from patient 1 section H2.1 labelled with Cytocell *MYC*/8cen amplification probes. Coloured-dotted boxes on the zoom-out whole section scan represent tumour (red), benign (yellow), benign (green) regions that are further zoomed-in to single cell resolution respectively for visualization purpose. *MYC* FISH signal is shown in (**red**), chr 8 centromere as internal control in (**aqua**), and nuclei are counterstained with DAPI in (**dark blue**). Consecutive sections were used for H&E staining and FISH. Regions of interest confirmed as epithelial by histo-pathologist from adjacent H&E-stained sections.

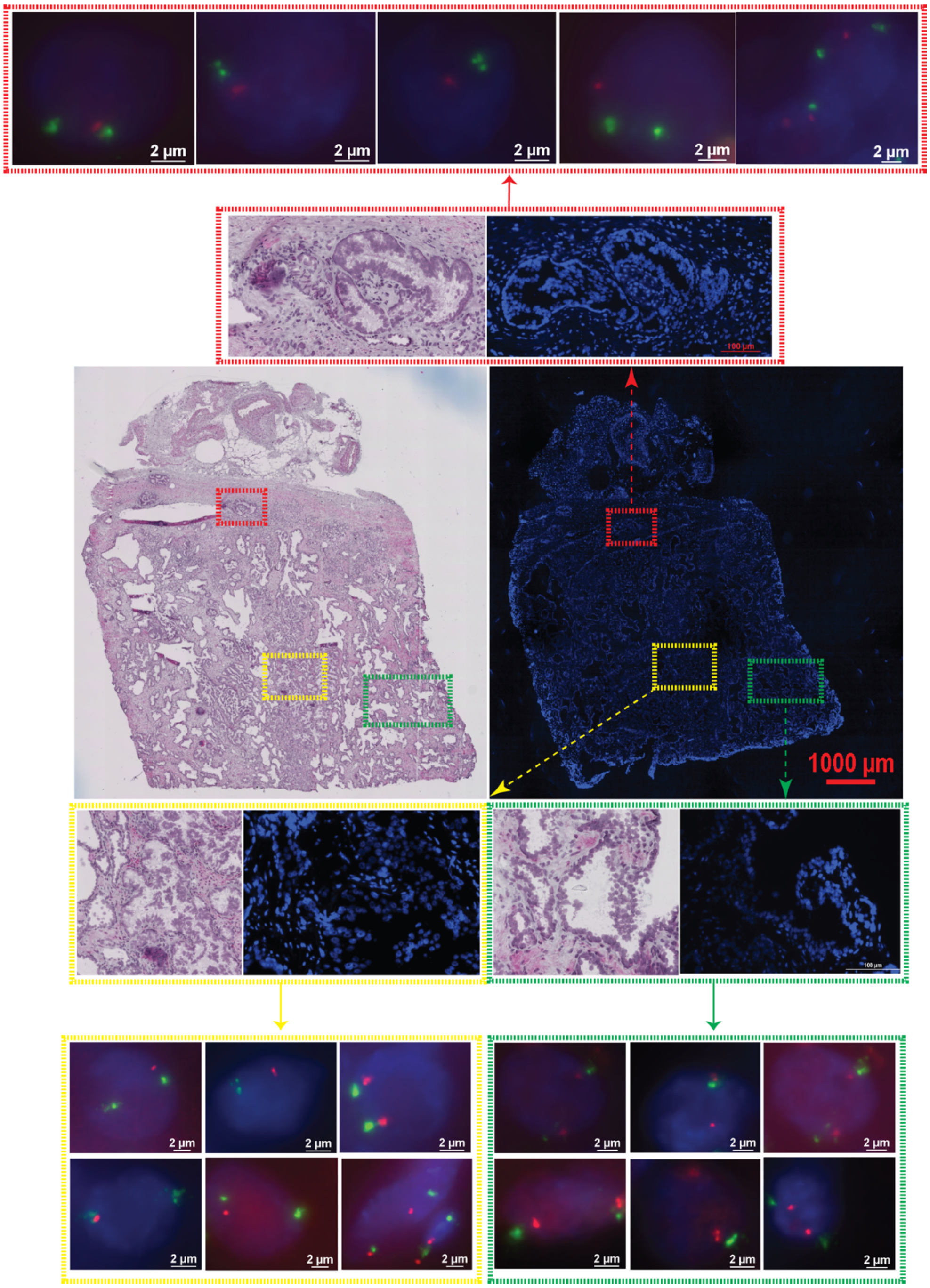


**Fig. S19. DNA FISH targeting PTEN gene.**
Representative DNA FISH images of fresh frozen prostate tissue sections obtained from patient 1 section H2.1 labelled with Cytocell *PTEN*/10cen deletion probes. Coloured-dotted boxes on the zoom-out whole section scan represent tumour (red), benign (yellow), benign (green) regions that are further zoomed-in to single cell resolution respectively for visualization purpose. *PTEN* FISH signal is shown in (**red**), chromosome 10 centromere as internal control in (**green**), and nuclei are counterstained with DAPI in (**dark blue**). Consecutive sections were used for H&E staining and FISH. Regions of interest confirmed as epithelial by histo-pathologist from adjacent H&E-stained sections.

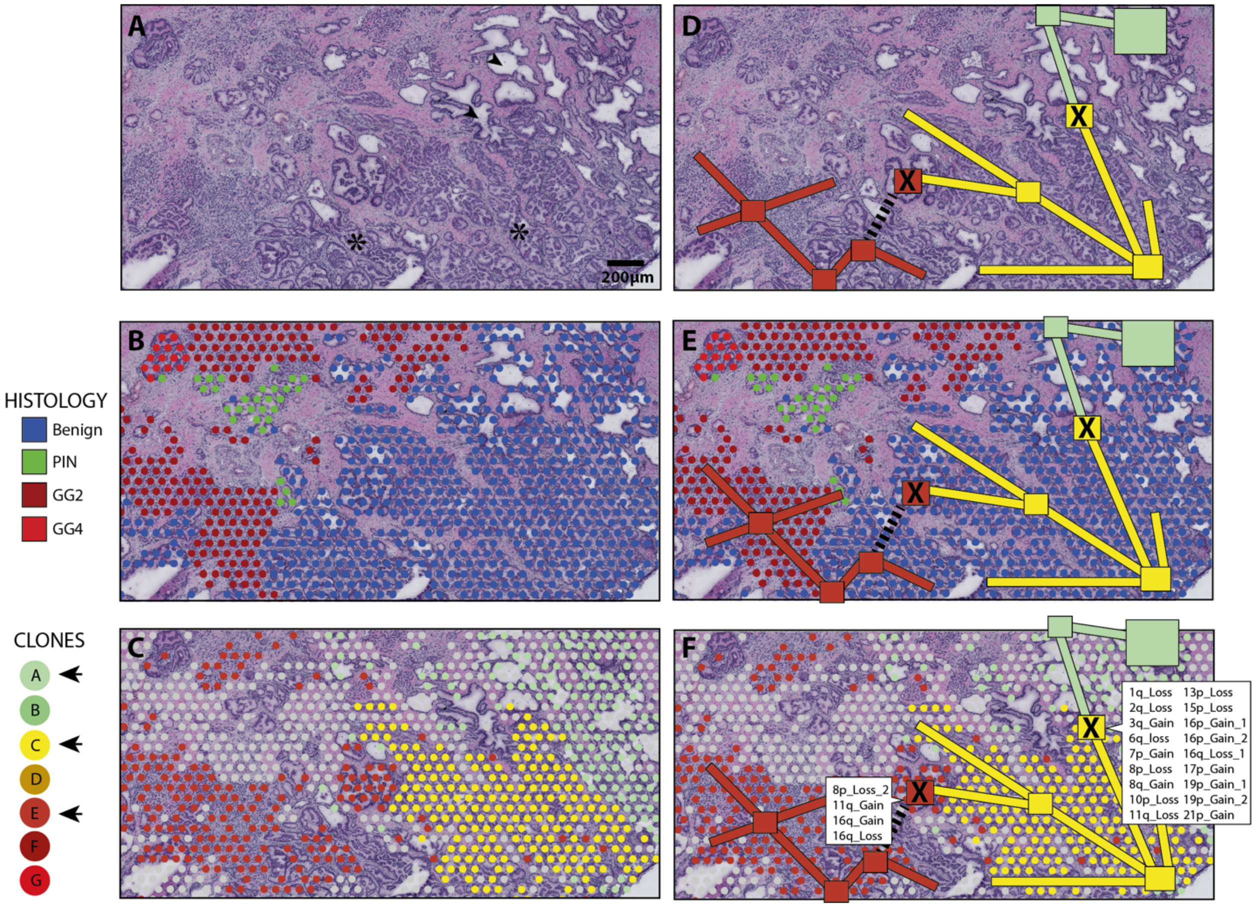


**Fig. S20. Branching morphogenesis and somatic mosaicism in prostate epithelium.**
**(A)** Close up histology of section H2_1 demonstrating clear ductal (e.g. arrow heads) and acinar (e.g. stars) branching patterns. **(B)** Overlayed spot-level histology. **(C)** Overlayed clone groupings (from Figure 3). **(D-F)** Possible arrangement of clonal expansion during branching morphogenesis with key mutational events (marked with X, iCNV events from Figure 3, Supp Table) passed on to downstream branches. Dotted line represents presumed branch/duct not visible in two-dimensional plane.

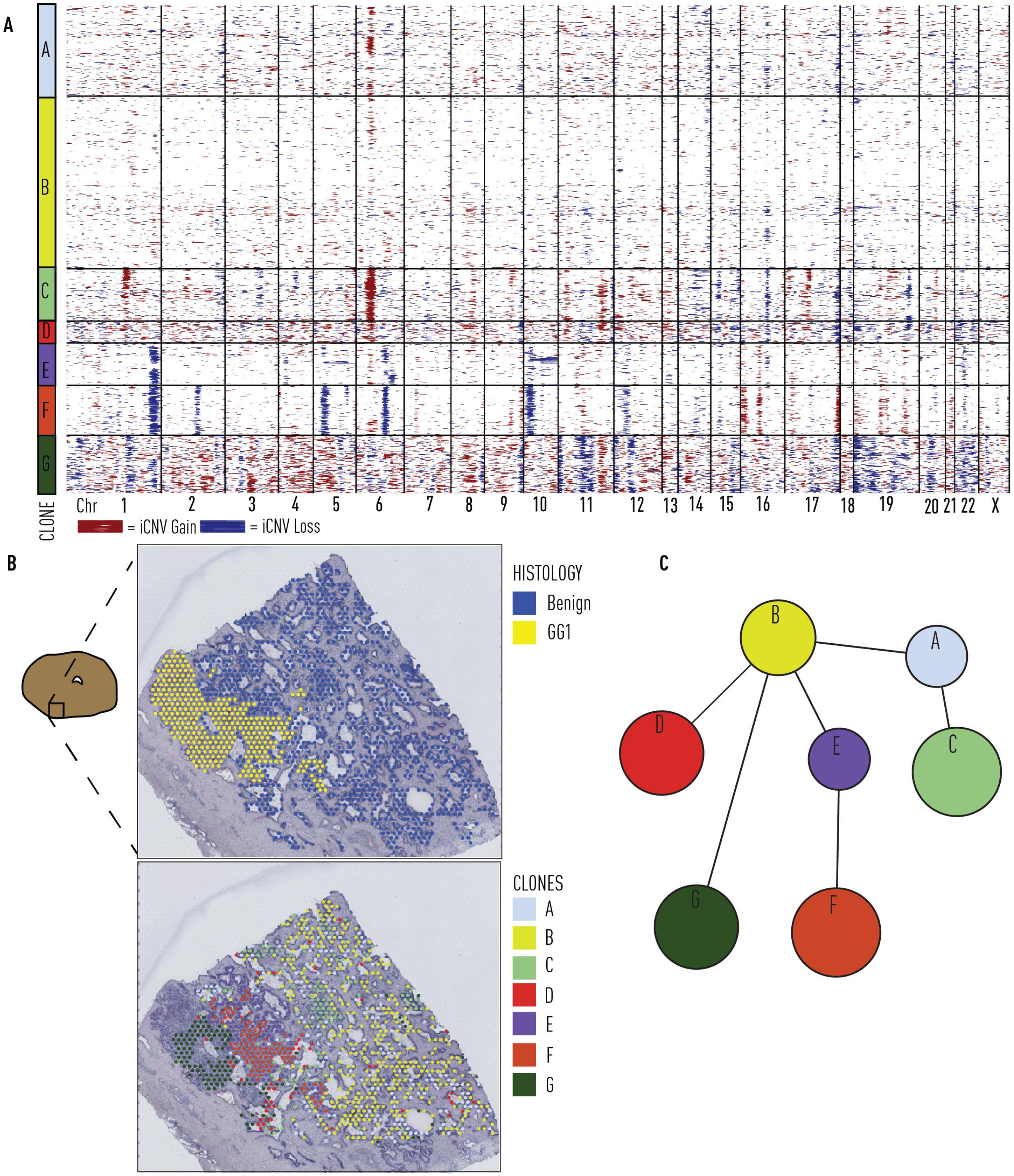


**Fig. S21. Distinct iCNV profile of GG1 tumour focus from organ scale prostate patient 1.** **(A)** iCNV profiling of epithelial Visium spots from section H1_2. **(B)** Spot level histology and iCNV clone calls. **(C)** iCNV clone tree.

iCNV = inferred copy number variant, GG1 = Gleason Grade Group 1

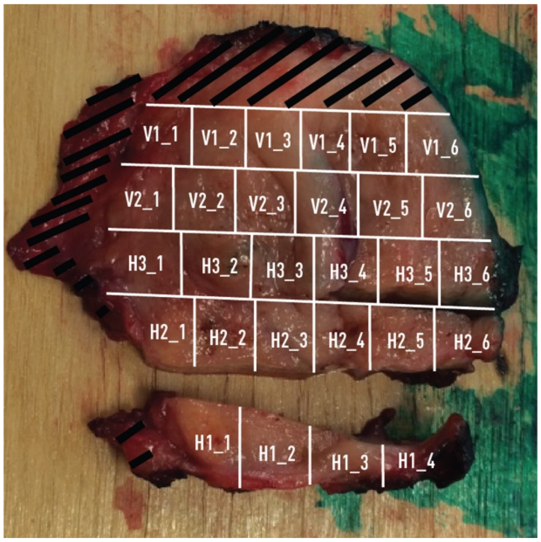


**Fig. S22. Demarcation of cubes from whole prostate cross-section.**
Image showing cut location from patient 2

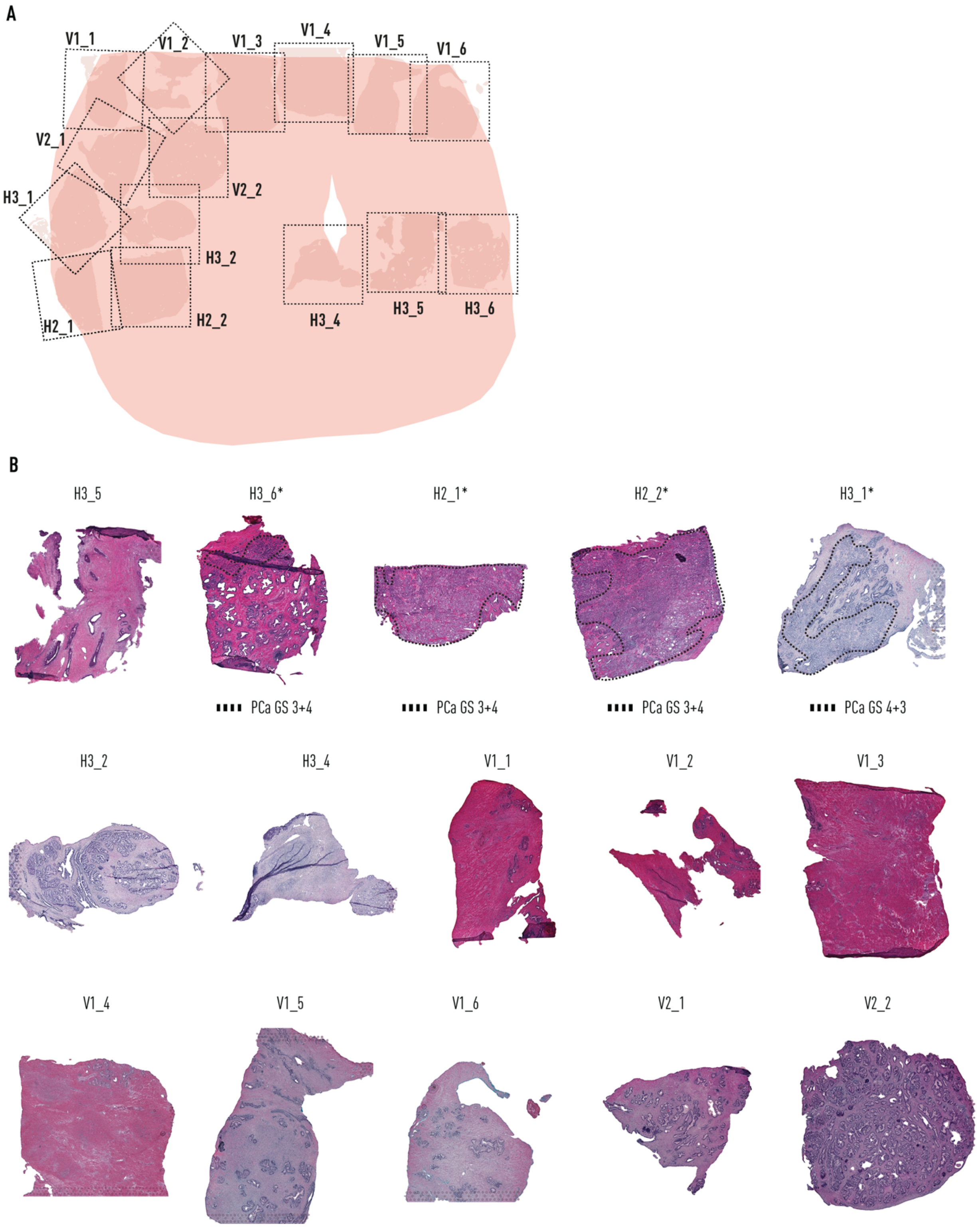


**Fig. S23. Histology of tissue sections of prostate sections from patient 2.**

**(A)** Cartoon diagram representing all tissue sections assessed through high resolution spatial transcriptomics (Visium) from prostate patient 2 and their location in the organ. **(B)** H&E-stained images of sections. * mark sections containing cancer.

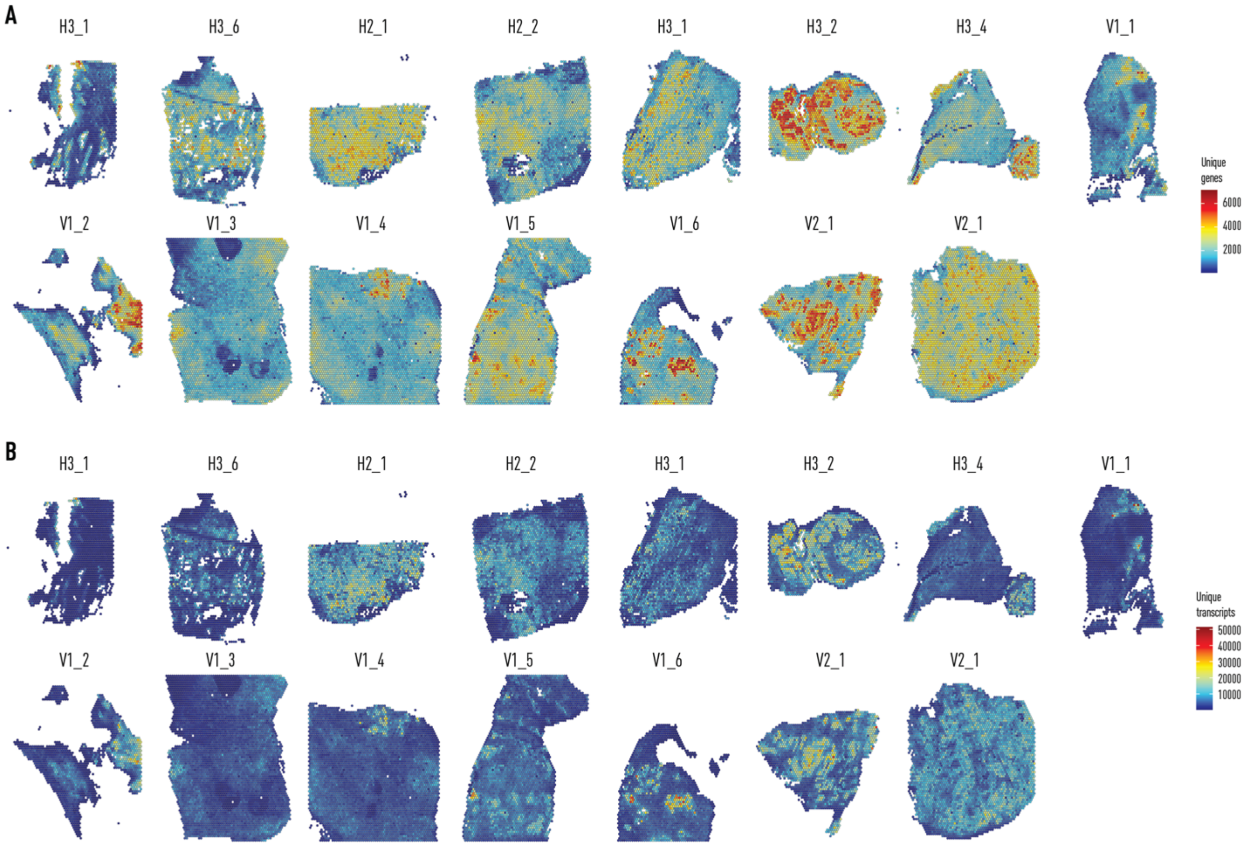


**Fig. S24. Quality control features per spot visualized spatially for high resolution spatial transcriptomics (Visium) of patient 2.**
**(A)** Unique genes per spot. **(B)** unique transcripts per spot.

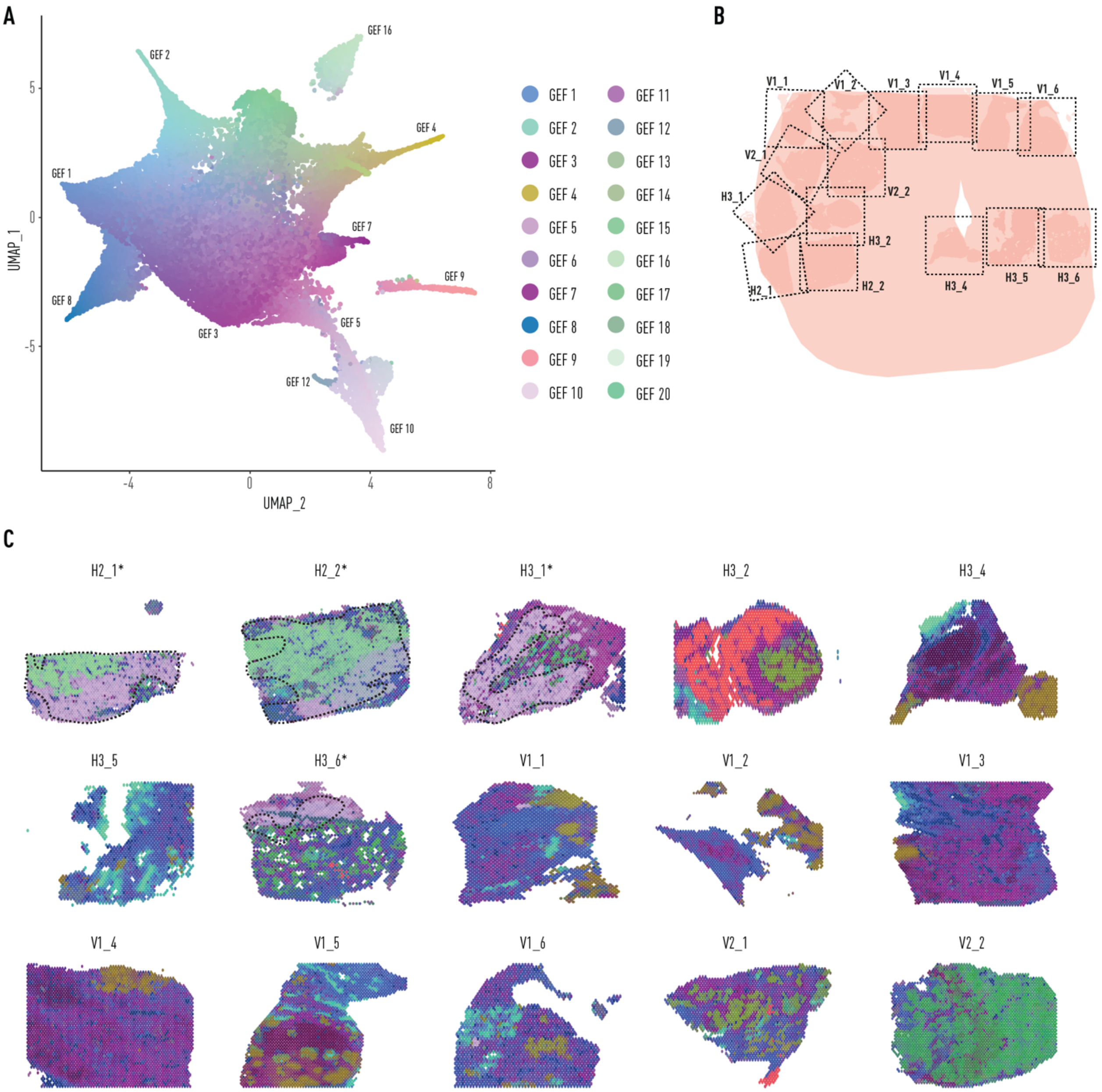


**Fig. S25. High resolution spatial transcriptomics (Visium) analysis of prostate from patient 2.**

**(A)** UMAP summary of GEFs. Marker genes for each factor available in table S3. **(B)** Cartoon diagram representing tissue sections assessed through high resolution spatial transcriptomics from prostate patient 2 and their location in the organ. **(C)** UMAP projections overlaid onto tissue sections. Similar colours represent similar gene expression patterns, * mark sections annotated with cancer; dotted lines show cancer areas.

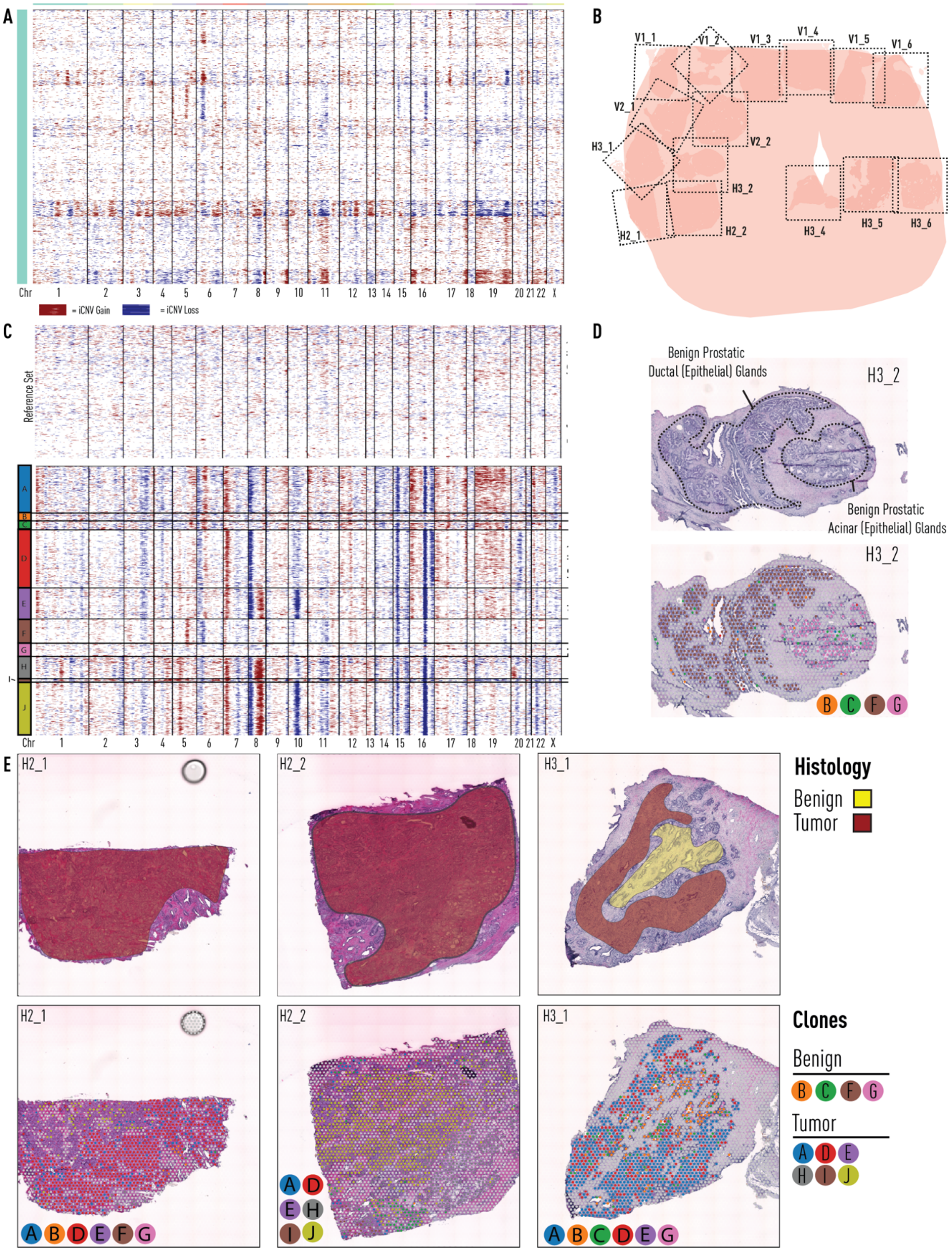


**Fig. S26. Organ scale prostate patient 2.**
**(A)** iCNV profiles of histologically benign prostatic epithelial cells from 11 sections from prostate patient 2. **(B)** Reference overview of 15 sections available for analysis: sections H2_1, H2_2, H3_1, and H3_6 harbour tumour. Black dotted lines represent the area covered by spatial transcriptomics array surface. **(C)** Analysis of tumour foci in sections H3.1, H2.1 and H2.2.

Analysis includes section H3.2, a non-tumour bearing section which included spatially co-localized benign spots harbouring iCNV alterations from panel A. **(D)** Spatial histology and clone distribution in section H3.2 (no-tumour). Benign ductal histology (Clone F) harbours distinct iCNVs (chr 5 amplification, chr6 deletion), not harboured in neighbouring benign acinar glands (Clone G). **(E)** Section histology (transparent red indicates tumour, and transparent yellow denotes benign) and clones from tumour-bearing sections H3.1, H2.1, and H2.2.

iCNV = inferred Copy-Number Variant, chr = chromosome

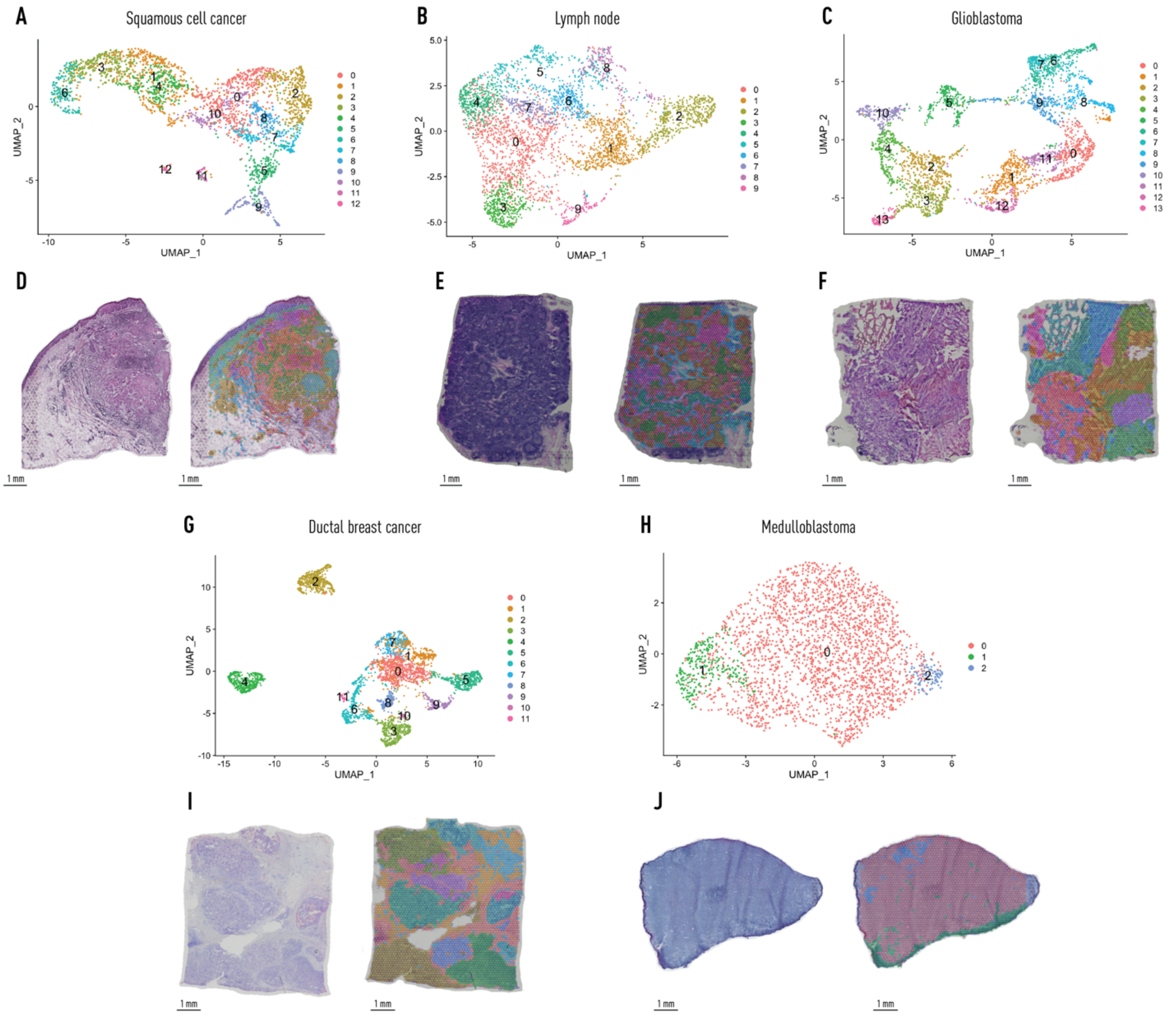


**Fig. S27. Spatial transcriptomics analysis of multiple sample types.**
(**A-C, G and H**) Transcript UMAPs of all spots labelled by cluster from human squamous cell carcinoma (A), human lymph node (B), human glioblastoma multiforme (C), human invasive ductal breast carcinoma (G), malignant childhood brain tumour diagnosed as medulloblastoma SHH grade IV (H). **(D-F, I and J)** H&E stain and unbiased cluster spots visualized spatially on tissue from human squamous cell carcinoma (D), human lymph node (E), human glioblastoma multiforme (F), human invasive ductal breast carcinoma (I), childhood medulloblastoma (J). Top gene markers for each cluster can be found in supplementary Table S4-S8.

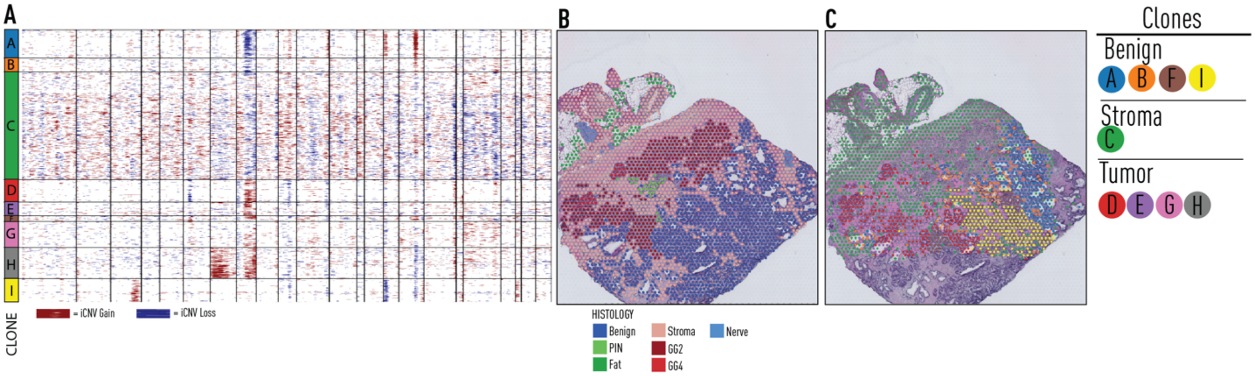


**Fig. S28. Repeat iCNV clone calls on prostate patient 1, section H2_1 without reference set.** (A) iCNV heatmap and clone calls. (B) Consensus spot-level histology. (C) Spatial mapping of clone calls.

iCNV = inferred Copy-Number Variant, PIN = Prostatic Intra-epithelial Neoplasia, GG = Gleason Grade Group.

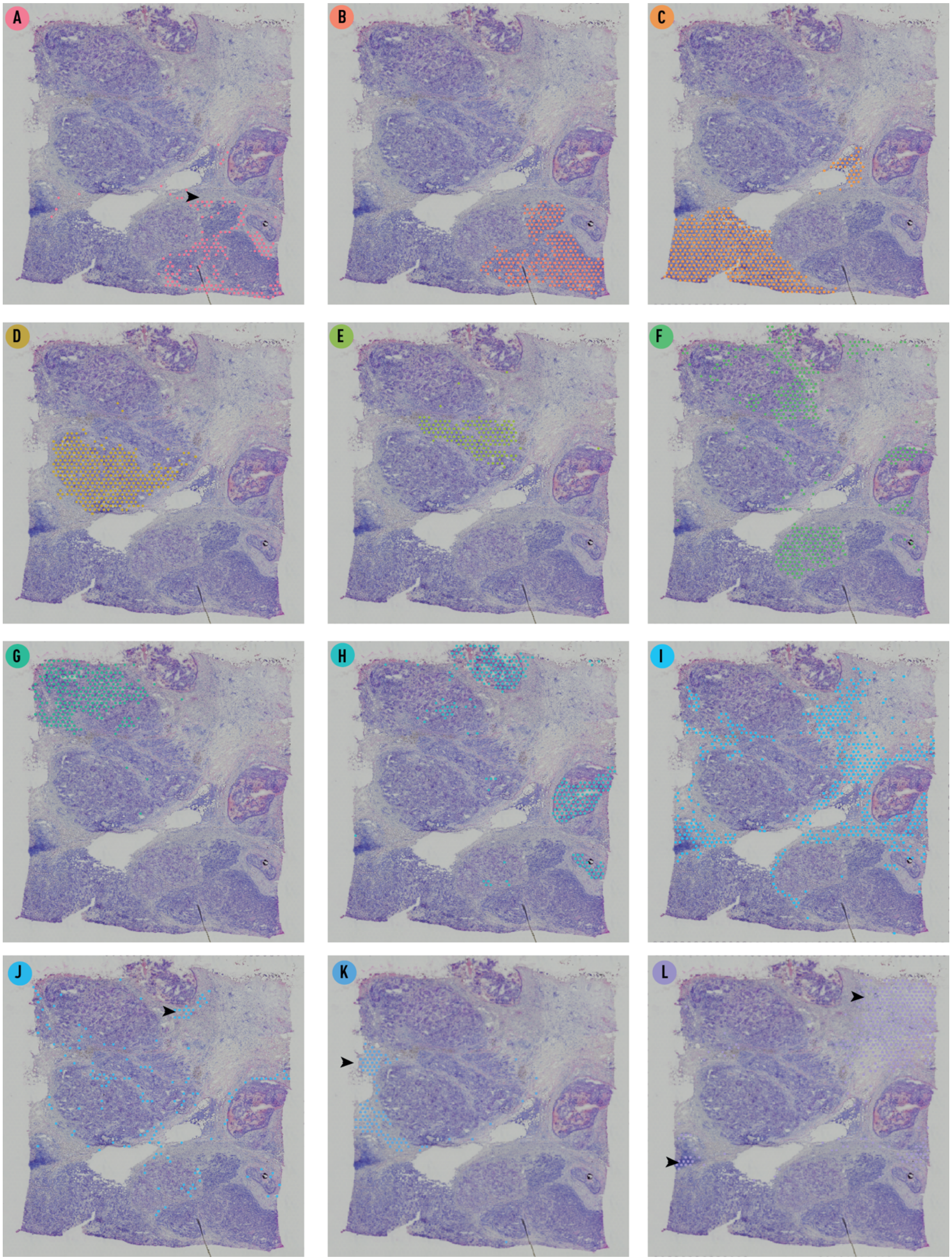


**Fig. S29. Clones identified by SpatialInferCNV for ductal breast carcinoma.**
(**A-J**) Clones A-J (Figure 4) visualized separately. Arrows added to aid reader in locating clones.

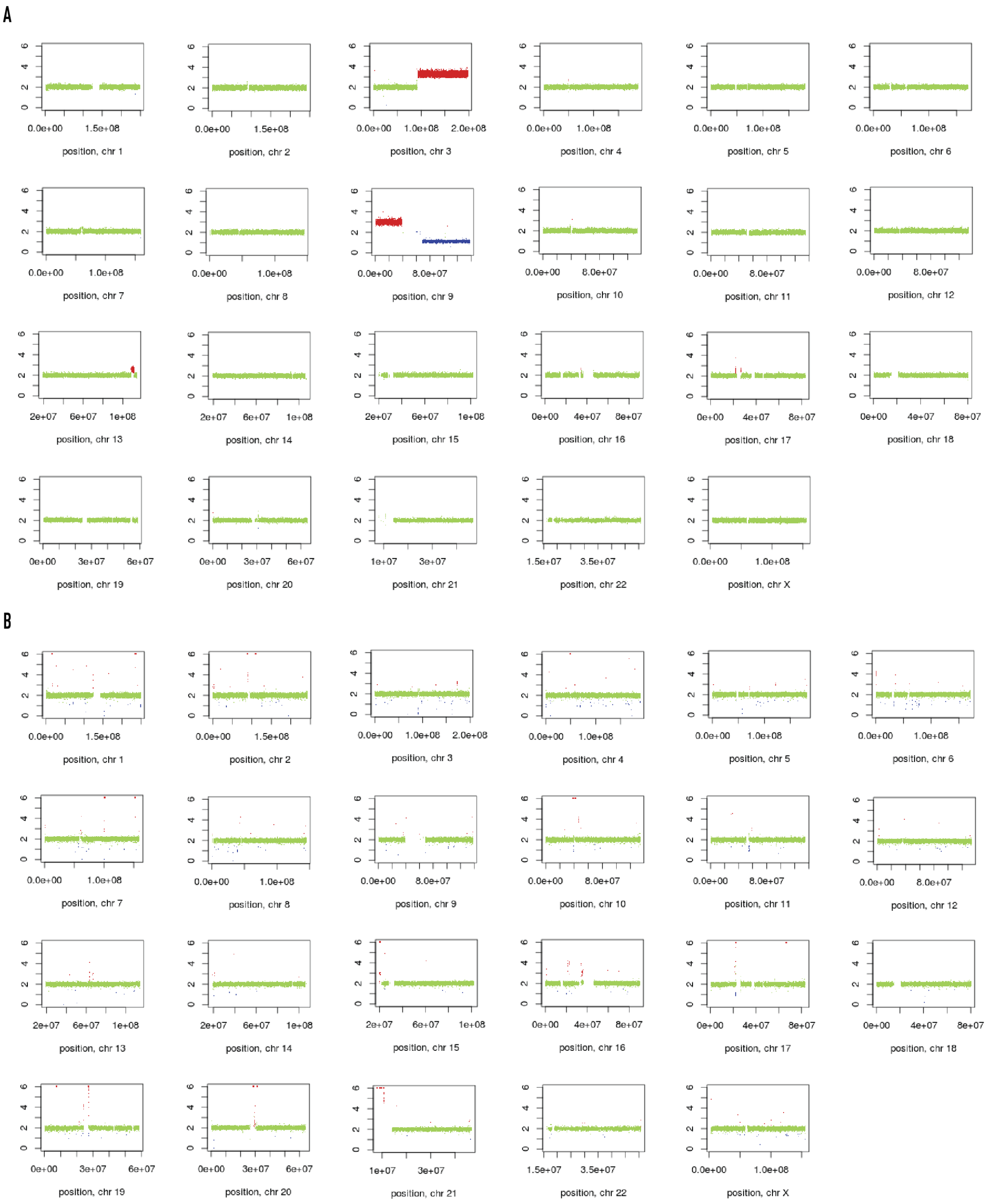


**Fig. S30. Whole-genome sequencing-based copy number profiles for paediatric brain tumour patients 1.**
**(A)** Somatic WGS CNV profile of patient diagnosed with medulloblastoma (grade IV, desmoplastic/nodular, SHH-activated) with **(B)** match normal blood.

WGS = Whole-genome sequencing. Chr = Chromosome. SHH = Sonic hedgehog. CNS = Central nervous system. NOS = Not otherwise specified.

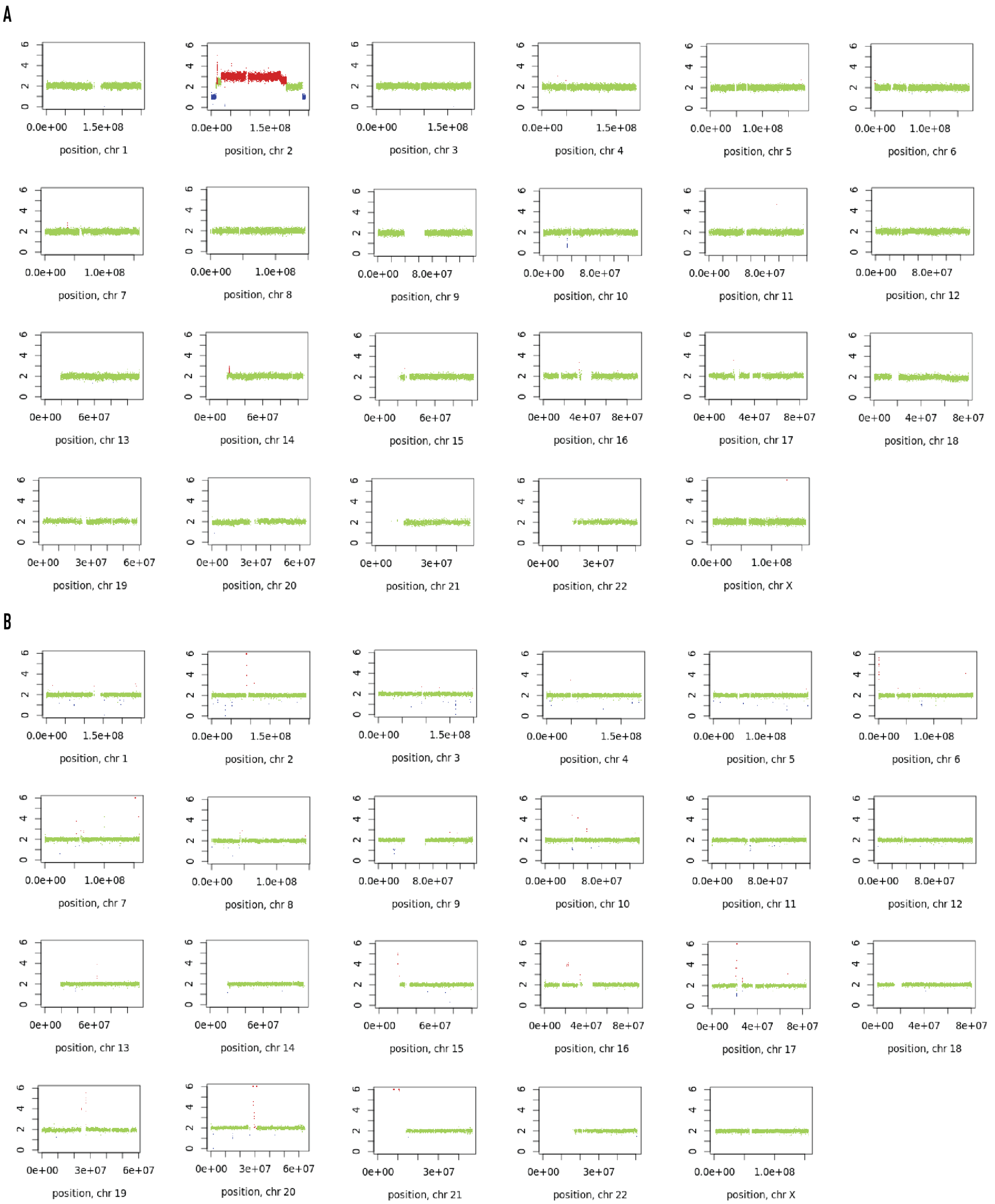


**Fig. S31. Whole-genome sequencing-based copy number profiles for paediatric brain tumour patients 2.**
**(A)** Somatic WGS CNV profile of patient diagnosed with medulloblastoma (grade IV, classic morphology, SHH-activated) with **(B)** match normal blood.

WGS = Whole-genome sequencing. Chr = Chromosome. SHH = Sonic hedgehog. CNS = Central nervous system. NOS = Not otherwise specified.

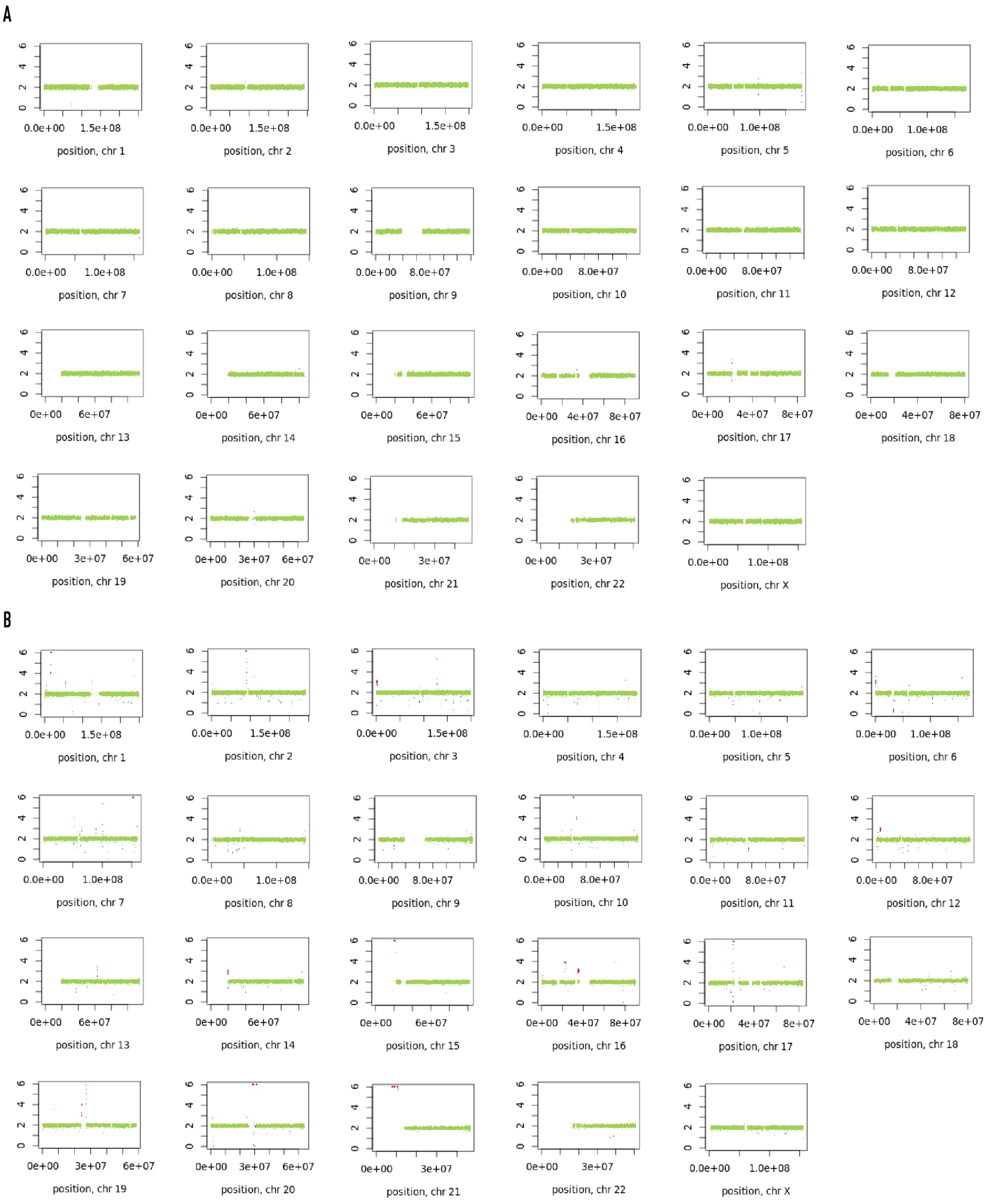


**Fig. S32. Whole-genome sequencing-based copy number profiles for paediatric brain tumour patients 3.**
**(A)** Somatic WGS CNV profile for patient diagnosed with CNS embryonal tumour (grade IV, multi-layered rosettes, NOS) with **(B)** match normal blood.

WGS = Whole-genome sequencing. Chr = Chromosome. SHH = Sonic hedgehog. CNS = Central nervous system. NOS = Not otherwise specified.

**Table S1. Marker genes for factors from patient 1 (organ wide).**
* GEFs representing cancer

| GEF | Marker genes |
|---|---|
| 1 | IGHG2, OGHGP IGHG1, IGHG3, JCHAIN |
| 2 | PCSK7, TAGLN CNN1, MYL9, DES |
| 3 | FOSB, TAGLN, MYL9, DES, PI16 |
| 4 | NEFH, CD177, VEGFA, AOC1, MSMB |
| 5* | SLC4A4, GALNT3, STEAP2, ARMCX5, TTC32, HIF1A |
| 6 | MIR6087, FABP4, HBA2, ADIPOQ, LEP |
| 7 | CFD, PENK, NELL2, HELLPAR, C1QL1 |
| 8 | CHGA, TUBB2A, CALCA, TUBB2B, PRPH |
| 9 | OLFM4, CXCL1, CXCL6, SERPINB3, SAA1 |
| 10 | KLK4, KLK3, KLK2, NEFH, MSMB |
| 11 | MS4A1 (CD20), CD79A, FCER2, VPREB3, TCL1A |
| 12 | ACKR1, RERGL LGI4, MMRN1, ACAN, IGFBP7 |
| 13* | TBC1D3K, TBC1D3H, TBC1D3G, TBC1D3B, TBC1D3C |
| 14* | VGLL3, TMPRSS2, SLC45A3, MME, NCAPD3 |
| 15 | MMP7 ARHGAP40 PIGR TFCP2L1 KRT15 |
| 16* | DDC, NCAPD3, PCOTH, SCGB1D2, SRARP |
| 17* | SOD2, SERPINA3, CHI3L2, SAA2, FAM177B |
| 18 | HSPB1, JUND, NBL1, NPIPB2, CD81 |
| 19 | A2M−AS1, SSTR2, PABPC3, A2M, CCDC152, POSTN |
| 20* | FOLH1B, FOLH1, CRISP3, PPFIA2, ALB, KLK14, DDC, NPY |
| 21 | LTF, SAA2, SAA1, PIGR, CHI3L2 |
| 22 | PTGDS |
| 23* | DNALI1, HIST1H4K, HIST1H4J, CRISP3 CACNA1G |
| 24 | FOLH1B, FOLH1 GSTA2, TRPM8 TGM4, PSCA |
| 25 | FOS, FOSB, EGR1, ZFP36, NR4A1 |

**Table S2. Marker genes for factors from patient 1 (high resolution)**
* GEFs representing cancer

| GEF | Marker genes |
|---|---|
| 1 | PIGR, MMP7, SAA1, CP, CFB |
| 2 | AL078639.1, CCDC80, PI16, LYVE1, OGN |
| 3 | MYH11, CNN1, ACTG2, DES; TPM2 |
| 4 | ACKR1, CCL14, CXorf36, PECAM1, RGS5, IGFBP7 |
| 5 | CD177, NEFH, AOC1, SLC2A5, VEGFA, MSMB |
| 6 | AIF1, LYZ, HLA-DQA1, LAPTM5, CD74 |
| 7* | FABP5, PCA3, PPFIA2, DDC, TFF3 |
| 8 | RIMS1, EHF, HIST1H1E, SLC5A4-AS1, RMRP |
| 9 | KRT15, S100A14, KRT7, KRT5, DEFB1 |
| 10* | TRCG1, OR51E2, TRPM4, HMGCS2, TFF3 |
| 11* | SRARP, PCA3, PHGR1, TRGC1 |
| 12 | NEFH, CD177, LMAN1L, SLC22A3, VEGFA |
| 13 | CCK, NCAPD3, AC016730.1, ANOEO, SLC22A3 |
| 14* | PPFIA2, AC015712.2, CLPTM1L, SAPCD2, TRIR |
| 15* | CITED1, SPON2, KLK12, DDC, PSCA, BANK1 |
| 16 | ANPEP, LMAN1L, ACPP, CPE, KLK3 |
| 17* | NEAT1, FOSB, EGR1, FOS, EGR3 |
| 18 | CCDC152, PHKG1, AC092069.1, SYNC |
| 19* | SNHG8, SNHG19, CSKMT, OST4, ZFAS1 |
| 20* | NPR3, PCAT14, TRGC1, OR51E2, PCA3 |
| 21 | PRR4, CPB1, KCNN4, COL22A1, GGT1 |
| 22* | UBE2C, BIRC5, TOP2A, NUSAP1, CDC20, CDKN3, CRISP3 |
| 23* | PDZK1IP1 (MAP17), FOLH1, RPRML, GDF15 |
| 24* | TRGC1, TMEFF2, PCAT18, ACSM3, PCSK1N, SCGN, GAL, F5 |

**Table S3. Marker genes for factors from patient 2**

| GEF | Top marker genes |
|---|---|
| 1 | PI16, ITGA8, NEXN, IGLC3, JCHAIN |
| 2 | MMP7, PIGR, KRT7, CXCL1, CXCL17 |
| 3 | ACTG2, CNN1, TAGLN, ACTA2, TPM2 |
| 4 | SLC45A3, ACPP, CD177, KLK2, NEFH |
| 5 | TAGLN, PHGR1, LGALS1, MYL9, FLNA |
| 6 | LGALS2, LYZ, HLA-DQA1, LTB, HLA-DQB1, HLA-DRA |
| 7 | KCNQ2, IGF1, PLAC9, PNMT, NELL2 |
| 8 | ACKR1, ESAM, ECSCR, CLEC14A, CDH5 |
| 9 | AGR2, SLC4A4, TSPAN8, PCA3, SERPINB11 |
| 10 | AMACR, PCAT19, SRARP, PCAT14, TRGC1 |
| 11 | AL078639.1, AL355075.4, HIST1H1E, ATRNL1, PAGE4 |
| 12 | NELL2, IGF1, MYLK, LUM, SYNM |
| 13 | EGR1, FOSB, EGR3, NR4A3, FOS |
| 14 | KRT15, WIF1, SLC14A1, KRT23, SMR3B |
| 15 | CCK, AC011043.2, DHRS7, DBI, CPE |
| 16 | ADGRF5, OR51E1, CSRP2, TMEFF2, PPFIA2 |
| 17 | PLEKHH1, MLPH, CPLX3, AC087741.1, FAM13A-AS1 |
| 18 | SNHG19, PRAC1, UQCRH, SNHG8, PCA3 |
| 19 | SPDEF, SPON2, CLDN4, IER2, CLDN3 |
| 20 | AC068389.1, RIMS1, PPDPF, VHL, NKX3-1 |

**Table S4. Top genes for squamous cell carcinoma clusters**

| Cluster | Top Marker genes |
|---|---|
| 0 | COL1A2, COL6A2, C1QA, HLA-DRA, COL1A1 |
| 1 | S100A2, PFN1, MMP1, ATP1B3, SFN |
| 2 | IGLC2, IGHG1, IGKC, IGHG3, IGHG4 |
| 3 | SFN, KRT5, KRT6B, KRT16, KRT6C |
| 4 | MMP1, LAMC2, TNC, TGFBI, S100A2 |
| 5 | KRT1, KRT10, S100A8, KRTDAP, S100A7 |
| 6 | IVL, KRT6B, KRT17, PI3, KRT6C |
| 7 | DCN, COL1A1, SFRP2, COL1A2, IGFBP4 |
| 8 | IGFBP7, A2M, MYL9, SPARC, SPARCL1 |
| 9 | S100A7, S100A8, LGALS7B, KRT2, LY6D |
| 10 | ISG15, CD74, CCL18, HLA-DPB1, TMSB4X |
| 11 | LY6D, RPLP1, LGALS7B, RPS23, RPL41 |
| 12 | DCD, SCGB2A2, MUCL1, LTF, SCGB1D2 |

**Table S5. Top genes for human lymph node clusters**

| Cluster | Top Marker genes |
|---|---|
| 0 | CCL2, SPARCL1, CCL21, GZMB, CCL14 |
| 1 | CXCL13, FDCSP, FCER2, LINC00926, BANK1 |
| 2 | CR2, SERPINE2, MEF2B, RGS13, ELL3 |
| 3 | CXCL9, CCL19, IL7R, TCF7 LEF1 |
| 4 | IGHG2, IGHG1, IGHG4, IGKC, IGHG3 |
| 5 | MYL9, MGP, ACTA2, TAGLN, IGHGP |
| 6 | CCL20, SERPINE1, CHI3L1, CEMIP, CXCL5 |
| 7 | CD209, CLEC4M, SIGLEC1, LYVE1, MARCO |
| 8 | MT-CO1, MT-ND2, MT-ND3, MT-CYB, MT-ATP6 |
| 9 | ISG15, IFI6, MX1, IFI44L, IFIT1 |

**Table S6. Top genes for human glioblastoma multiforme clusters**

| Cluster | Top Marker genes |
|---|---|
| 0 | METTL7B, SLN, CD63, NNMT, CALB1 |
| 1 | VEGFA, SREBF1, MT3, NRN1, GAPDH |
| 2 | HBA2, HBB, HBA1, VEGFA, DKK1 |
| 3 | HSPA1B, PPP1R15A, CCN1, TCIM, SOD2 |
| 4 | SULF1, FABP7, SRPX, COL8A1, TMOD1 |
| 5 | TAC1, VGF, NNAT, STMN2, TRH |
| 6 | APOD, DCN, PRELP, VWF, IGKC |
| 7 | APOD, IGKC, ISLR, DCN, CLDN5 |
| 8 | NNMT, APOC1, HLA-DRA, IGKC, CD74 |
| 9 | COL4A1, COL4A2, FN1, COL3A1, BGN |
| 10 | GPR37L1, ALDOC, SLC6A1, KCNJ16, FXYD1 |
| 11 | FABP5, SLN, S100B, PTN, DBI |
| 12 | FOS, ADM, HILPDA, CEBPD, DDIT4 |
| 13 | TM4SF1, SDC1, ALDH1A3, GJB2, PLAUR |

**Table S7. Top genes for breast cancer (invasive ductal carcinoma) clusters**

| Cluster | Top Marker genes |
|---|---|
| 0 | IGLC2, IGHG3, IGKC, IGLC3, IGHG1 |
| 1 | MALAT1, AL627171.2, SAMHD1, C1QA, CCDC80 |
| 2 | CXCL14, TTLL12, GFRA1, DEGS1, ARMT1 |
| 3 | LINC00052, COX6C, SNCG, WFDC2, SLC39A6 |
| 4 | CRISP3, SLITRK6, C6orf141, VTCN1, SERHL2 |
| 5 | CPB1, FCGR3B, LINC02224, KLHDC7B, SCGB1D2 |
| 6 | ALB, MGP, ZNF350-AS1, S100G, SERPINA3 |
| 7 | ACKR1, AQP1, IGFBP7, VWF, MALAT1 |
| 8 | AC087379.2, KRT37, S100G, SCGB2A2, PGM5-AS1 |
| 9 | LINC00645, SLC30A8, MUC5B, COLEC12, PVALB |
| 10 | MRPS30-DT, PDE5A, WFDC2, RBM20, MRPS30 |
| 11 | S100A9, S100A8, MGP, IER3, LTF |

**Table S8. Top genes for medulloblastoma clusters**

| Cluster | Top Marker genes |
|---|---|
| 1 | MALAT1, MT-ND2, MT-ATP6MT-ND4, MT-CYB |
| 2 | KRT31, CXCR4, RPS27A, EEF1B2, RPS7 |

**Table S9. Squamous cell carcinoma clone cell annotation**
Percentage of spots for each clone that harbour "Stroma", "Tumour Epithelia" or "Non-Invasive Epithelia".

| Clone | Stroma | Tumour Epithelia | Non-Invasive Epithelia | Total (should = 100 when complete) | Notes |
|---|---|---|---|---|---|
| A | 15 | 80 | 5 | 100 | |
| B | 96 | 2 | 2 | 100 | |
| C | 2 | 0 | 98 | 100 | |
| D | 98 | 2 | 0 | 100 | About 90% of the stromal spots are lymphocytes |

**Table S10. Breast cancer clone cell annotation**

Percentage of spots for each clone that harbour “Stroma”, “Tumour Epithelia”, “DCIS”, “Lymphatic cells”

| Clone | Stroma | Tumour Epithelia | DCIS | Normal Epithelia | lymphatic cells | Total | Notes | Annotation Certainty |
|---|---|---|---|---|---|---|---|---|
| A | 90 | 10 | 0 | 0 | 0 | | Stroma, periphery of clone B (invasive Ca), possibly TILs | |
| B | 0 | >95% | 0 | 0 | 0 | | Invasive Cancer, possible lower grade than the other invasive | Uncertain |
| C | 0 | >95% | 0 | 0 | 0 | | Invasive Ductal Carcinoma | |
| D | 0 | >95% | 0 | 0 | 0 | | Invasive Ductal Carcinoma | |
| E | 0 | >95% | 0 | 0 | 0 | | Invasive Ductal Carcinoma | |
| F | 15 | 85% | 0 | 0 | 0 | | Mixed, Stroma + Invasive Carcinoma | |
| G | 0 | >95% | 0 | 0 | 0 | | Invasive Ductal Carcinoma | |
| H | 0 | 10% | 90% | 0 | 0 | | DCIS | Certain |
| I | >95% | 0 | 0 | 0 | 0 | | Stroma | |
| J | >95% | 0 | 0 | 0 | 0 | | Stroma | |
| K | >95% | 0 | 0 | 0 | 0 | | Stroma | |
| L | 50% | 0 | 0 | 0 | ≥50% | | Definite immune cells in upper right corner | |

**Data D1. Infer CNV HMM annotations for clone trees**

Tables and images showing annotations (HMM and manual) used for clone trees.

**Data D2. Synthetic data analysis (separate zip file)**

Files related to method evaluation with synthetic data, such as: design file, the generated synthetic data, results from spatial inferCNV when applied to the synthetic data, and script for evaluation of results.
