## Supplementary Data D2 for "The spatial landscape of clonal somatic mutations in benign and malignant tissue": evaluate.R

```
library(pheatmap)
library(dendextend)
library(RColorBrewer)
library(dplyr)

# define input paths
res.dir <- "../results"
synth.dir <- "../sythetic_data"
spatial.dir <- file.path(synth.dir,"spatial_data")
single.cell.dir <- file.path(synth.dir,"single_cell_data")

genome.pth <- file.path(synth.dir,"genome.tsv")

ann.pth <- file.path(spatial.dir,"st-annotation.tsv")
cell.by.spt.pth <- file.path(spatial.dir,"cell_by_spot.txt")

sc.gen.pth <-  file.path(single.cell.dir,"sc-genome_profile.tsv")
tree.pth <- file.path(res.dir,"infercnv.observations_dendrogram.txt")
pred.state.pth <- file.path(res.dir,"17_HMM_predHMMi6.hmm_mode-samples.pred_cnv_genes.dat")


# define output path
out.dir <- "../analysis"


# read data
tree <- ape::read.tree(tree.pth)
sc.gen.data <- read.table(sc.gen.pth,sep = "\t",header = 1,row.names = 1)
genome.data <- read.table(genome.pth,sep = "\t",header = 1,row.names = 1)
ann.data <- read.table(ann.pth,sep = "\t",header = 1,row.names = 1) 
pred.state <- read.table(pred.state.pth,header = 1)

# -- synthetic data 

# get spots found in each clone
groups <- list()
for (i in 1:length(tree)){
  groups[[i]] <- tree[[i]]$tip.label
}

# get all spots in vector, reverse order to match inferCNV heatmap
all.spots <- rev(unlist(groups))
n.spots <- length(all.spots)

# get clone annotations for each spot
annotations <- data.frame(matrix(0,n.spots,1))
rownames(annotations) <- all.spots
colnames(annotations) <- "Clone"

for (g in 1:length(groups)){
  for (s in groups[[g]]){
    annotations[s,"Clone"] <- g
  }
}
# convert to factor, to allow column coloring
annotations[,"Clone"] <- as.factor(annotations$Clone)

# read cell-by-spot file
cbs <- read.delim(cell.by.spt.pth,
                  header = F,
                  sep = " ",
                  row.names = 1,
                  as.is = T)

# format cell-by-spot matrix

# list of cells that reside on each spot
cell.on.spot <- list()
# vector to hold clone identity of each cell, derived from spot
cell.cluster <- c()
for (s in rownames(annotations)){
  # register cells on spot s
  cell.on.spot[[s]] <- as.numeric(strsplit(cbs[s,'V3'],',')[[1]])
  # number of cells at spot s
  n.cells <- length(cell.on.spot[[s]])
  # get spot s clone id
  clu.id <- as.character(annotations[s,"Clone"])
  # add cluster id for all cells to vector
  cell.cluster <- c(cell.cluster,rep(clu.id,n.cells))
}
# get names of cells on selected spots
sel.cells <- paste0("Cell_",unlist(cell.on.spot))
# grab genomic profile of selected cells
cell.genome <- sc.gen.data[sel.cells,]

# convert single cell data annotation to data frame
sc.annotations <- as.data.frame(cell.cluster)
# add row and colnames to single cell nnotation
rownames(sc.annotations) <- rownames(cell.genome)
colnames(sc.annotations) <- "Clone"
# convert annotation to factor, for plotting
sc.annotations[,"Clone"] <- as.factor(sc.annotations$Clone)

# set annotation colors, selected to agree with inferCNV
anno.colors <- list(Clone = c("4"="#FFED6F",
                              "3"= "#F0D1E1",
                              "2"="#A9A0B2",
                              "1"="#8DD3C7"))


png(file.path(out.dir,"sc-genome-profile-true.png"),
    width=1000,height=500)

pheatmap(cell.genome,
         cluster_rows = F,
         cluster_cols = F,
         color = c("blue","#FAF5F5","red","darkred"),
         annotation_row = sc.annotations,
         show_colnames = F,
         show_rownames =  F,
         annotation_colors = anno.colors,
         )

dev.off()

# -- inferred results

# add column indicating clone identity
pred.state$clone <- sapply(pred.state$cell_group_name,
                           function(x){strsplit(as.character(x),"\\.")[[1]][1]})

# mapping between clone and the cluster identities used in synth data
clone.to.clust <- c("4" ="Clone_D",
                    "3" = "Clone_C",
                    "2" = "Clone_B",
                    "1" = "Clone_A")

# mapping between inferCNV state and amplification/deletion
state.to.cnv <- c("1"=0,"2"=0,"3"=1,"4"=2,"5"=2,"6"=2)
# convert inferCNV state to amp/del
pred.state$state <- state.to.cnv[pred.state$state]

# generate "dummy" genome based on the inferred state
pred.cell.genome <- matrix(1,nrow(cell.genome),ncol(cell.genome))
rownames(pred.cell.genome) <- rownames(cell.genome)
colnames(pred.cell.genome) <- colnames(cell.genome)

# iterate over cells
for (s in rownames(pred.cell.genome)){
  # get clone id of the given cell
  clone.id <- clone.to.clust[as.character(sc.annotations[s,"Clone"])]
  # make sure clone is in predicted file
  if (clone.id %in% pred.state$clone){
    # subset with predictions w.r.t. clone of cell s
    sub.pred <- pred.state[pred.state$clone == clone.id,]
    # update genome with predicted gene alterations for clone
    pred.cell.genome[s,as.character(sub.pred$gene)] <- sub.pred$state
  }
}

# save inferred state visualization
png(file.path(out.dir,"sc-genome-profile-inferred.png"),
    width=1000,height=500)

pheatmap(pred.cell.genome,
         cluster_rows = F,
         cluster_cols = F,
         color = c("blue","#FAF5F5","red"),
         annotation_row = sc.annotations,
         show_colnames = F,
         show_rownames =  F,
         annotation_colors = anno.colors,
         border_color = "black",
         )

dev.off()

# -- Evaluate Performance

# get clones present in GT data
clones <- unique(sc.annotations$Clone)
# create matrix to store GT clonotype states
true.genome <- data.frame(matrix(0,length(clones),ncol(pred.cell.genome)))
rownames(true.genome) <- paste0("Clone_",clones)
colnames(true.genome) <- colnames(pred.cell.genome)

# get mean profile for cells in clone
for (clone in clones){
  true.genome[paste0("Clone_",clone),] <- round(colMeans(cell.genome[sc.annotations$Clone == clone,]))
}

# convert to categorical values
is.amp <- true.genome > 1
is.del <- true.genome < 1
is.none <- true.genome == 1

true.genome[is.amp] = "A"
true.genome[is.del] = "D"
true.genome[is.none] = "N"

# get clones present in inferred data and representative cell id
rep.cells <- unique(sc.annotations)
# create matrix to store inferred clonotype states
pred.genome <- data.frame(matrix(0,length(clones),ncol(pred.cell.genome)))
rownames(pred.genome) <- paste0("Clone_",clones)
colnames(pred.genome) <- colnames(pred.cell.genome)

# get profile for representative cell
for (clone in clones){
  cell.id <- rownames(rep.cells)[rep.cells$Clone == clone]
  pred.genome[paste0("Clone_",clone),] <- pred.cell.genome[cell.id,]
}

# convert to categorical values
is.amp <- pred.genome > 1
is.del <- pred.genome < 1
is.none <- pred.genome == 1

pred.genome[is.amp] = "A"
pred.genome[is.del] = "D"
pred.genome[is.none] = "N"

# compute accuracy mean and std across clones
acc <- rowSums(pred.genome == true.genome) / ncol(pred.genome)
acc.mean<- mean(acc)
acc.std <- sd(acc)

# prepare data frame to save accuracy results
eval.res <- c(acc.mean,acc.std)
names(eval.res) <- c("mean","std")
eval.res <- data.frame(eval.res)
# save accuracy results
write.table(eval.res,file.path(out.dir,"total-eval-res.tsv"),sep = "\t",quote = F)

# creat TPR/TNR table
res.summary <- data.frame(matrix(0,2,4))
rownames(res.summary) <- c("A","D")
colnames(res.summary) <- c("TPR","FPR","TNR","FNR")

n.tot <- nrow(true.genome) * ncol(true.genome)
for (x in c("A","D")){
  p.X <- pred.genome == x
  n.p.X <- sum(p.X)
  res.summary[x,"TPR"] <- sum(pred.genome[p.X] == true.genome[p.X]) / n.p.X
  res.summary[x,"FPR"] <- 1 - res.summary[x,"TPR"]
  res.summary[x,"TNR"] <- sum(pred.genome[!p.X] == true.genome[!p.X]) / (n.tot - n.p.X)
  res.summary[x,"FNR"] <- 1 - res.summary[x,"TNR"]
}

write.table(res.summary,file.path(out.dir,"score-res.tsv"),sep = "\t",quote = F)
```
