## Supplementary Data D2 for "The spatial landscape of clonal somatic mutations in benign and malignant tissue": README

# Content of this supplementary data file is:

├── code
│   └── evaluate.R # code used to evaluate results
├── README
├── results # output from spatial InferCNV, documented on associated web-page
│   ├── 17_HMM_predHMMi6.hmm_mode-samples.genes_used.dat
│   ├── 17_HMM_predHMMi6.hmm_mode-samples.pred_cnv_genes.dat
│   ├── 17_HMM_predHMMi6.hmm_mode-samples.pred_cnv_regions.dat
│   ├── CloneSelection_21052021.pdf
│   ├── infercnv.17_HMM_predHMMi6.hmm_mode-samples.png
│   ├── infercnv.21_denoised.png
│   ├── infercnv.observations_dendrogram.txt
│   └── syntheticdata_clones.tsv
└── sythetic_data
    ├── design.toml
    ├── single_cell_data
    │   ├── sc-expression.tsv # single cell expression matrix
    │   ├── sc-genome_profile.tsv # single cell genome profile
    │   └── sc-meta_data.tsv # single cell meta data
    └── spatial_data
        ├── cell_by_spot.txt # id of cells residing at each spot
        ├── st-annotation.tsv # annotation file (benign/aberrant)
        ├── st-expression.tsv # spatial transcriptomics expression data
        ├── st-genome_profile.tsv # spatial transcriptomcs genome (average per spot) profile
        └── st-meta.tsv # spatial transcriptmics meta data
