## Supplementary Data D2 for "The spatial landscape of clonal somatic mutations in benign and malignant tissue": design.toml

```
[genome]
n_genes = 500
genome_size = 1e5
event_spread = 10
concentration = 1
initial_event_size = 15


[population]
n_initial_cells = 2
domain_side_size = 200
fraction_drop = 0.05
n_clusters = 10
max_cells = 5e3

    [population.transitions]
    s0s1 = 0.2
    s1s0 = 0.2

    [population.rates_state_0]
    move_rate = 0.4
    stay_rate = 0.1
    spawn_rate = 0.4
    death_rate  = 0
    mutation_rate = 0.0
    [population.rates_state_1]
    move_rate = 0.3
    stay_rate = 0.45
    spawn_rate = 0.2
    death_rate  = 0
    mutation_rate = 0.8


    [[population.initial_coordinates]]
    x = 50
    y = 50

    [[population.initial_coordinates]]
    x = 150
    y = 150


[spatial]
depth = 5000
n_spots_x = 20
n_spots_y = 20
spot_radius = 2.5
```
