## Supplementary figures and images for "The spatial landscape of clonal somatic mutations in benign and malignant tissue"

### CloneSelection_21052021.pdf

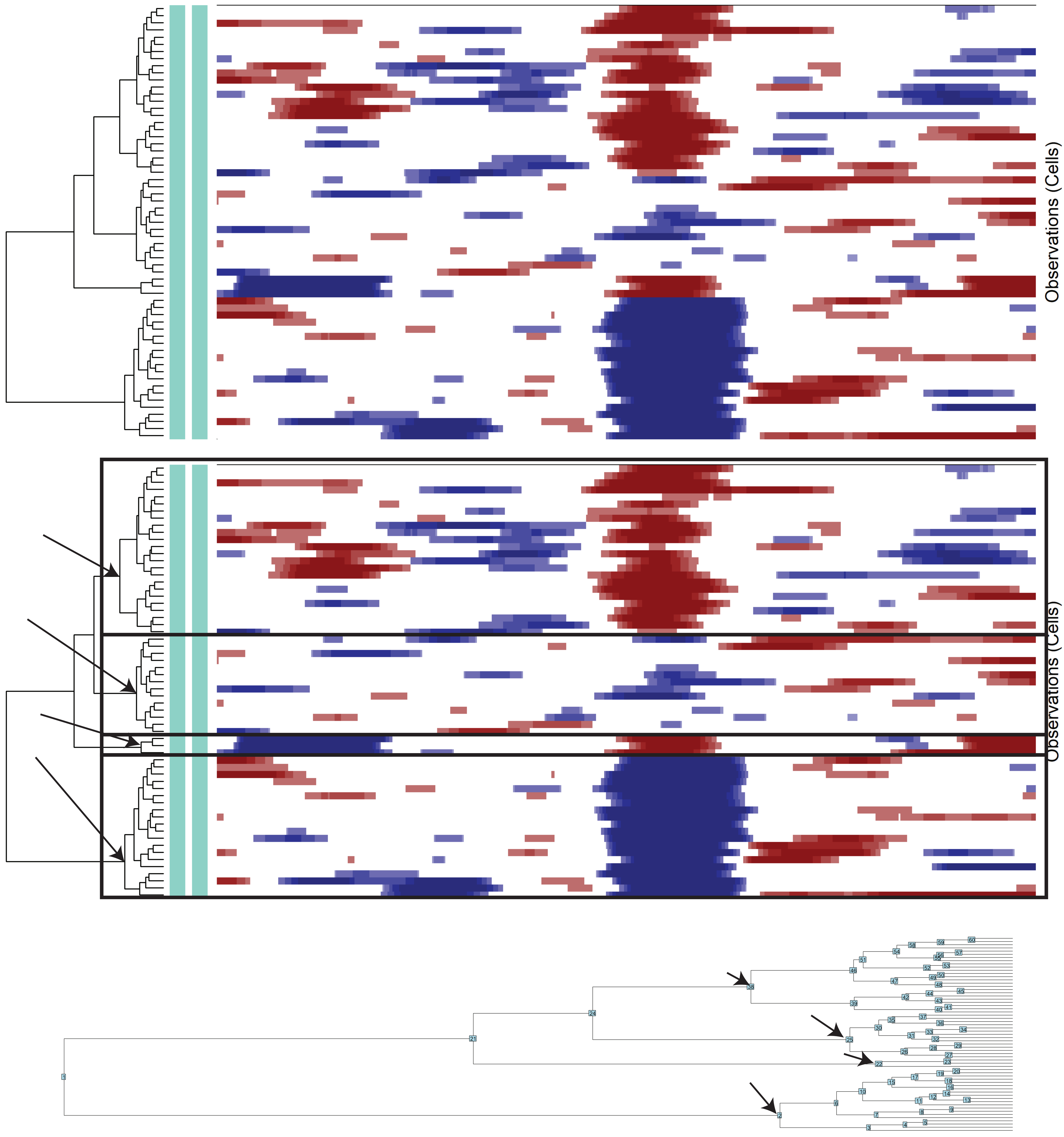


**17_HMM_preds**

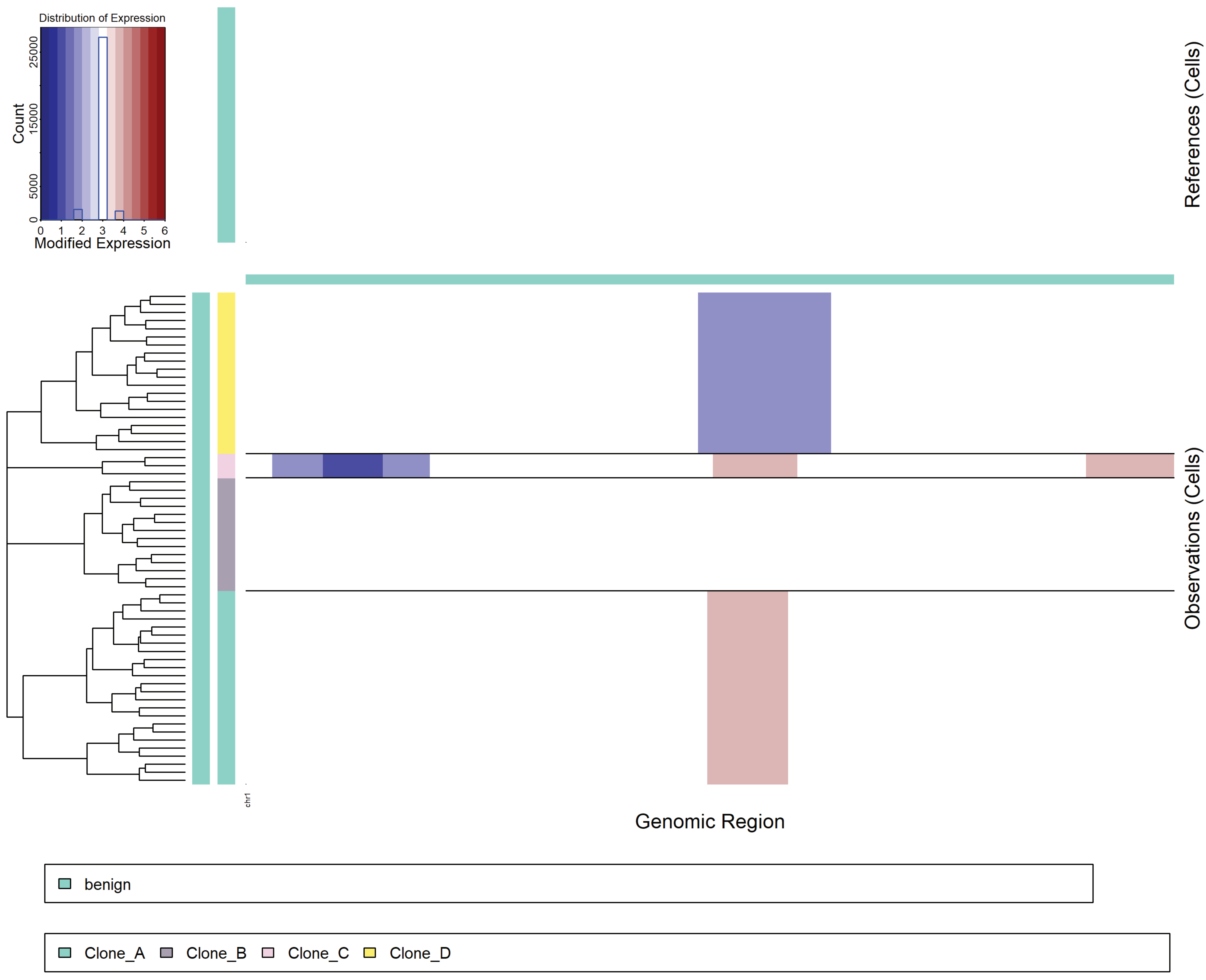


**21_denoised**

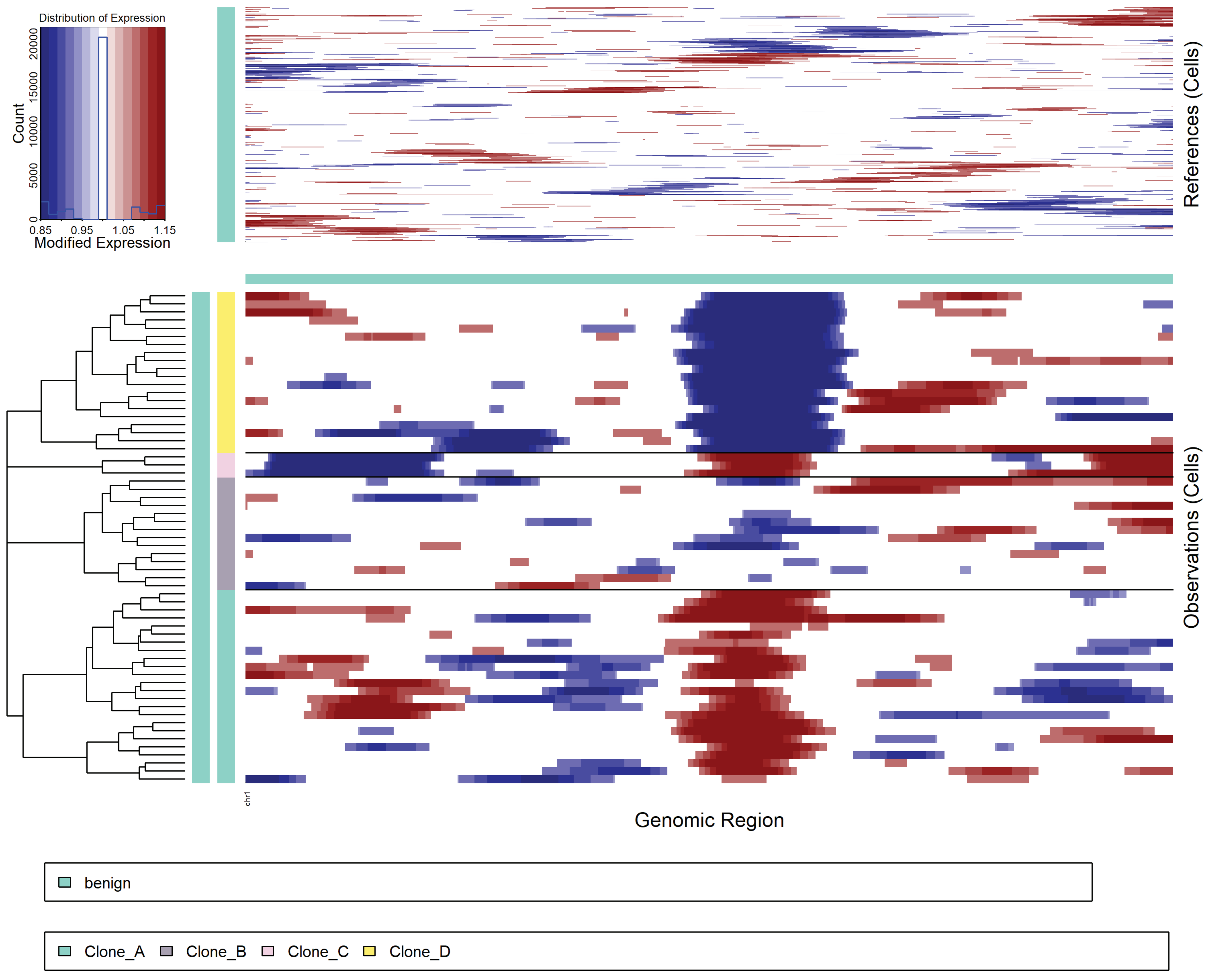

### infercnv.17_HMM_predHMMi6.hmm_mode-samples.png

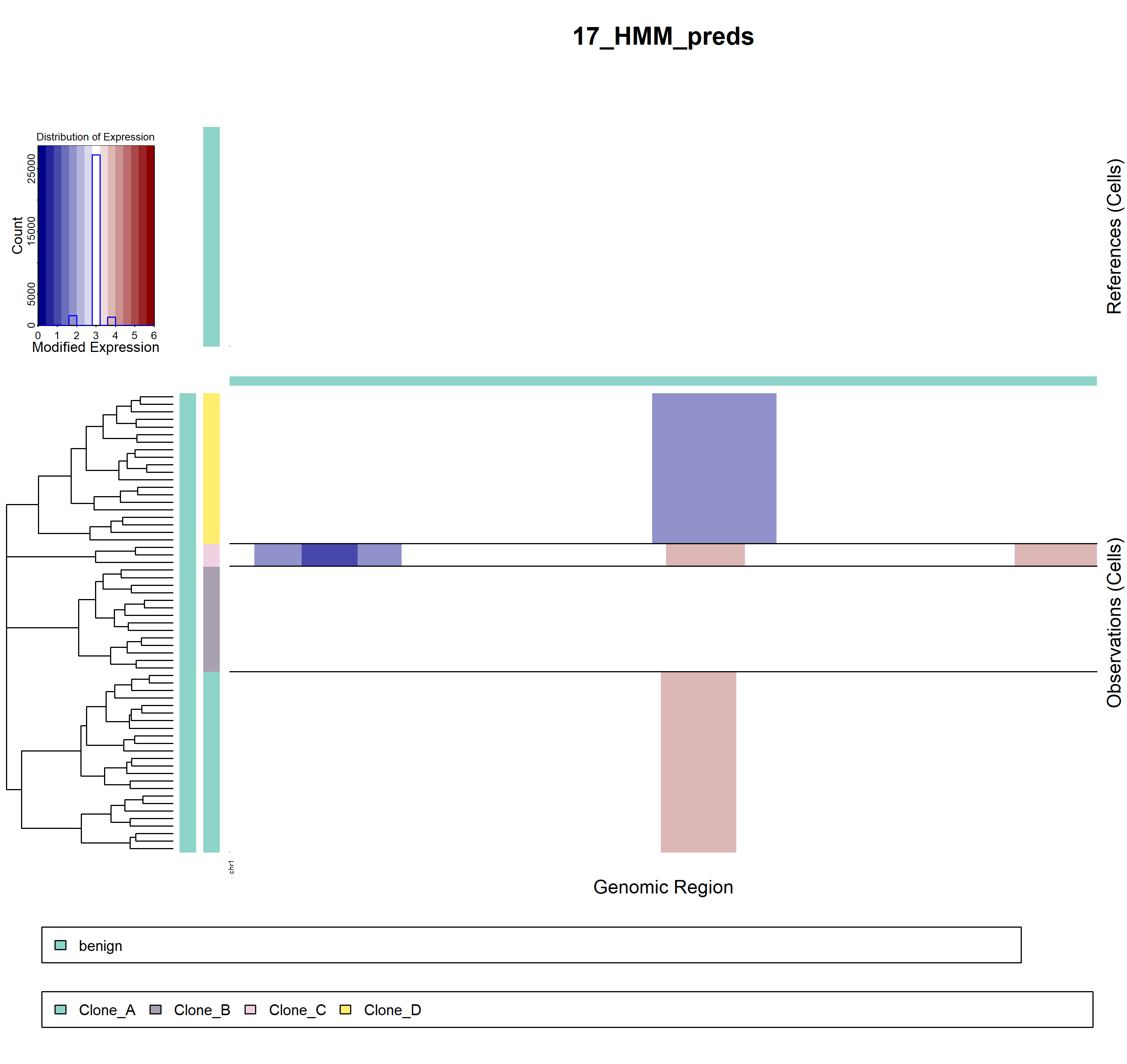

### infercnv.21_denoised.png

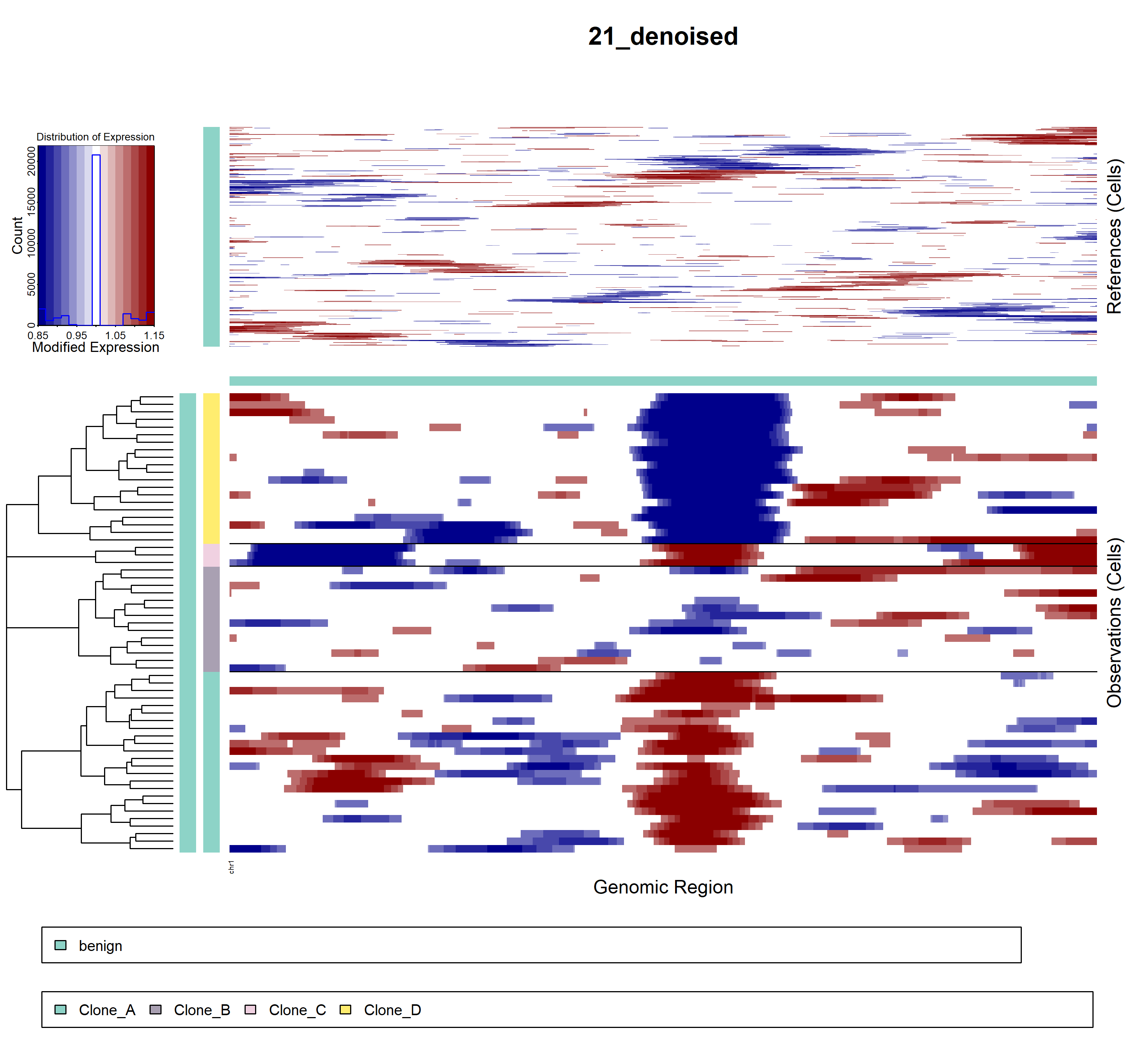
